# One-stop analysis of DIA proteomics data using MSFragger-DIA and FragPipe computational platform

**DOI:** 10.1101/2022.10.28.514272

**Authors:** Fengchao Yu, Guo Ci Teo, Andy T. Kong, Ginny Xiaohe Li, Vadim Demichev, Alexey I. Nesvizhskii

## Abstract

Liquid chromatography (LC) coupled with data-independent acquisition (DIA) mass spectrometry (MS) has been increasingly used in quantitative proteomics studies. Here, we present a fast and sensitive approach for direct peptide identification from DIA data, MSFragger-DIA, which leverages the unmatched speed of the fragment ion indexing-based search engine MSFragger. MSFragger-DIA conducts a database search of the DIA tandem mass (MS/MS) spectra prior to spectral feature detection and peak tracing across the LC dimension. We have integrated MSFragger-DIA into the FragPipe computational platform for seamless support of peptide identification and spectral library building from DIA, data dependent acquisition (DDA), or both data types combined. We compared MSFragger-DIA with other DIA tools, such as DIA-Umpire based workflow in FragPipe, Spectronaut, and *in silico* library-based DIA-NN and MaxDIA. We demonstrated the fast and sensitive performance of MSFragger-DIA across a variety of sample types and data acquisition schemes, including single-cell proteomics, phosphoproteomics, and large-scale tumor proteome profiling studies.

## Introduction

Liquid chromatography (LC) coupled to data independent-independent acquisition (DIA) mass spectrometry (MS) strategy has emerged as a widely used technological platform for quantitative protein profiling, especially in large-scale studies [1, 2]. Compared to the data dependent acquisition (DDA) MS strategy, in which a selected peptide ion population is isolated and subjected to tandem MS (MS/MS) fragmentation, the fragment ion information in DIA is acquired on all peptide ions within a certain window of m/z values, sequentially covering the entire relevant range (e.g., 400-1200 Da). This helps alleviate the stochastic nature of peptide identification in DDA-based strategies, and DIA has been shown to produce more complete (i.e., less missing values) peptide and protein quantification matrices across multiple samples [2]. DIA has been successfully used in a variety of proteomics applications, including post-translational modification (PTM) analysis [3-6], protein-protein interaction [7, 8], immunopeptidomics [9-11], and heritability analysis [12].

MS data used as part of a DIA analysis workflow can be categorized into primary and auxiliary data. Auxiliary MS data is used solely to build a spectral library for subsequent targeted extraction of quantification for each peptide ion in the library from the primary DIA data. Examples of such auxiliary, “library-only” MS data include data acquired from pooled samples (e.g., all individual samples profiled in the study) fractionated using offline LC and analyzed using DDA [2], fractionated in the gas phase, and analyzed using narrow-window DIA (GPF-DIA) [13, 14]. For low-input proteomics, including single-cell proteomics, library-only data can be acquired in DDA or DIA mode from samples prepared from higher amounts of starting material [15-17]. The primary DIA data are DIA runs acquired on individual study samples, typically without fractionation, although the use of fractionation has been explored [18]. The primary DIA data are used for extracting peptide ion quantification, but they may also be used for building the spectral library, either alone or in combination with auxiliary DDA or DIA data [19].

The computational analysis of DIA data has two major components:1) creation of the target spectral library, a collection of peptide ions that are targets for the subsequent quantification step, along with their LC retention times (RT), as well as m/z values and intensities of the corresponding fragment ions; 2) extraction of quantification from the primary DIA data for all peptide ions in the target spectral library. Quantification is performed using targeted extraction tools, such as Skyline [20], OpenSWATH [21], EncyclopeDIA [13], Spectronaut [22], and DIA-NN [23], with additional intensity data normalization and peptide-to-protein rollup [24, 25]. The targeted quantification step depends on the quality of the input spectral library [26]. An ideal library would be as experiment specific as possible, that is, it would be complete (containing all peptide ions that are likely to be detectable in the analyzed samples) and as specific (i.e., it would not contain unrelated peptides) as possible [27, 28]. Library information, such as the RT and fragment ion intensities, should match the actual DIA data being analyzed well. Thus, building spectral libraries via the direct identification of peptides from primary DIA data [19] has emerged as a widely used computational data analysis strategy.

Direct identification of peptides from DIA data was first proposed and implemented in DIA-Umpire [19], along with the concept of building combined (hybrid) spectral libraries from primary DIA and auxiliary (e.g., DDA) data. The “spectrum-centric” identification approach of DIA-Umpire is based on feature detection in MS1 and MS/MS data, followed by grouping precursor peptide and fragment ion signals exhibiting similar LC elution profile to generate the so-called pseudo-MS/MS spectra. These pseudo-MS/MS spectra can be searched using tools developed for conventional DDA data, such as MSFragger [29], X! Tandem [30], or Comet [31], followed by peptide-spectrum match (PSM) validation with PeptideProphet [32] or Percolator [33], protein inference with ProteinProphet [34], and target-decoy based false discovery (FDR) filtering [35]. The DIA-Umpire strategy has been increasingly adopted in other tools and pipelines, including Spectronaut’s directDIA. A drawback of this strategy is that the untargeted signal extraction of the precursor and fragment ion peak curves from DIA MS1 and MS/MS scans can be time-consuming. The sensitivity of the peptide identification process may also be suboptimal compared to targeted peptide detection methods. Other strategies for the direct identification of peptides from DIA data have also emerged, often taking the peptide-centric perspective [36] to the same problem, as exemplified by tools such as PECAN [37] (Walnut in EncyclopeDIA [13]). Direct search of DIA MS/MS spectra against repository-wide spectral libraries or sequence databases has also been explored [38, 39]. The ability to predict proteome-wide spectral libraries using deep learning [40-46], followed by the creation of more refined, experiment-specific libraries for targeted extraction, has also been shown to be very useful [47], and formed the basis for the *in silico* library-based (also known as “library-free”) mode of DIA-NN [23, 48], EncyclopeDIA [13], and MaxDIA [49]. However, only peptides with common modifications (such as oxidation, acetylation, and phosphorylation) can typically be predicted using the current *in silico* spectral library software. Furthermore, workflows based on proteome-wide spectral library prediction suffer from additional limitations, such as long prediction times or requirements for additional graphics processing units (GPUs).

We have developed a new approach for direct peptide identification from DIA data, leveraging the unmatched search speed of fragment ion indexing [29]. It is based on conducting a database search of DIA MS/MS spectra prior to feature detection or peak tracing, blurring the difference between the analysis of DIA and DDA MS/MS spectra. It has been implemented as MSFragger-DIA, a separate module in the MSFragger search engine [29, 50, 51], and publicly available since MSFragger version 3.1 (released on September 30, 2020). Using MSFragger software, one can identify peptides from either DDA or DIA data, or from both data types combined, allowing the seamless generation of a hybrid spectral library for the most sensitive analysis. We compared MSFragger-DIA with other tools, such as DIA-Umpire-based workflow in FragPipe, Spectronaut, DIA-NN *in silico* library-based, and MaxDIA. We demonstrated the fast and sensitive performance of MSFragger-DIA across a variety of sample types and acquisition schemes, including single-cell proteomics and phosphoproteomics applications. MSFragger-DIA has been fully integrated into the FragPipe computational platform (https://fragpipe.nesvilab.org/). In tandem with the spectral library building module EasyPQP [48] and the quantification tool DIA-NN, MSFragger-DIA in FragPipe enables the complete analysis of DIA data, from peptide identification to peptide and protein quantification.

## Results

### MSFragger-DIA algorithm

An overview of the algorithm and its implementation within the FragPipe computational platform is shown **in Figure 1**. MSFragger-DIA starts with a direct search of MS/MS spectra against the entire protein sequence database, before any peak tracing or feature detection procedures (**Figure 1a**). MSFragger-DIA starts with deisotoping [50] and extraction of the isolation window information from the spectra. Because no precursor charge information is available for DIA MS/MS spectra, it enumerates all predefined charge states to calculate the lower and upper precursor mass bounds for each spectrum. Each spectrum is then searched against all database peptides within the precursor mass range. To calibrate fragment masses, MSFragger-DIA performs two searches: a fast calibration search and a full search. The first, more restricted search (which considers peptides with precursor charge states 2 and 3 only) is used to select high-quality spectra to build a mass error profile and perform mass calibration, as described previously [51]. MSFragger-DIA then performs a full search using the calibrated data. The outcome of this step is a list of peptide candidates (by default, 20 best-scoring peptides per spectrum in wide window DIA data; 12 in narrow window GFP-DIA data), which is then further refined as described below.

**Figure 1.**
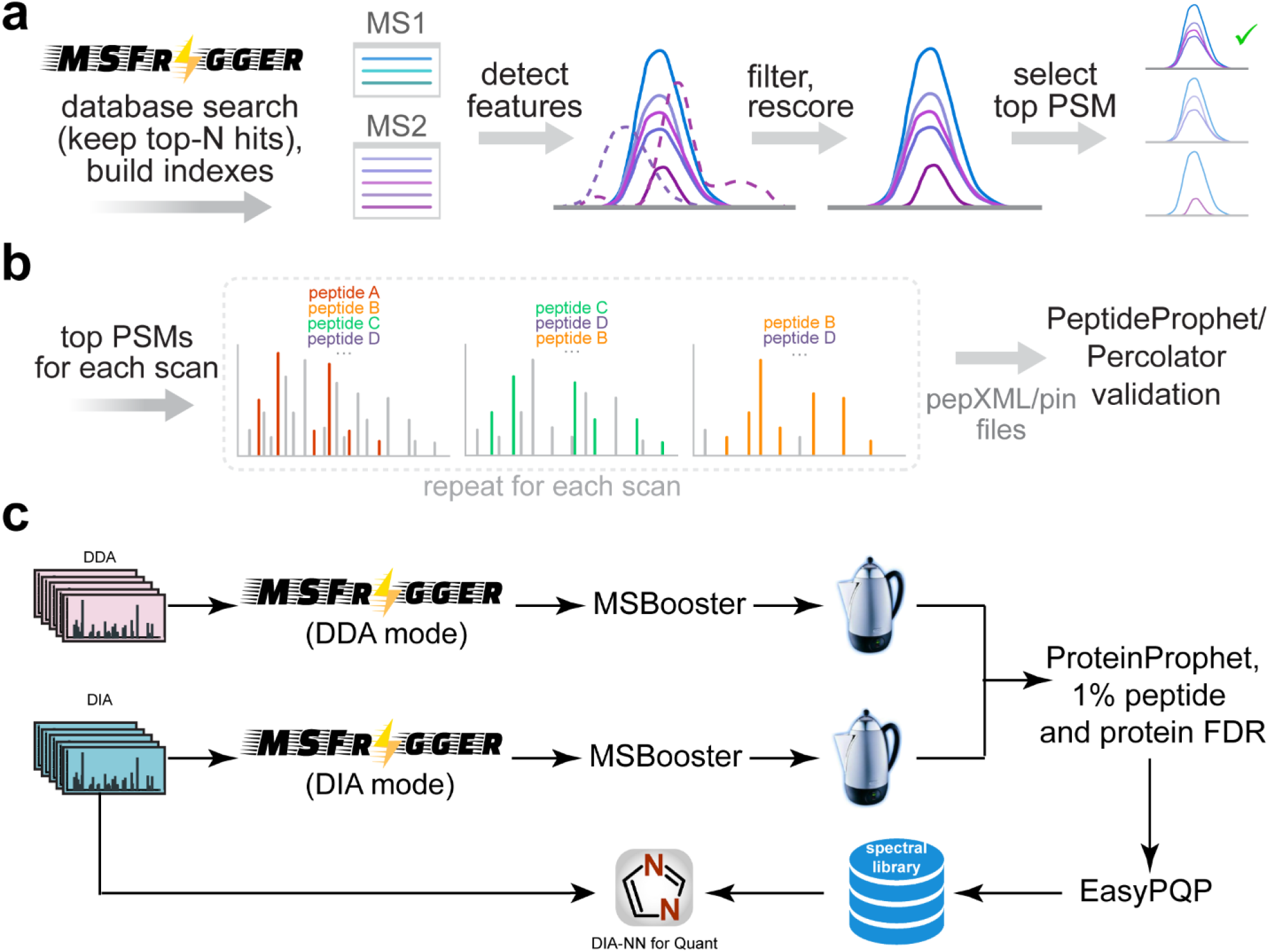
Overview of MSFragger-DIA in FragPipe. **(a)** DIA spectra are searched by MSFragger-DIA directly using precursor candidates determined from the isolation window. MSFragger-DIA builds MS1 and MS/MS spectral indexes, which are then used to detect extracted ion chromatogram (XIC) features for all fragment and precursor peaks in a peptide-spectrum match (PSM). Noisy fragment peaks are filtered out based on the XIC, PSMs are rescored, and only the top scoring PSM from each feature is kept. **(b)** Within each DIA MS/MS scan, a greedy method is used to remove matched peaks and iteratively rescore peptide candidates from the top-N list. Finally, MSFragger-DIA generates pepXML and pin files for PeptideProphet and Percolator to estimate the peptide probability in FragPipe. **(c)** Hybrid (combined DIA and DDA) data analysis workflow in FragPipe (“FP-MSF hybrid” in the main text). Both DDA and DIA data are used to build a combined spectral library. This spectral library is used to quantify peptides from the DIA data using DIA-NN.

In the second step, MSFragger-DIA traces peaks, extracts ion chromatograms, and detects features of all candidate peptides for each spectrum determined as described above. This can be done quickly using the MS1 and MS/MS spectral indexing technology that we described previously [52]. For each peptide, MSFragger-DIA refines the list of matched fragments that can contribute to the score. The apex LC retention times of the fragments and precursor features are collected, and the median retention time is calculated. Fragments with retention times that differ from the median by more than a certain value (0.1 minutes by default) are filtered out. If a precursor ion is detected in the MS1 data and has an apex retention time outside the allowable range, the PSM is filtered out. Note that if no precursor ion feature is detected in the MS1 spectrum, MSFragger-DIA still retains the corresponding PSM, as there are cases where fragment ion signals exist despite missing (undetectable) precursor features. MSFragger-DIA also filters out PSMs with aberrant isotope distributions [53]. After peak tracing and peptide and fragment filtering, MSFragger-DIA normalizes and rescores peptide matches using hyperscore [29, 30].

Since each MS/MS spectrum is matched against candidate peptides independently, the same experimental peaks may match and contribute to the score of multiple different peptides in the candidate peptide list; a greedy method is used to prevent this from happening (**Figure 1b**). Given a spectrum and a list of candidate peptides, MSFragger-DIA selects the highest-scoring peptide and removes matched fragment peaks from the spectrum. Then, it normalizes the remaining peaks and rescores the peptides on the candidate list to obtain the second-highest scoring PSM. The matched peaks are again removed, peptides are rescored, and the process continues until there are fewer than four peaks left in the spectrum, or no peptide candidates remained. Finally, MSFragger-DIA generates output files compatible with PeptideProphet and Percolator for rescoring and FDR estimation.

MSFragger-DIA has been fully integrated into FragPipe, allowing one-stop DIA data analysis (**Figure 1c and Supplementary Figure 1**). In FragPipe, the output of MSFragger-DIA and MSFragger can be processed by MSBooster [54] to leverage additional scores using deep-learning prediction. The output from MSBooster is fed to Percolator for additional rescoring and posterior error probability calculation, followed by ProteinProphet [34] for protein inference, Philosopher [55] for false discovery rate (FDR) filtering, and spectral library building with EasyPQP [48]. With FragPipe, users can build the libraries from DDA (with MSFragger), DIA (with MSFragger-DIA), or all data combined (hybrid library). The resulting library is then used to extract peptide ion quantification from the primary DIA data using DIA-NN, which is available as a part of FragPipe.

### Evaluating false discovery rates (FDR) using a benchmark dataset

To benchmark the MSFragger-DIA algorithm and evaluate the actual false discovery rates of the entire workflow, we used the dataset from Frohlich et al. [56], consisting of four experiments: “Lymphnode”, “1-25”, “1-12”, and “1-06”. The samples used in the first experiment contained peptides from *H. sapiens* only. The other three experiments have a mixture of peptides from *H. sapiens* and *E. coli* (mixed with the indicated *H. sapiens* to *E. coli* ratios). There are 92 wide window DIA runs (primary DIA runs), and auxiliary MS files used for spectral library building: six narrow window GPF-DIA runs, and 20 DDA runs. We refer to this as “benchmark” dataset. We performed two MSFragger-DIA based analyses (**Figure 2**): (1) building the spectral library using the primary DIA data plus the narrow window GPF-DIA data (FP-MSF workflow, see **Methods**); (2) building the spectral library from all available data, that is, also including the DDA data (FP-MSF hybrid workflow). DIA-NN, which is available as part of FragPipe, was used to quantify the precursors from the DIA runs using these spectral libraries. The Spectronaut and DIA-NN *in silico* library-based (labelled “DIA-NN” in the Figures) results were used for comparison. The number of *E. coli* precursors detected in the “Lymphnode” experiment (that contained *H. sapiens* proteins only) was used to estimate the actual FDR (see **Methods)**.

**Figure 2.**
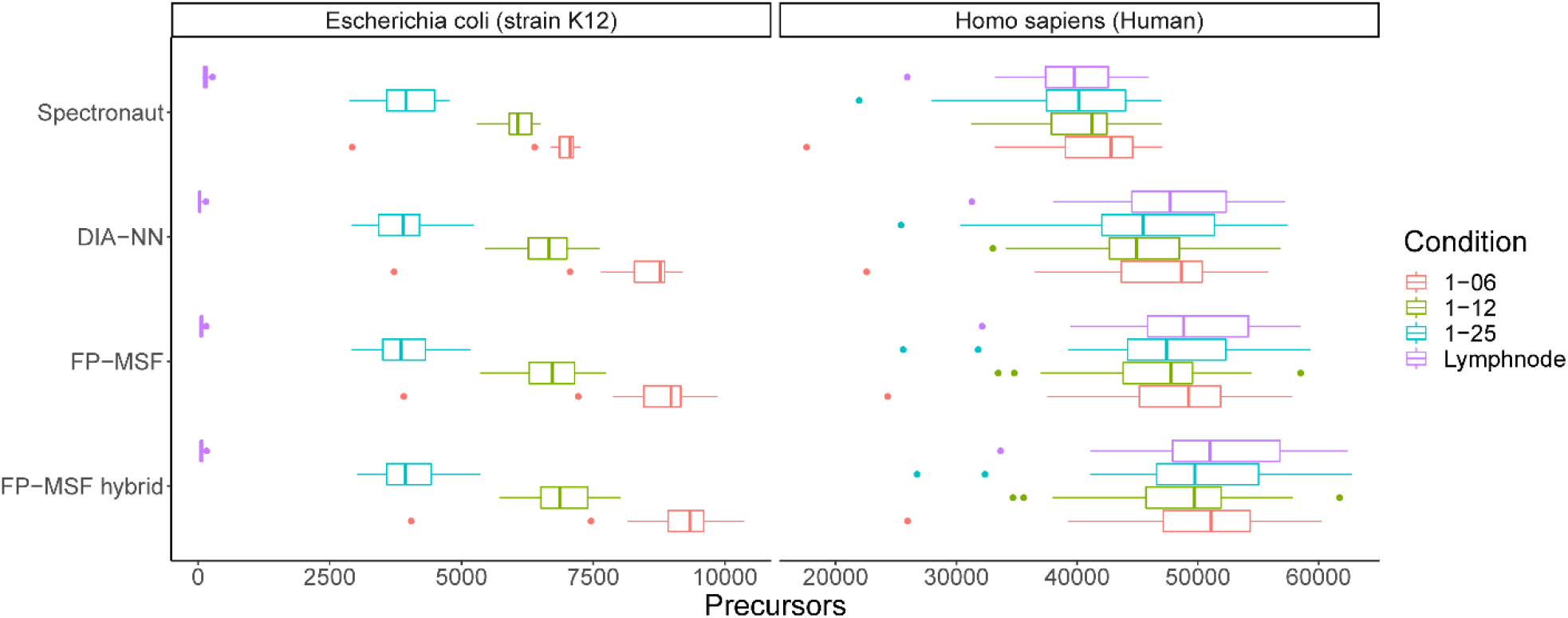
Boxplots of the number of quantified peptide ions in the benchmark dataset. There are four experiments with four different colors. The “Lymphnode” experiment has peptides from *H. sapiens* only. The other three experiments have peptides from both *H. sapiens* and *E. coli*. The peptides from *E. coli* and *H. sapiens* are shown in two separate panels. *E-coli* peptides identified in the “Lympnode” experiment are considered false identifications.

We also investigated replacing, when performing targeted quantification in DIA-NN, empirically observed fragments and their intensities in the FragPipe/MSFragger-DIA generated spectral libraries with the *in silico* predicted values. Note that the list of peptide ions included in the library and their retention time values remained unchanged. Replacing the experimental fragment peaks with the predicted peaks resulted in a lower actual FDR (**Supplementary Table S2**). Thus, this option was enabled for all analyses reported in this work.

**Figure 2** shows the boxplots of the number of precursors quantified from the DIA runs, plotted separately for *H. sapiens* and *E. coli* precursors, for each of the four experiments. The *E. coli* boxplots for the Lymphnode experiment show that all tools control the false positives well (all *E. coli* peptide detections in the “Lymphnode” experiment are assumed to be false), with slightly higher number of false identifications observed for Spectronaut (**Supplementary Table S2)**. The boxplots from the other three experiments, for both *H. sapiens* and *E. coli* precursors, show that the MSFragger-DIA based workflows resulted in the most quantified precursors. Including DDA data in the spectral library building step (FP-MSF hybrid workflow) provided an additional small boost in the number of quantified precursors.

### Staggered-windows DIA with gas phase fractionation (GFP)

We then tested the ability of MSFragger-DIA to identify peptides from DIA data acquired using the “staggered” windows approach [57, 58], with additional narrow window GFP-DIA data acquired for spectral library building. The first dataset, taken from Searle et al. [13], contains six narrow window GPF-DIA runs plus three wide window, single-injection DIA runs (the primary DIA runs for quantification) from a HeLa lysate, 90 minutes gradient time. The isolation windows of the GPF-DIA and single-injection DIA runs are 4 Th and 24 Th, respectively. After demultiplexing the staggered windows using ProteoWizard [59], the effective isolation windows were halved. We refer to this dataset as “2018-HeLa”. The second, similar dataset was taken from Searle et al. [60] and contains six GPF-DIA runs, and four single-injection DIA runs of *S. cerevisiae* lysate,115 minutes gradient time. The isolation windows are 4 Th and 8 Th, respectively, with effective widths halved after demultiplexing. We refer to this dataset as “2020-Yeast”. More information regarding these data can be found in **Supplementary Table 1**.

We used the FP-MSF workflow to analyze these two datasets (in each dataset, GFP-DIA and primary DIA data were processed together using MSFragger-DIA). We also compared the results with that from the original publication [60], and with the results of running DIA-NN *in silico* library and MaxDIA pipelines (see **Methods**). **Figure 3a** and **3c** show the numbers of quantified peptides from single-injection DIA runs. Although the 2018-HeLa dataset was published earlier [13], Searle et al. [60] re-analyzed the same dataset using an optimized spectral library. Thus, we included the results from Searle et al. [60] in the Figure (labeled “Searle et al. 2020”). Bar heights indicate the average number of quantified peptides, and circles show the number of quantified peptides in each individual run. MaxDIA gave unreasonably low numbers of quantified peptides in these data, and thus was excluded from plotting (see **Supplementary Figure 2a and 2c**). FP-MSF demonstrated the highest sensitivity among the compared tools. **Figure 3b** and **3d** show the distributions of coefficient of variation (CV) of the quantified peptides from the single-injection DIA runs. No CVs data were available from Searle et al. [60]. DIA-NN *in silico* library and FP-MSF (which also uses DIA-NN, but for quantification using only the FP-MSF generated library) yielded similarly low CVs, whereas MaxDIA yielded significantly higher CVs (**Supplementary Figure 2b and 2d**). Overall, this analysis shows that MSFragger-DIA works well with GPF-DIA and staggered window DIA data, exhibiting the highest sensitivity compared with other pipelines.

**Figure 3.**
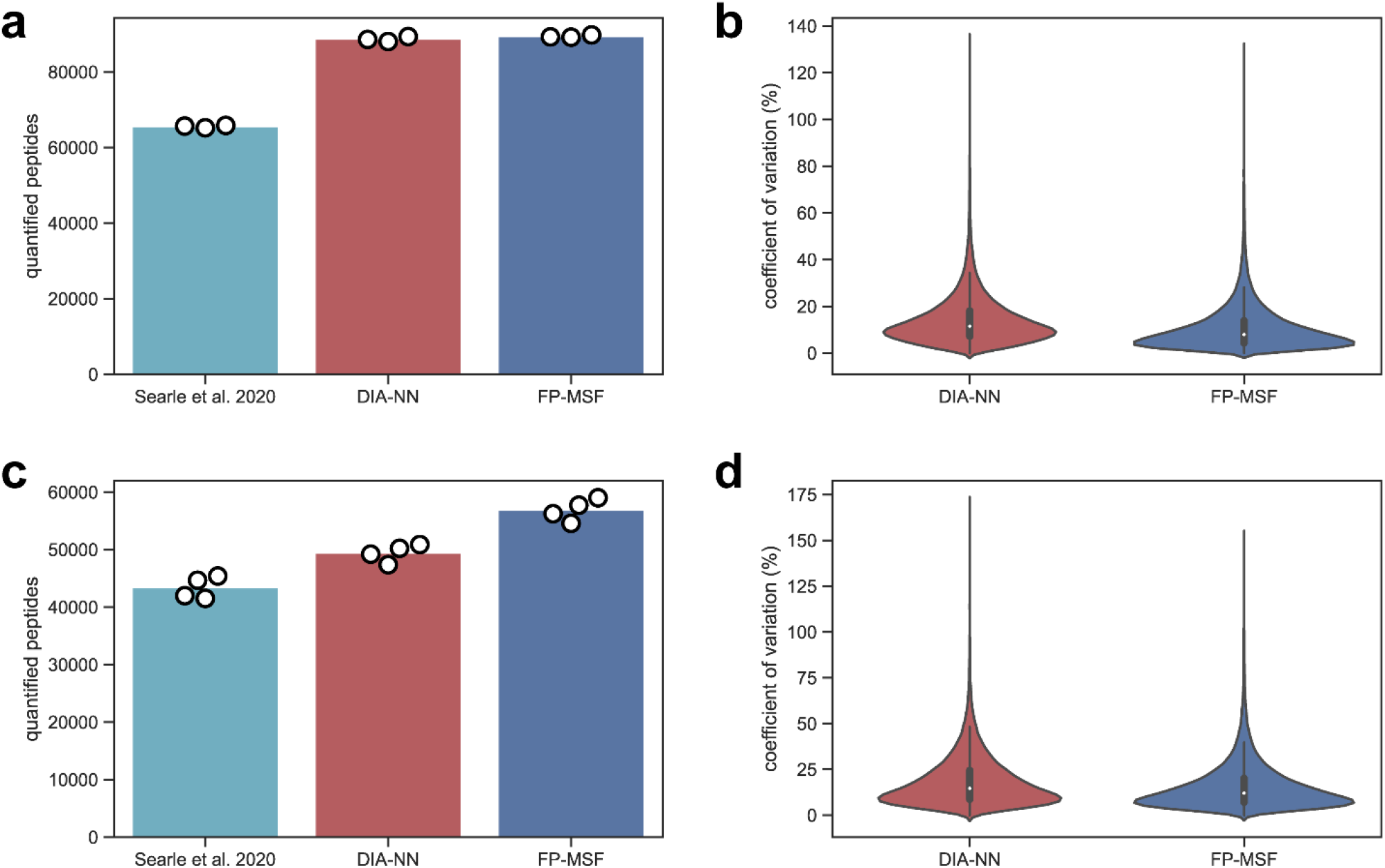
Quantified peptides and coefficient of variation (CV) from the 2018-HeLa and the 2020-Yeast datasets using DIA-NN *in silico* library-based and FP-MSF pipelines. **(a)** Bar plots of the quantified peptides from 2018-HeLa dataset. The bar height is the average number of three replicates. The white dots indicate the numbers from individual replicates. The results from the original publication (obtained using EncyclopeDIA) are also shown. **(b)** Violin plots of peptide CVs from 2018-HeLa dataset. **(c)** Same as (a) for 2020-Yeast dataset. **(d)** Same as (b) for 2020-Yeast dataset.

### Low input and single-cell proteomics data

We then used a dataset published by Siyal et al. [16] to demonstrate the performance of MSFragger-DIA in analyzing low-input cell data. We selected two experiments containing 0.75 ng and 7.5 ng of starting material. Each experiment was performed in three replicates. A detailed list of these files is provided in **Supplementary Table 1**. We refer to this dataset as “low-input-cell”. The authors also generated DIA data on samples with higher amounts of starting material, 1.5 ng and 1 μg, for the purpose of building spectral libraries. To compare with the published result, we used the FP-MSF pipeline to perform two analyses. The first analysis used the spectral files from the 0.75 ng samples (primary DIA data for quantification) and the 1.5 ng samples (DIA data used for spectral library building only). The second analysis used the spectral files from the 7.5 ng and 1 μg samples. We also used FP-DIAU, DIA-NN *in silico* library, and MaxDIA pipelines to analyze the same data for comparison (see **Methods**). **Figure 4a** and **4b** show the number of quantified proteins from 0.75 ng and 7.5 ng samples, respectively. All the tools were run using the same set of input DIA data. We used the MaxLFQ approach [61] for peptide-protein intensity roll-up in all workflows that used DIA-NN quantification (see **Methods**). However, we used the “Intensity” columns for MaxDIA because of a very high rate of missing quantification values with the “LFQ Intensity” columns (MaxLFQ approach, **Supplementary figure S3a and S3b**). The figures show that FP-MSF and DIA-NN *in silico* library-based pipelines have similar sensitivities that are higher than those of the other tools.

**Figure 4.**
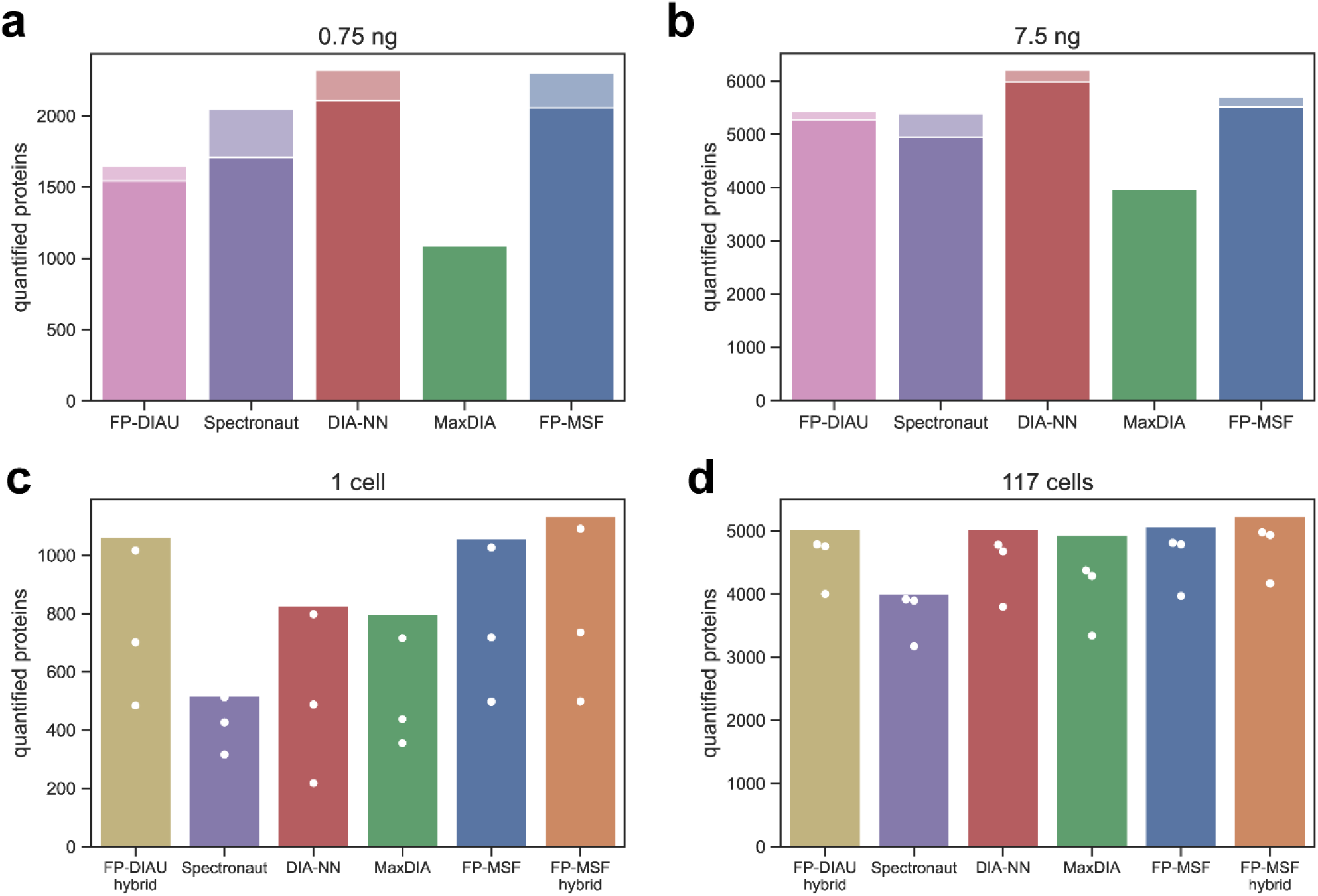
The number of quantified proteins from the low-input-cell and the single-cell datasets. **(a-b)** Bar plots from analyzing the **low-input-cell** dataset with 0.75ng and 7.5ng of starting material. Proteins with missing values (zero intensities) were discarded. The dark color is from the proteins with CVs less than 20%, and the light color is from the proteins with CVs greater than or equal to 20%. **(c-d)** Same as above, for the **single-cell** dataset, for 1-cell and for 117 cell data.

We also used the single-cell dataset published by Gebreyesus et al. [15]; we refer to this dataset as “single-cell”. We picked two experiments with 1 cell and 117 cells. Each experiment has three replicates. The authors also generated two sets of DIA runs to build two spectral libraries, referred to as “small-sample-lib” and “large-sample-lib” (with the latter generated using a higher amount of starting material, see **Methods and Supplementary Table 1)**. There are 9 DIA runs and 3 DDA runs in the small-sample-lib list of files. There are 4 DIA runs and 4 DDA runs used for large-sample-lib. We used FP-DIAU hybrid, MaxDIA, DIA-NN *in silico* library, FP-MSF, and FP-MSF hybrid (using DDA data in addition to the library-only DIA data) pipelines to analyze this dataset (see **Methods**). **Figure 4c** and **4d** show the number of quantified proteins from 1 cell and 117 cells experiments, respectively. As for the low-input-cell dataset, for MaxDIA we used the “Intensity” instead of “LFQ Intensity” because a significant fraction of the proteins had zero MaxLFQ intensity (**Supplementary figure S3c and S3d**). The results show that FP-MSF hybrid pipeline quantified more proteins than the other tools in the 1-cell experiment. In the 117-cell experiment, the difference between different pipelines was less noticeable, except for Spectronaut (based on the results provided by the authors in the publication).

### Phosphoproteomics data

After showing that MSFragger-DIA performs well in global proteome data, including single-cell data, we used a phosphopeptide-enriched dataset [62] (**Supplementary Table 1)** to evaluate the performance of MSFragger-DIA in identifying phosphorylated peptides. The dataset contains two single-injection replicates for six different melanoma cell lines. We refer to this dataset as the “melanoma-phospho” dataset. The data were acquired with variable-width isolation windows over a 120 minute gradient. Because no DDA data were generated in this experiment, we used FP-DIAU, DIA-NN *in silico* library-based, and FP-MSF pipelines (see **Methods** for details). Proteins with more than 50% of missing values were discarded. Although all pipelines produced similar numbers of quantified phosphopeptides (**Figure 5a**), this dataset highlights the speed advantage of MSFragger-DIA. We used a Windows desktop and a Linux server (see **Methods**) to benchmark the computational times (**Figure 5b and 5c)**. The total runtime was broken down into different steps. For FP-DIAU workflow there are pseudo-MS/MS extraction by DIA-Umpire, database search using MSFragger in the DDA mode, rescoring, FDR filtering, and DIA-NN quantification steps. For DIA-NN *in silico* library, there are *in silico* spectral library prediction, identification, and quantification steps. For the FP-MSF pipeline there are database searching using MSFragger-DIA, rescoring and FDR filtering, and DIA-NN quantification steps. FragPipe with MSFragger-DIA had the fastest overall speed; it is at least six times faster than that of the DIA-NN *in silico* library-based workflow for these data.

**Figure 5.**
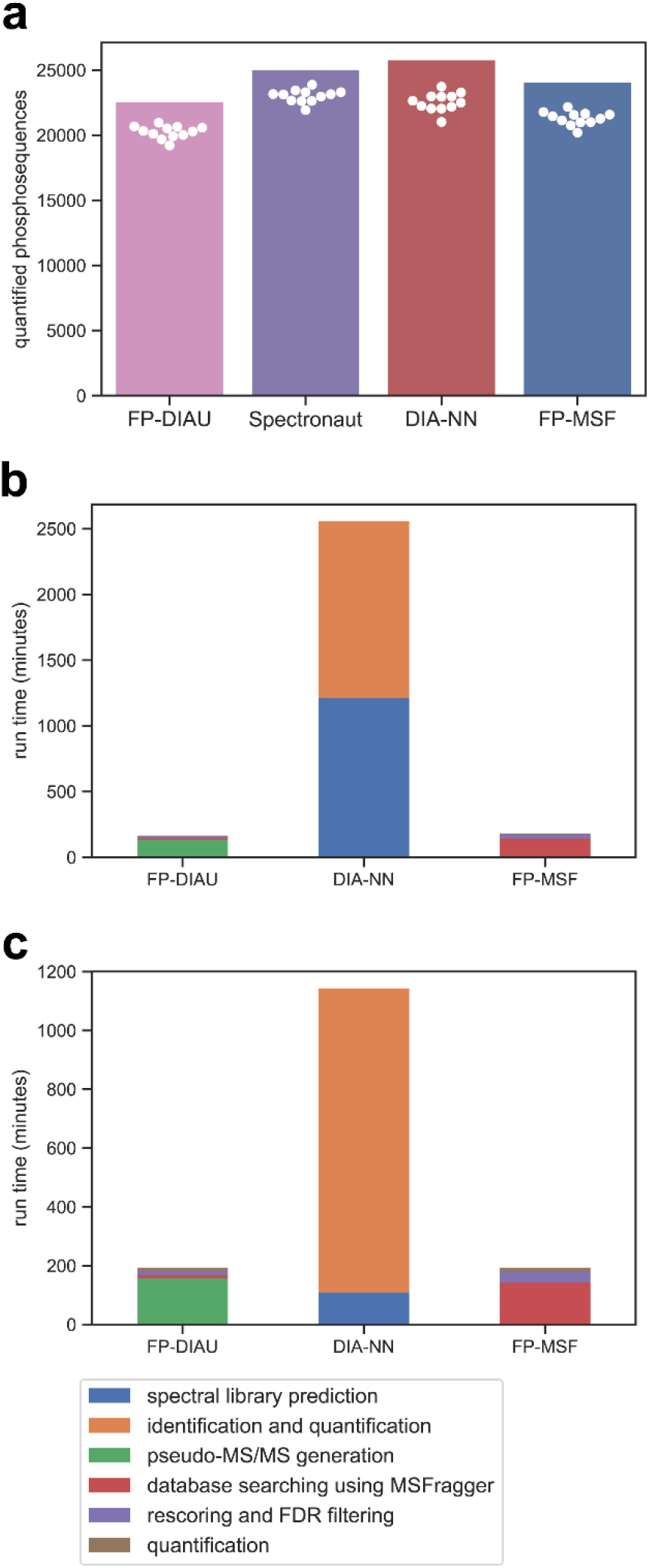
The number of quantified phosphopeptide sequences and runtime for the melanoma-phospho dataset. **(a)** Bar plots of the number of quantified phosphopeptide sequences. The bar height is the total number, and the white dots are the numbers of sequences quantified in individual runs. **(b)** The runtime analysis performed on a Windows desktop. **(c)** The runtime analysis performed on a Linux server.

### Large-scale DIA based quantification study

We used a clear cell renal cell carcinoma (ccRCC) cohort [63] from a recent Clinical Proteomic Tumor Analysis Consortium (CPTAC) project to demonstrate large-scale DIA analysis with MSFragger-DIA. This dataset contains 187 single-injection DIA runs from normal and tumor samples, plus 8 fractionated DDA runs from pooled samples, all acquired with a 140 minute LC gradient. Detailed file lists are provided in **Supplementary Table 1**. We used the FP-DIAU, FP-DIAU hybrid, DIA-NN *in silico* library, FP-MSF, and FP-MSF hybrid pipelines to analyze the data (see **Methods). Figure 6a** shows the numbers of proteins (counting unique genes after protein to gene mapping) quantified from the single-injection DIA runs. Proteins with more than 50% of missing values were discarded. Without using DDA data, all three pipelines (FP-DIAU, FP-MSF, and DIA-NN *in silico* library) performed similarly, quantifying ∼7000 proteins. Adding DDA data as part of the spectral library building step resulted in a small increase in the identification numbers (FP-DIAU hybrid and FP-MSF hybrid), again confirming the utility of using auxiliary data, such as fractionated DDA data, when available. Running FP-MSF hybrid pipeline took less than 20 hours on the Linux server, and MSFragger-DIA based analyses had the fastest run time. Because it was impractical to time the entire dataset of 187 files uninterruptedly, a more detailed comparison between the pipelines was carried out using a subset of 20 DIA runs. The jobs were run on the same standard Windows desktop (**Figure 6b**) and the same Linux server (**Figure 6c)**, as mentioned in the previous section (see **Methods**). The time spent on predicting the *in silico* spectral library for MaxDIA was not included because a previously generated library was used. FP-MSF pipeline (including the quantification using DIA-NN) was at least two times faster than DIA-NN *in silico* library-based analysis, and five times (Windows desktop) or ten times (Linux server) faster than MaxDIA. Overall, these results show that FragPipe with MSFragger-DIA for peptide identification directly from DIA data can be used to process large-scale datasets, even with relatively standard desktop hardware. In contrast, other pipelines for DIA analysis are likely to require the use of cloud computing pipelines [64], which can be harder for a typical user to install and deploy.

**Figure 6.**
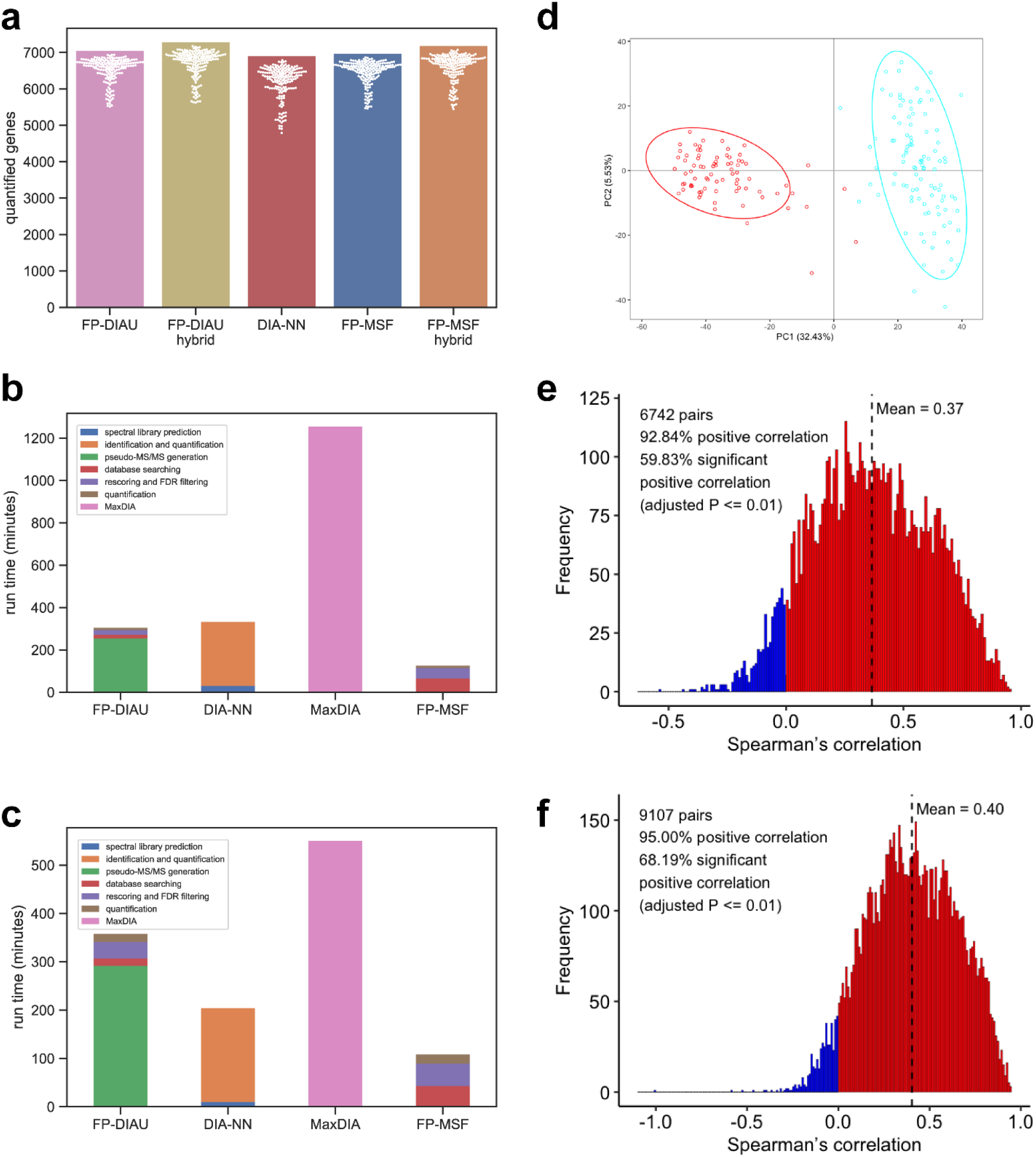
**(a)** Bar plots of the number of quantified proteins (unique gene symbols) in the ccRCC dataset. The bar height is the total number. The white dots are the numbers from individual runs. **(b)** The runtime of analyzing 20 runs of the **ccRCC** dataset on a Windows desktop. **(c)** The runtime of analyzing 20 runs on a Linux server. **(d)** PCA plot based on the FP-MSF hybrid results, showing tumor (blue) and normal (red) samples. **(e)** Histogram of Spearman’s correlation coefficients between the RNA-Seq and the DIA protein abundance data (FP-MSF hybrid pipeline). **(f)** Histogram of the Spearman’s correlation coefficients between the RNA-Seq and the TMT DDA-based protein abundance data.

Given the availability of tandem mass tag (TMT) DDA proteomics data – the main quantitative proteomics platform used by the CPTAC [65]–and RNA-seq data for the same ccRCC patients [63], we performed additional analyses across these data types. We used OmicsEV [66], a recently described tool for quality control and comparative evaluation of omics data tables. We used as input, in addition to the RNA-seq (taken from [63]) and TMT-based quantification data (mzML files from [63], reprocessed as part of this work using the default FragPipe TMT workflow), DIA protein quantification tables from FP-DIAU, FP-DIAU hybrid, FP-MSF, FP-MSF hybrid, and DIA-NN *in silico* library-based pipelines. OmicsEV produces comprehensive visual and quantitative plots that help evaluate data quality of individual data tables and facilitate the comparisons (for the full output from OmicsEV, see **Supplementary Data 2**). Reassuringly, all quantitative proteomics data tables produced similar results across all OmicsEV metrics. **Figure 6d** shows the principal component analysis (PCA) plot for the quantification tables from the FP-MSF hybrid pipeline. The tumor and normal samples are well separated, as observed in the original study, based on the TMT DDA data. OmicsEV also calculated the gene-wise correlation between the RNA and protein abundances, with Spearman’s correlation for the FP-MSF hybrid pipeline results shown in **Figure 6e**. For comparison, the gene-wise correlation between the RNA and TMT-based protein abundance data is shown in **Figure 6f**. DIA and TMT DDA-based protein quantification data have similar gene-wise correlations with RNA abundances, albeit slightly higher for the TMT DDA-based data. Furthermore, more proteins were quantified using TMT DDA. This is not unexpected, however, given that the TMT DDA data was generated on highly fractionated peptide samples (25 fractions), whereas DIA data was acquired without fractionation. The functional pathways identified as enriched in tumor vs. normal samples were nevertheless similar between the DIA and TMT based data (**Supplementary Data 2**). Overall, our analysis shows that DIA data is comparable to that from the more established, TMT-based protein quantification platform.

## Discussion

We described a new method for identification of peptides directly from DIA data. The MSFragger-DIA strategy differs from that of DIA-Umpire by reversing the order in which the two key steps are performed: detection and correlation of precursor and fragment ion features, and peptide-spectrum matching. While DIA-Umpire starts with untargeted feature detection, MSFragger-DIA first determines a list of all meritorious candidate peptides for each MS/MS scan without spectral deconvolution. Feature detection and peak tracing across the LC dimension are performed in MSFragger-DIA as a second step, in a targeted manner, and for selected candidate peptide ions. Using several common experimental DIA workflows, we demonstrated that MSFragger-DIA is faster than DIA-Umpire followed by the conventional (DDA mode) MSFragger search. It also has a higher sensitivity for peptide identification, in part due to improved feature detection and peak tracing performed in MSFragger-DIA in the targeted mode. However, we consider the MSFragger-DIA and DIA-Umpire workflows to be complementary. One advantage of the DIA-Umpire is that extraction of pseudo-MS/MS spectra needs to be done once; these spectra can then be re-searched using MSFragger (in DDA mode) multiple times, for example, using different sequence databases or search parameters. In contrast, MSFragger-DIA needs to be run each time, starting from the raw data. Furthermore, the speed advantage of MSFragger over DIA-Umpire is less significant for searches containing multiple variable modifications. For nonspecific searches (e.g., HLA peptidome data), MSFragger-DIA is significantly slower; thus, the default FragPipe workflow for nonspecific DIA searches is based on DIA-Umpire.

MSFragger-DIA, with its direct database search-first approach, essentially blurs the difference between the analysis of DIA and DDA MS/MS spectra. Thus, we believe that MSFragger-DIA can be further extended to the analysis of wide-window DDA data [67, 68], enabling identification of chimeric (co-fragmented) peptides from such data. The strategy described here is also different from those based on peptide-centric searches using *in silico* predicted spectral libraries. Using MSFragger-DIA, it is not necessary to predict the entire spectral library in advance, and there is no restriction on the size of peptides, peptide ion charge states, or modifications that can be considered. However, our workflows in FragPipe (for both DDA and DIA data) do benefit from deep learning-based predictions at the subsequent, rescoring stage with the help of MSBooster and Percolator [54]. Interestingly, we observed some advantages of using predicted spectra (instead of empirical ones) at the final targeted quantification extraction step. This indicates that while MSFragger-DIA has high sensitivity for peptide identification, some fragment ions may not be detectable in every MS/MS scan where the peptide was confidently identified. Because the spectral library building step in EasyPQP considers each MS/MS scan in isolation, some fragments that are detectable using targeted peak tracing (in FragPipe, using DIA-NN) may be missing in the empirical spectra in the library. Replacing the empirical spectra with the predicted spectra improves the number of quantified precursors. Such a replacement, however, is not advised in all situations, for example, when performing MSFragger-DIA searches with less common variable modifications, because such spectra may not be predicted well.

Recently, Bruker developed a timsTOF platform to couple ion mobility separation with time-of-flight (TOF) mass spectrometer. To generate DIA data using the timsTOF platform, researchers proposed diaPASEF data acquisition strategy [69], and demonstrated a very promising performance of this platform. diaPASEF data poses new challenges to data analysis, as it becomes necessary to accommodate an additional dimension of ion mobility separation. There are also more spectra to process because of the high speed of the TOF mass spectrometer. In future work, we plan to leverage the fast speed of database search using fragment ion indexing and indexed MS1 and MS/MS signal processing to support direct peptide identification from diaPASEF data in MSFragger.

In summary, we have developed a new direct DIA peptide identification method and implemented it as a module of the MSFragger search engine. By integrating MSFragger-DIA into the FragPipe computational platform, users can perform one-stop DIA data analysis, from peptide identification to quantification, optionally with the use of auxiliary (e.g., DDA) data to achieve optimal performance. With experiments covering various data acquisition schemes and sample types, we show that MSFragger-DIA demonstrates high sensitivity, fastest speed, and comparable quantification precision to other state-of-the-art DIA analysis tools. Coupled with the ease of use of FragPipe, we believe that this work describes an attractive computational solution for the analysis of a wide range of DIA datasets.

## Methods

### DIA data analysis pipelines

MSFragger and FragPipe can perform one-stop DIA data analysis with and without the use of auxiliary DDA data to build the library. The comparisons were done with the DIA-Umpire workflow in FragPipe based on pseudo-MS/MS generation, Spectronaut directDIA (library-free) analysis, DIA-NN *in silico* library-based analysis (also known as DIA-NN “library free” mode), and MaxDIA *in silico* library-based analyses. We used DIA-Umpire (version 2.2.8), Spectronaut results as reported in the original publications, DIA-NN (version 1.8.1), MaxDIA (version 2.1.3.0 for the runtime comparison and 2.1.0.0 for other experiments), and MSFragger (version 3.5). The pipelines are briefly described below.

#### FP-MSF

Only DIA data were used in this pipeline. MSFragger-DIA was used to directly search the DIA data. The search results were processed using MSBooster for deep learning-based score calculation, Percolator [33] for rescoring and posterior error probability calculation, ProteinProphet [34] for protein inference, Philosopher [55] for FDR filtering, and EasyPQP for spectral library building. The peptide ions in the spectral library were filtered with 1% global peptide and protein FDR. The resulting library was passed to DIA-NN [23] to extract and quantify precursors, peptides, and proteins from the DIA data. For peptide-protein quantification roll-up, MaxLFQ normalization was performed using the R package available at https://github.com/vdemichev/diann-rpackage.

#### FP-MSF hybrid

Both DIA and DDA data were used in this pipeline. MSFragger in the DIA and DDA modes was used to search the DIA and DDA data, respectively. The subsequent steps are the same as those in the FP-MSF pipeline.

#### FP-DIAU

Only DIA data was used in this pipeline. DIA-Umpire was used to generate pseudo-MS/MS spectra from the DIA data. Then, MSFragger in DDA mode was used to search these spectra. The remaining steps are the same as those used in the FP-MSF pipeline.

#### FP-DIAU hybrid

Both DIA and DDA data were used in this pipeline. MSFragger in the DDA mode was used to search the DIA extracted pseudo-MS/MS spectra and DDA data. The remaining steps are the same as those used in the FP-MSF pipeline.

#### Spectronaut

The results were extracted from the sne files provided as part of the publication.

#### DIA-NN *in silico* library-based

Only DIA data was used in this pipeline. DIA-NN predicted an *in silico* spectral library from the protein sequence database and then used the library to search and quantify the DIA data. This mode of DIA-NN is also known as DIA-NN library free mode. MaxLFQ normalization was used for peptide–protein quantification roll-up.

#### MaxDIA

Only DIA data was used in this pipeline. MaxDIA [49] was used to analyze the DIA data using the *in silico* spectral library downloaded from https://datashare.biochem.mpg.de/s/qe1IqcKbz2j2Ruf?path=%2FDiscoveryLibraries. This is also known as the “MaxDIA discovery mode” [49]. Unless otherwise noted, MaxLFQ intensity was used.

### FDR evaluation

The dataset published by Frohlich et al. [56] was used to estimate the actual FDR. The Spectronaut (version 15) directDIA results were used as provided by the authors. We used the DIA and GPF-DIA runs in the DIA-NN *in silico* library and FP-MSF pipelines. We used all the DIA, GPF-DIA, and DDA runs in the FP-MSF hybrid analysis. The FASTA file was a combination of *H. sapiens, E. coli*, and common contaminant proteins (downloaded on February 18, 2022, UP000005640 and UP000000625, 24750 proteins). The enzyme was set to restricted trypsin (i.e., allowing cleavage before Proline). Carbamidomethyl cysteine was set as a fixed modification. Protein N-terminal acetylation and oxidation of methionine were set as the variable modifications. The maximum allowed number of missed cleavages was set to 1. For DIA-NN and MSFragger, the precursors were filtered with the combination of 1% run-specific precursor FDR, global precursor FDR, run-specific protein FDR, and global protein FDR. For Spectronaut, the precursors were filtered with the combination of 1% global precursor FDR, run-specific protein FDR, and global protein FDR. Detailed parameters are provided in **Supplementary Data 1**.

There are only *H. sapiens* peptides in the “Lymphnode” experiment. Thus, the actual FDR can be calculated as:

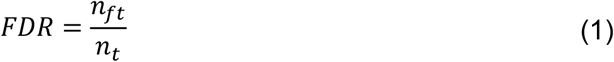

where *n*_*ft*_ is the number of false *H. sapiens* peptides and *n*_*t*_ is the total number of detected *H. sapiens* peptides. To estimate the actual FDR, we first have:

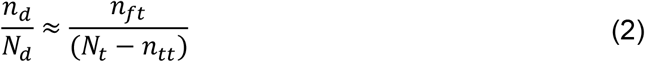

where *n*_*d*_ is the number of detected *E. coli* peptides, *N*_*d*_ is the number of *E. coli* peptides in the spectral library, *N*_*t*_ is the number of *H. sapiens* peptides in the spectral library, and *n*_*tt*_ is the number of detected true *H. sapiens* peptides. Combining **Equation (1)** and **(2)**, and assuming *n*_*tt*_ ≈ *n*_*t*_, we have:

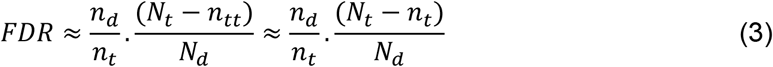

### Staggered isolation window data

We used two sets of DIA data acquired with staggered isolation windows to demonstrate the performance of MSFragger-DIA. The first dataset was published by Searle et al. [13] from HeLa samples acquired on a Thermo Q Exactive HF mass spectrometer. It contains six GPF-DIA runs with 4 Th isolation windows, and three single-injection DIA runs with 24 Th isolation windows. After demultiplexing [59], the effective isolation windows were 2 Th and 12 Th, respectively. We refer to this dataset as 2018-HeLa. The second dataset was published by Searle et al. [60] from *S. cerevisiae* strain BY4741 samples acquired on a Thermo Fusion Lumos mass spectrometer. It contains six GPF-DIA runs with 4 Th isolation windows and four single-injection DIA runs with 8 Th isolation windows. After demultiplexing, the effective isolation windows are 2 Th and 4 Th, respectively. We refer to this dataset as 2020-Yeast. The raw files were converted to mzML format using ProteoWizard [70] with the vendor’s peak picking and demultiplexing. Details of the sample preparation and data acquisition can be found in the original publications [13, 60].

DIA-NN *in silico* library-based, MaxDIA, and FP-MSF pipelines were used. The enzyme was set to restricted trypsin. Carbamidomethyl cysteine was set as a fixed modification. Protein N-terminal acetylation and oxidation of methionine were set as the variable modifications. The maximum allowed number of missed cleavages was set to 1. For MaxDIA pipeline, the *H. sapiens* and *S. cerevisiae* spectral libraries named “missed_cleavages_1” from the MaxQuant project website https://datashare.biochem.mpg.de/s/qe1IqcKbz2j2Ruf?path=%2FDiscoveryLibraries were used. For DIA-NN *in silico* library-based and FP-MSF pipelines, reviewed *H. sapiens* proteins and common contaminant sequences were downloaded from UniProt (downloaded on March 23, 2021, UP000005640, 20431 proteins) for 2018-HeLa dataset. The reviewed *S. cerevisiae* proteins and common contaminants were downloaded from UniProt (downloaded on March 16, 2021, UP000002311, 6164 proteins) for 2020-Yeast dataset. Decoy (reversed) sequences were appended to the original database for MSFragger-DIA analysis. For DIA-NN *in silico* library-based and FP-/MSF pipelines, the quantified peptides were filtered with the combination of 1% run-specific precursor FDR, run-specific protein FDR, global precursor FDR, and global protein FDR. For MaxDIA, the peptides were filtered with 1% global PSM and protein FDR. The detailed parameter files are provided in **Supplementary Data 1**.

### Low-input-cell data

A dataset from Siyal et al. [16] was used to demonstrate the performance of MSFragger-DIA in analyzing low-input data. We picked two experiments with 0.75 ng and 7.5 ng cells. Each experiment has three replicates. The authors also generated two sets of auxiliary DIA runs (“library runs”) for spectral library building, using samples with 1.5 ng and 1 μg of starting material. The database was obtained from the original publication (20387 proteins). Spectronaut (version 14) results were extracted from the sne files provided by the authors. FP-DIAU, DIA-NN *in silico* library-based, MaxDIA, and FP-MSF pipelines were used because no DDA data was available. For MaxDIA, the *H. sapiens* spectral library named “missed_cleavages_1” from https://datashare.biochem.mpg.de/s/qe1IqcKbz2j2Ruf?path=%2FDiscoveryLibraries was used. For FP-DIAU and FP-MSF workflows, the primary DIA data from the low-input cell runs and library DIA runs were used together to build a spectral library. For DIA-NN *in silico* library-based and FP-MSF pipelines, the proteins were filtered with the combination of 1% run-specific precursor FDR, run-specific protein FDR, global precursor FDR, and global protein FDR. For MaxDIA, proteins were filtered with 1% global PSM and protein FDR. For Spectronaut, the proteins were filtered with the combination of 1% global precursor and protein FDR. The remaining parameters are the same as those described in the preceding section (**Supplementary Data 1**).

### Single-cell data

A single-cell dataset published by Gebreyesus et al. [15] was used in this study. We picked two experiments, with 1 and 117 cells. Each experiment has three replicates. The authors also generated two sets of auxiliary runs for spectral library building. The first one has 9 DIA runs and 3 DDA runs from low-input data. We refer to this library as small-sample-lib. The second one has 4 DIA runs and 4 DDA runs from the bulk cells. We refer to it as large-sample-lib. For the 1 cell experiment, we used the small-sample-lib, as suggested by the authors. For the 117 cells experiment, we used the large-sample-lib. The protein sequence database was obtained from the original publication (20194 proteins). The Spectronaut (version 13) results were extracted from the sne files provided by the authors. FP-DIAU hybrid, DIA-NN *in silico* library-based, MaxDIA, FP-MSF, and FP-MSF hybrid pipelines were used. For the FP-DIAU and FP-MSF hybrid pipelines, both DIA and DDA data from the single-cell and library runs were used to generate a spectral library. For the DIA-NN *in silico* library-based, FP-MSF, and MaxDIA pipelines, the DIA data from the single-cell and library runs were used. The remaining parameters are the same as above (**Supplementary Data 1**).

### Phosphoproteome data

We used a phosphoproteome dataset obtained from Gao et al. [71]. There are six experiments from six melanoma cell lines. Each experiment has two replicates. FP-DIAU, Spectronaut, DIA-NN *in silico* library, and FP-MSF pipelines were used. MaxDIA was not used to analyze these data because no phosphoproteome spectral library was available on the corresponding tool website. The enzyme was set to restricted trypsin. Carbamidomethyl cysteine was set as a fixed modification. Protein N-terminal acetylation, oxidation of methionine, and phosphorylation of serine, threonine, and tyrosine were set as variable modifications. The maximum allowed number of missed cleavage was set to 1. The *H. sapiens* proteins and common contaminant sequences downloaded from UniProt (downloaded on March 23, 2021, UP000005640, 20431 proteins) were used as the target protein sequence database. The Spectronaut (version 14) sne files were downloaded from the original publications. The remaining parameters are the same as above (**Supplementary Data 1)**.

### Clear cell renal cell carcinoma (ccRCC) data

In the clear cell renal cell carcinoma (ccRCC) cohort [63], there are 187 single-injection DIA runs from normal and tumor tissues, and 8 fractionated DDA runs from pooled tissues generated for spectral library building. We used FP-DIAU, FP-DIAU hybrid, DIA-NN *in silico* library, FP-MSF, and FP-MSF hybrid pipelines to analyze the data. The enzyme was set to restricted trypsin. Carbamidomethyl cysteine was set as a fixed modification. Protein N-terminal acetylation and oxidation of methionine were set as variable modifications. The maximum allowed number of missed cleavages was set to 1. The database contains the reviewed *H. sapiens* proteins and common contaminant sequences downloaded from UniProt (downloaded on March 23, 2021, UP000005640, 20431 proteins). The detailed parameter files can be found in **Supplementary Data 1**.

### Speed benchmarks

The runtime for each pipeline was measured on two computers: (1) Windows desktop: Intel Core i9-10900K, 3.70GHz, 10 cores, 20 logical processors, 128Gb of memory; and (2) Linux server: Intel Xeon E5-2690 v4, 2.6 GHz, 28 cores, 56 logical processors, and 512 GB of memory. We used 19 threads on Windows desktop and 56 threads on the Linux server.

## Acknowledgements

This work was funded in part by NIH grants R01-GM-094231, U24-CA210967, U24-CA271037 (to AIN), as well as German Ministry of Education and Research (BMBF), as part of the National Research Node “Mass spectrometry in Systems Medicine” (MSCoreSys), under grant agreement 161L0221 (to VD). We thank Sarah Haynes for help with the manuscript, George Rosenberger for help with EasyPQP, and Kai Li for adopting the PDV [72] viewer in FragPipe to support MSFragger-DIA output. We also thank Michael MacCoss for the helpful discussions.

## Software Availability

MSFragger-DIA and MSFragger can be downloaded as a single JAR binary file at https://msfragger.nesvilab.org/. FragPipe is available on GitHub at https://github.com/Nesvilab/FragPipe. The source code for generating the figures is available at https://github.com/Nesvilab/MSFragger-DIA-manuscript.

## Data Availability

The result files from all tools used in this manuscript can be found from https://doi.org/10.5281/zenodo.7261712.

## Author Contributions

F.Y. developed MSFragger-DIA, A.T.K. contributed to the algorithm and software development at an early stage of the project. G.C.T. made the FragPipe support Percolator and EasyPQP. V.D. modified DIA-NN for FragPipe and contributed to data analysis. F.Y., G.X.L., and A.I.N. analyzed the data. F.Y. and A.I.N. wrote the manuscript with input from all authors, and A.I.N. conceived and supervised the project.

## Competing Interests Statement

The authors declare no competing financial interests.

## Notes

### Competing Interest Statement

The authors have declared no competing interest.

## References

1. Kitata, R.B., J.C. Yang, and Y.J. Chen, Advances in data-independent acquisition mass spectrometry towards comprehensive digital proteome landscape. Mass Spectrom Rev, 2022: p. e21781.

2. Ludwig, C., et al., Data-independent acquisition-based SWATH-MS for quantitative proteomics: a tutorial. Mol Syst Biol, 2018. 14(8): p. e8126.

3. Robinson, A.E., et al., Lysine and Arginine Protein Post-translational Modifications by Enhanced DIA Libraries: Quantification in Murine Liver Disease. J Proteome Res, 2020. 19(10): p. 4163–4178.

4. Kitata, R.B., et al., A data-independent acquisition-based global phosphoproteomics system enables deep profiling. Nat Commun, 2021. 12(1): p. 2539.

5. Steger, M., et al., Time-resolved in vivo ubiquitinome profiling by DIA-MS reveals USP7 targets on a proteome-wide scale. Nat Commun, 2021. 12(1): p. 5399.

6. Bekker-Jensen, D.B., et al., Rapid and site-specific deep phosphoproteome profiling by data-independent acquisition without the need for spectral libraries. Nature Communications, 2020. 11(1): p. 787.

7. Lambert, J.P., et al., Mapping differential interactomes by affinity purification coupled with data-independent mass spectrometry acquisition. Nat Methods, 2013. 10(12): p. 1239–45.

8. Fossati, A., et al., PCprophet: a framework for protein complex prediction and differential analysis using proteomic data. Nat Methods, 2021. 18(5): p. 520–527.

9. Caron, E., et al., An open-source computational and data resource to analyze digital maps of immunopeptidomes. Elife, 2015. 4.

10. Pak, H., et al., Sensitive Immunopeptidomics by Leveraging Available Large-Scale Multi-HLA Spectral Libraries, Data-Independent Acquisition, and MS/MS Prediction. Mol Cell Proteomics, 2021. 20: p. 100080.

11. Ritz, D., et al., Data-Independent Acquisition of HLA Class I Peptidomes on the Q Exactive Mass Spectrometer Platform. Proteomics, 2017. 17(19).

12. Liu, Y., et al., Quantitative variability of 342 plasma proteins in a human twin population. Mol Syst Biol, 2015. 11(1): p. 786.

13. Searle, B.C., et al., Chromatogram libraries improve peptide detection and quantification by data independent acquisition mass spectrometry. Nat Commun, 2018. 9(1): p. 5128.

14. Heil, L.R., et al., Building Spectral Libraries from Narrow-Window Data-Independent Acquisition Mass Spectrometry Data. J Proteome Res, 2022. 21(6): p. 1382–1391.

15. Gebreyesus, S.T., et al., Streamlined single-cell proteomics by an integrated microfluidic chip and data-independent acquisition mass spectrometry. Nat Commun, 2022. 13(1): p. 37.

16. Siyal, A.A., et al., Sample Size-Comparable Spectral Library Enhances Data-Independent Acquisition-Based Proteome Coverage of Low-Input Cells. Anal Chem, 2021. 93(51): p. 17003–17011.

17. Brunner, A.D., et al., Ultra-high sensitivity mass spectrometry quantifies single-cell proteome changes upon perturbation. Mol Syst Biol, 2022. 18(3): p. e10798.

18. Cho, K.C., et al., Deep Proteomics Using Two Dimensional Data Independent Acquisition Mass Spectrometry. Anal Chem, 2020. 92(6): p. 4217–4225.

19. Tsou, C.C., et al., DIA-Umpire: comprehensive computational framework for data-independent acquisition proteomics. Nat Methods, 2015. 12(3): p. 258–264.

20. MacLean, B., et al., Skyline: an open source document editor for creating and analyzing targeted proteomics experiments. Bioinformatics, 2010. 26(7): p. 966–8.

21. Rost, H.L., et al., OpenSWATH enables automated, targeted analysis of data-independent acquisition MS data. Nat Biotechnol, 2014. 32(3): p. 219–23.

22. Bruderer, R., et al., Extending the limits of quantitative proteome profiling with data-independent acquisition and application to acetaminophen-treated three-dimensional liver microtissues. Mol Cell Proteomics, 2015. 14(5): p. 1400–10.

23. Demichev, V., et al., DIA-NN: neural networks and interference correction enable deep proteome coverage in high throughput. Nat Methods, 2020. 17(1): p. 41–44.

24. Teo, G., et al., mapDIA: Preprocessing and statistical analysis of quantitative proteomics data from data independent acquisition mass spectrometry. J Proteomics, 2015. 129: p. 108–120.

25. Tsai, T.H., et al., Selection of Features with Consistent Profiles Improves Relative Protein Quantification in Mass Spectrometry Experiments. Mol Cell Proteomics, 2020. 19(6): p. 944–959.

26. Parker, S.J., V. Venkatraman, and J.E. Van Eyk, Effect of peptide assay library size and composition in targeted data-independent acquisition-MS analyses. Proteomics, 2016. 16(15-16): p. 2221–37.

27. Rosenberger, G., et al., Statistical control of peptide and protein error rates in large-scale targeted data-independent acquisition analyses. Nat Methods, 2017. 14(9): p. 921–927.

28. Barkovits, K., et al., Reproducibility, Specificity and Accuracy of Relative Quantification Using Spectral Library-based Data-independent Acquisition. Mol Cell Proteomics, 2020. 19(1): p. 181–197.

29. Kong, A.T., et al., MSFragger: ultrafast and comprehensive peptide identification in mass spectrometry-based proteomics. Nat Methods, 2017. 14(5): p. 513–520.

30. Craig, R. and R.C. Beavis, TANDEM: matching proteins with tandem mass spectra. Bioinformatics, 2004. 20(9): p. 1466–7.

31. Eng, J.K., T.A. Jahan, and M.R. Hoopmann, Comet: an open-source MS/MS sequence database search tool. Proteomics, 2013. 13(1): p. 22–4.

32. Keller, A., et al., Empirical statistical model to estimate the accuracy of peptide identifications made by MS/MS and database search. Anal Chem, 2002. 74(20): p. 5383–92.

33. Kall, L., et al., Semi-supervised learning for peptide identification from shotgun proteomics datasets. Nat Methods, 2007. 4(11): p. 923–5.

34. Nesvizhskii, A.I., et al., A statistical model for identifying proteins by tandem mass spectrometry. Anal Chem, 2003. 75(17): p. 4646–58.

35. Nesvizhskii, A.I., A survey of computational methods and error rate estimation procedures for peptide and protein identification in shotgun proteomics. J Proteomics, 2010. 73(11): p. 2092–123.

36. Ting, Y.S., et al., Peptide-Centric Proteome Analysis: An Alternative Strategy for the Analysis of Tandem Mass Spectrometry Data. Mol Cell Proteomics, 2015. 14(9): p. 2301–7.

37. Ting, Y.S., et al., PECAN: library-free peptide detection for data-independent acquisition tandem mass spectrometry data. Nat Methods, 2017. 14(9): p. 903–908.

38. Lu, Y.Y., et al., DIAmeter: matching peptides to data-independent acquisition mass spectrometry data. Bioinformatics, 2021. 37(Suppl_1): p. i434–i442.

39. Wang, J., et al., MSPLIT-DIA: sensitive peptide identification for data-independent acquisition. Nat Methods, 2015. 12(12): p. 1106–8.

40. Gessulat, S., et al., Prosit: proteome-wide prediction of peptide tandem mass spectra by deep learning. Nat Methods, 2019. 16(6): p. 509–518.

41. Zhou, X.X., et al., pDeep: Predicting MS/MS Spectra of Peptides with Deep Learning. Anal Chem, 2017. 89(23): p. 12690–12697.

42. Zeng, W.F., et al., MS/MS Spectrum Prediction for Modified Peptides Using pDeep2 Trained by Transfer Learning. Anal Chem, 2019. 91(15): p. 9724–9731.

43. Tiwary, S., et al., High-quality MS/MS spectrum prediction for data-dependent and data-independent acquisition data analysis. Nat Methods, 2019. 16(6): p. 519–525.

44. Tarn, C. and W.F. Zeng, pDeep3: Toward More Accurate Spectrum Prediction with Fast Few-Shot Learning. Anal Chem, 2021. 93(14): p. 5815–5822.

45. Yang, Y., et al., In silico spectral libraries by deep learning facilitate data-independent acquisition proteomics. Nat Commun, 2020. 11(1): p. 146.

46. Lou, R., et al., DeepPhospho accelerates DIA phosphoproteome profiling through in silico library generation. Nat Commun, 2021. 12(1): p. 6685.

47. Gotti, C., et al., Extensive and Accurate Benchmarking of DIA Acquisition Methods and Software Tools Using a Complex Proteomic Standard. J Proteome Res, 2021. 20(10): p. 4801–4814.

48. Demichev, V., et al., dia-PASEF data analysis using FragPipe and DIA-NN for deep proteomics of low sample amounts. Nat Commun, 2022. 13(1): p. 3944.

49. Sinitcyn, P., et al., MaxDIA enables library-based and library-free data-independent acquisition proteomics. Nat Biotechnol, 2021.

50. Teo, G.C., et al., Fast Deisotoping Algorithm and Its Implementation in the MSFragger Search Engine. J Proteome Res, 2021. 20(1): p. 498–505.

51. Yu, F., et al., Identification of modified peptides using localization-aware open search. Nat Commun, 2020. 11(1): p. 4065.

52. Yu, F., et al., Fast Quantitative Analysis of timsTOF PASEF Data with MSFragger and IonQuant. Mol Cell Proteomics, 2020. 19(9): p. 1575–1585.

53. Yu, F., S.E. Haynes, and A.I. Nesvizhskii, IonQuant Enables Accurate and Sensitive Label-Free Quantification With FDR-Controlled Match-Between-Runs. Mol Cell Proteomics, 2021. 20: p. 100077.

54. Yang, K.L., et al., MSBooster: Improving Peptide Identification Rates using Deep Learning-Based Features. bioRxiv, 2022: p. 2022.10.19.512904.

55. Leprevost, F.V., et al., Philosopher: a versatile toolkit for shotgun proteomics data analysis. Nature Methods, 2020. 17(9): p. 869–870.

56. Frohlich, K., et al., Benchmarking of analysis strategies for data-independent acquisition proteomics using a large-scale dataset comprising inter-patient heterogeneity. Nat Commun, 2022. 13(1): p. 2622.

57. Egertson, J.D., et al., Multiplexed MS/MS for improved data-independent acquisition. Nature Methods, 2013. 10(8): p. 744–746.

58. Pino, L.K., et al., Acquiring and Analyzing Data Independent Acquisition Proteomics Experiments without Spectrum Libraries. Mol Cell Proteomics, 2020. 19(7): p. 1088–1103.

59. Amodei, D., et al., Improving Precursor Selectivity in Data-Independent Acquisition Using Overlapping Windows. J Am Soc Mass Spectrom, 2019. 30(4): p. 669–684.

60. Searle, B.C., et al., Generating high quality libraries for DIA MS with empirically corrected peptide predictions. Nat Commun, 2020. 11(1): p. 1548.

61. Cox, J., et al., Accurate proteome-wide label-free quantification by delayed normalization and maximal peptide ratio extraction, termed MaxLFQ. Mol Cell Proteomics, 2014. 13(9): p. 2513–26.

62. Gao, E., et al., Data-independent acquisition-based proteome and phosphoproteome profiling across six melanoma cell lines reveals determinants of proteotypes. Mol Omics, 2021. 17(3): p. 413–425.

63. Clark, D.J., et al., Integrated Proteogenomic Characterization of Clear Cell Renal Cell Carcinoma. Cell, 2019. 179(4): p. 964-983.e31.

64. Allen, C., et al., nf-encyclopedia: A cloud-ready pipeline for chromatogram library data-independent acquisition proteomics workflows. 2022: p. 2022.09.30.510329.

65. Mertins, P., et al., Reproducible workflow for multiplexed deep-scale proteome and phosphoproteome analysis of tumor tissues by liquid chromatography–mass spectrometry. Nature Protocols, 2018. 13(7): p. 1632–1661.

66. Wen, B., E.J. Jaehnig, and B. Zhang, OmicsEV: a tool for comprehensive quality evaluation of omics data tables. Bioinformatics, 2022.

67. Truong, T., et al., Data-Dependent Acquisition with Precursor Coisolation Improves Proteome Coverage and Measurement Throughput for Label-Free Single-Cell Proteomics. bioRxiv, 2022: p. 2022.10.18.512791.

68. Mayer, R.L., et al., Wide Window Acquisition and AI-based data analysis to reach deep proteome coverage for a wide sample range, including single cell proteomic inputs. bioRxiv, 2022: p. 2022.09.01.506203.

69. Meier, F., et al., diaPASEF: parallel accumulation-serial fragmentation combined with data-independent acquisition. Nature Methods, 2020. 17(12): p. 1229-+.

70. Chambers, M.C., et al., A cross-platform toolkit for mass spectrometry and proteomics. Nat Biotechnol, 2012. 30(10): p. 918–20.

71. Gao, E., et al., Data-independent acquisition-based proteome and phosphoproteome profiling across six melanoma cell lines reveals determinants of proteotypes. Mol Omics, 2021.

72. Li, K., et al., PDV: an integrative proteomics data viewer. Bioinformatics, 2019. 35(7): p. 1249–1251.

