## Supplementary Data 1 for "One-stop analysis of DIA proteomics data using MSFragger-DIA and FragPipe computational platform": diann-output.log.txt

DIA-NN 1.8.1 (Data-Independent Acquisition by Neural Networks)
Compiled on Apr 15 2022 08:45:18
Current date and time: Fri Aug 19 15:42:56 2022
Logical CPU cores: 56
/storage/yufe/tools/FragPipe-18.0/fragpipe/tools/diann/1.8.1/linux/diann-1.8.1 --lib library.tsv --threads 56 --verbose 1 --out diann-output.tsv --qvalue 0.01 --matrix-spec-q 0.01 --matrices --no-prot-inf --smart-profiling --no-quant-files --peak-center --no-ifs-removal --predictor --dl-no-rt --dl-no-im --strip-unknown-mods --cfg /storage/yufe/results/msfragger_dia_paper/2020_narrow_wide_yeast/msfraggerdia/filelist_diann.txt 

Thread number set to 56
Output will be filtered at 0.01 FDR
Protein x sample matrices will be also filtered using run-specific protein q-value
Precursor/protein x samples expression level matrices will be saved along with the main report
Protein inference will not be performed
When generating a spectral library, in silico predicted spectra will be retained if deemed more reliable than experimental ones
.quant files will not be saved to the disk
Fixed-width center of each elution peak will be used for quantification
Interference removal from fragment elution curves disabled
Deep learning will be used to generate a new in silico spectral library from peptides provided
RTs will not be predicted using deep learning
IMs will not be predicted using deep learning
DIA-NN will use deep learning to predict spectra/RTs/IMs even for peptides carrying modifications which are not recognised by the deep learning predictor. In this scenario, if also generating a spectral library from the DIA data or using the MBR mode, it might or might not be better (depends on the data) to also use the --out-measured-rt option - it's recommended to test it with and without this option
DIA-NN will optimise the mass accuracy automatically using the first run in the experiment. This is useful primarily for quick initial analyses, when it is not yet known which mass accuracy setting works best for a particular acquisition scheme.

4 files will be processed
[0:00] Loading spectral library library.tsv
[0:01] Finding proteotypic peptides (assuming that the list of UniProt ids provided for each peptide is complete)
[0:01] Spectral library loaded: 4784 protein isoforms, 4784 protein groups and 91968 precursors in 72273 elution groups.
[0:04] [0:05] [0:13] [0:13] Saving the library to lib.predicted.speclib
[0:13] Initialising library

[0:13] File #1/4
[0:13] Loading run /storage/yufe/data/MSV000084000/mzml_sim_as_spectra/20190206_LUM1_CPBA_EASY04_060_30_SA_90mingrad_80B_DIA_400_1000_8mzol_15k_20IIT_4e5agc_1633-01_02.mzML
[4:05] 91968 library precursors are potentially detectable
[4:05] Processing...
[4:06] RT window set to 2.26582
[4:06] Peak width: 3.204
[4:06] Scan window radius set to 7
[4:06] Recommended MS1 mass accuracy setting: 4.26251 ppm
[4:07] Optimised mass accuracy: 20.08 ppm
[4:08] Removing low confidence identifications
[4:08] Removing interfering precursors
[4:10] Training neural networks: 82563 targets, 57224 decoys
[4:13] Number of IDs at 0.01 FDR: 73027
[4:14] Calculating protein q-values
[4:14] Number of protein isoforms identified at 1% FDR: 4523 (precursor-level), 4473 (protein-level) (inference performed using proteotypic peptides only)
[4:14] Quantification

[4:15] File #2/4
[4:15] Loading run /storage/yufe/data/MSV000084000/mzml_sim_as_spectra/20190206_LUM1_CPBA_EASY04_060_30_SA_90mingrad_80B_DIA_400_1000_8mzol_15k_20IIT_4e5agc_1633-01_01.mzML
[7:57] 91968 library precursors are potentially detectable
[7:57] Processing...
[7:58] RT window set to 2.29622
[7:58] Recommended MS1 mass accuracy setting: 4.90608 ppm
[7:59] Removing low confidence identifications
[7:59] Removing interfering precursors
[8:01] Training neural networks: 81880 targets, 55705 decoys
[8:04] Number of IDs at 0.01 FDR: 71211
[8:05] Calculating protein q-values
[8:05] Number of protein isoforms identified at 1% FDR: 4482 (precursor-level), 4390 (protein-level) (inference performed using proteotypic peptides only)
[8:05] Quantification

[8:06] File #3/4
[8:06] Loading run /storage/yufe/data/MSV000084000/mzml_sim_as_spectra/20190206_LUM1_CPBA_EASY04_060_30_SA_90mingrad_80B_DIA_400_1000_8mzol_15k_20IIT_4e5agc_1633-01_04.mzML
[11:43] 91968 library precursors are potentially detectable
[11:43] Processing...
[11:44] RT window set to 2.28876
[11:44] Recommended MS1 mass accuracy setting: 5.16792 ppm
[11:46] Removing low confidence identifications
[11:46] Removing interfering precursors
[11:48] Training neural networks: 80238 targets, 53939 decoys
[11:51] Number of IDs at 0.01 FDR: 68390
[11:52] Calculating protein q-values
[11:52] Number of protein isoforms identified at 1% FDR: 4438 (precursor-level), 4368 (protein-level) (inference performed using proteotypic peptides only)
[11:52] Quantification

[11:52] File #4/4
[11:52] Loading run /storage/yufe/data/MSV000084000/mzml_sim_as_spectra/20190206_LUM1_CPBA_EASY04_060_30_SA_90mingrad_80B_DIA_400_1000_8mzol_15k_20IIT_4e5agc_1633-01_05.mzML
[15:45] 91968 library precursors are potentially detectable
[15:45] Processing...
[15:46] RT window set to 2.27936
[15:46] Recommended MS1 mass accuracy setting: 4.42215 ppm
[15:47] Removing low confidence identifications
[15:47] Removing interfering precursors
[15:49] Training neural networks: 78245 targets, 49994 decoys
[15:52] Number of IDs at 0.01 FDR: 66125
[15:53] Calculating protein q-values
[15:53] Number of protein isoforms identified at 1% FDR: 4398 (precursor-level), 4302 (protein-level) (inference performed using proteotypic peptides only)
[15:53] Quantification

[15:54] Cross-run analysis
[15:54] Reading quantification information: 4 files
[15:54] Quantifying peptides
[15:59] Quantifying proteins
[15:59] Calculating q-values for protein and gene groups
[15:59] Calculating global q-values for protein and gene groups
[15:59] Writing report
[16:09] Report saved to diann-output.tsv.
[16:09] Saving precursor levels matrix
[16:10] Precursor levels matrix (1% precursor and protein group FDR) saved to diann-output.pr_matrix.tsv.
[16:10] Saving protein group levels matrix
[16:10] Protein group levels matrix (1% precursor FDR and protein group FDR) saved to diann-output.pg_matrix.tsv.
[16:10] Saving gene group levels matrix
[16:10] Gene groups levels matrix (1% precursor FDR and protein group FDR) saved to diann-output.gg_matrix.tsv.
[16:10] Saving unique genes levels matrix
[16:10] Unique genes levels matrix (1% precursor FDR and protein group FDR) saved to diann-output.unique_genes_matrix.tsv.
[16:10] Stats report saved to diann-output.stats.tsv

Finished
