## Supplementary Data 1 for "One-stop analysis of DIA proteomics data using MSFragger-DIA and FragPipe computational platform": 07_29_2022 22_15_14.txt

diann.exe --f "C:\Data\ccRCC\CCRCC_Discovery_20\mzml\CPTAC_CCRCC_W_JHU_20190112_LUMOS_C3L-00004_NAT.mzML
" --f "C:\Data\ccRCC\CCRCC_Discovery_20\mzml\CPTAC_CCRCC_W_JHU_20190112_LUMOS_C3L-00004_T.mzML
" --f "C:\Data\ccRCC\CCRCC_Discovery_20\mzml\CPTAC_CCRCC_W_JHU_20190112_LUMOS_C3L-00010_NAT.mzML
" --f "C:\Data\ccRCC\CCRCC_Discovery_20\mzml\CPTAC_CCRCC_W_JHU_20190112_LUMOS_C3L-00010_T.mzML
" --f "C:\Data\ccRCC\CCRCC_Discovery_20\mzml\CPTAC_CCRCC_W_JHU_20190112_LUMOS_C3L-00011_NAT.mzML
" --f "C:\Data\ccRCC\CCRCC_Discovery_20\mzml\CPTAC_CCRCC_W_JHU_20190112_LUMOS_C3L-00011_T.mzML
" --f "C:\Data\ccRCC\CCRCC_Discovery_20\mzml\CPTAC_CCRCC_W_JHU_20190112_LUMOS_C3L-00026_NAT.mzML
" --f "C:\Data\ccRCC\CCRCC_Discovery_20\mzml\CPTAC_CCRCC_W_JHU_20190112_LUMOS_C3L-00026_T.mzML
" --f "C:\Data\ccRCC\CCRCC_Discovery_20\mzml\CPTAC_CCRCC_W_JHU_20190112_LUMOS_C3L-00079_NAT.mzML
" --f "C:\Data\ccRCC\CCRCC_Discovery_20\mzml\CPTAC_CCRCC_W_JHU_20190112_LUMOS_C3L-00079_T.mzML
" --f "C:\Data\ccRCC\CCRCC_Discovery_20\mzml\CPTAC_CCRCC_W_JHU_20190112_LUMOS_C3L-00088_NAT.mzML
" --f "C:\Data\ccRCC\CCRCC_Discovery_20\mzml\CPTAC_CCRCC_W_JHU_20190112_LUMOS_C3L-00088_T.mzML
" --f "C:\Data\ccRCC\CCRCC_Discovery_20\mzml\CPTAC_CCRCC_W_JHU_20190112_LUMOS_C3L-00096_NAT.mzML
" --f "C:\Data\ccRCC\CCRCC_Discovery_20\mzml\CPTAC_CCRCC_W_JHU_20190112_LUMOS_C3L-00096_T.mzML
" --f "C:\Data\ccRCC\CCRCC_Discovery_20\mzml\CPTAC_CCRCC_W_JHU_20190112_LUMOS_C3L-00097_NAT.mzML
" --f "C:\Data\ccRCC\CCRCC_Discovery_20\mzml\CPTAC_CCRCC_W_JHU_20190112_LUMOS_C3L-00097_T.mzML
" --f "C:\Data\ccRCC\CCRCC_Discovery_20\mzml\CPTAC_CCRCC_W_JHU_20190112_LUMOS_C3L-00103_NAT.mzML
" --f "C:\Data\ccRCC\CCRCC_Discovery_20\mzml\CPTAC_CCRCC_W_JHU_20190112_LUMOS_C3L-00103_T.mzML
" --f "C:\Data\ccRCC\CCRCC_Discovery_20\mzml\CPTAC_CCRCC_W_JHU_20190112_LUMOS_C3L-00183_NAT.mzML
" --f "C:\Data\ccRCC\CCRCC_Discovery_20\mzml\CPTAC_CCRCC_W_JHU_20190112_LUMOS_C3L-00183_T.mzML
" --lib "" --threads 19 --verbose 1 --out "C:\Data\ccRCC\CCRCC_Discovery_20\DIANN-results_max3\report.tsv" --qvalue 0.01 --matrices  --out-lib "C:\Data\ccRCC\CCRCC_Discovery_20\DIANN-results_max3\report-lib.tsv" --gen-spec-lib --predictor --fasta "C:\Data\PXD022992_DIA_Melanoma\2021-03-23-reviewed-contam-UP000005640.fas" --fasta-search --min-fr-mz 200 --max-fr-mz 1800 --met-excision --cut K*,R* --missed-cleavages 1 --min-pep-len 7 --max-pep-len 30 --min-pr-mz 300 --max-pr-mz 1800 --min-pr-charge 1 --max-pr-charge 4 --unimod4 --var-mods 3 --var-mod UniMod:35,15.994915,M --var-mod UniMod:1,42.010565,*n --monitor-mod UniMod:1 --reanalyse --relaxed-prot-inf --smart-profiling --peak-center --no-ifs-removal 
DIA-NN 1.8.1 (Data-Independent Acquisition by Neural Networks)
Compiled on Apr 14 2022 15:31:19
Current date and time: Mon Aug  8 22:40:54 2022
CPU: GenuineIntel Intel(R) Core(TM) i9-10900K CPU @ 3.70GHz
SIMD instructions: AVX AVX2 FMA SSE4.1 SSE4.2 
Logical CPU cores: 20
Thread number set to 19
Output will be filtered at 0.01 FDR
Precursor/protein x samples expression level matrices will be saved along with the main report
A spectral library will be generated
Deep learning will be used to generate a new in silico spectral library from peptides provided
Library-free search enabled
Min fragment m/z set to 200
Max fragment m/z set to 1800
N-terminal methionine excision enabled
In silico digest will involve cuts at K*,R*
Maximum number of missed cleavages set to 1
Min peptide length set to 7
Max peptide length set to 30
Min precursor m/z set to 300
Max precursor m/z set to 1800
Min precursor charge set to 1
Max precursor charge set to 4
Cysteine carbamidomethylation enabled as a fixed modification
Maximum number of variable modifications set to 3
Modification UniMod:35 with mass delta 15.9949 at M will be considered as variable
Modification UniMod:1 with mass delta 42.0106 at *n will be considered as variable
A spectral library will be created from the DIA runs and used to reanalyse them; .quant files will only be saved to disk during the first step
Highly heuristic protein grouping will be used, to reduce the number of protein groups obtained; this mode is recommended for benchmarking protein ID numbers; use with caution for anything else
When generating a spectral library, in silico predicted spectra will be retained if deemed more reliable than experimental ones
Fixed-width center of each elution peak will be used for quantification
Interference removal from fragment elution curves disabled
DIA-NN will optimise the mass accuracy automatically using the first run in the experiment. This is useful primarily for quick initial analyses, when it is not yet known which mass accuracy setting works best for a particular acquisition scheme.
Exclusion of fragments shared between heavy and light peptides from quantification is not supported in FASTA digest mode - disabled; to enable, generate an in silico predicted spectral library and analyse with this library
The following variable modifications will be scored: UniMod:1 

20 files will be processed
[0:00] Loading FASTA C:\Data\PXD022992_DIA_Melanoma\2021-03-23-reviewed-contam-UP000005640.fas
[0:06] Processing FASTA
[0:16] Assembling elution groups
[0:23] 6190376 precursors generated
[0:23] Protein names missing for some isoforms
[0:23] Gene names missing for some isoforms
[0:23] Library contains 20406 proteins, and 20164 genes
[0:23] Encoding peptides for spectra and RTs prediction
[0:31] Predicting spectra and IMs
[26:21] Predicting RTs
[29:20] Decoding predicted spectra and IMs
[29:29] Decoding RTs
[29:33] Saving the library to C:\Data\ccRCC\CCRCC_Discovery_20\DIANN-results_max3\report-lib.predicted.speclib
[29:46] Initialising library

[29:48] First pass: generating a spectral library from DIA data
[29:48] File #1/20
[29:48] Loading run C:\Data\ccRCC\CCRCC_Discovery_20\mzml\CPTAC_CCRCC_W_JHU_20190112_LUMOS_C3L-00004_NAT.mzML
[29:58] 5621709 library precursors are potentially detectable
[29:59] Processing...
[31:34] RT window set to 9.01219
[31:34] Peak width: 3.996
[31:34] Scan window radius set to 8
[31:34] Recommended MS1 mass accuracy setting: 9.10451 ppm
[34:20] Optimised mass accuracy: 17.672 ppm
[46:30] Removing low confidence identifications
[46:30] Searching PTM decoys
[46:44] Removing interfering precursors
[46:49] Training neural networks: 143547 targets, 113655 decoys
[46:56] Number of IDs at 0.01 FDR: 76465
[46:57] Calculating protein q-values
[46:57] Number of genes identified at 1% FDR: 7222 (precursor-level), 6514 (protein-level) (inference performed using proteotypic peptides only)
[46:57] Quantification
[46:58] Precursors with monitored PTMs at 1% FDR: 822 out of 924
[46:58] Unmodified precursors with monitored PTM sites at 1% FDR: 205 out of 244
[46:59] Quantification information saved to C:\Data\ccRCC\CCRCC_Discovery_20\mzml\CPTAC_CCRCC_W_JHU_20190112_LUMOS_C3L-00004_NAT.mzML.quant.

[47:00] File #2/20
[47:00] Loading run C:\Data\ccRCC\CCRCC_Discovery_20\mzml\CPTAC_CCRCC_W_JHU_20190112_LUMOS_C3L-00004_T.mzML
[47:10] 5621709 library precursors are potentially detectable
[47:11] Processing...
[48:47] RT window set to 8.58556
[48:47] Recommended MS1 mass accuracy setting: 9.39879 ppm
[60:59] Removing low confidence identifications
[60:59] Searching PTM decoys
[61:13] Removing interfering precursors
[61:18] Training neural networks: 155746 targets, 122717 decoys
[61:26] Number of IDs at 0.01 FDR: 87545
[61:26] Calculating protein q-values
[61:27] Number of genes identified at 1% FDR: 7762 (precursor-level), 7013 (protein-level) (inference performed using proteotypic peptides only)
[61:27] Quantification
[61:28] Precursors with monitored PTMs at 1% FDR: 1025 out of 1157
[61:28] Unmodified precursors with monitored PTM sites at 1% FDR: 239 out of 272
[61:29] Quantification information saved to C:\Data\ccRCC\CCRCC_Discovery_20\mzml\CPTAC_CCRCC_W_JHU_20190112_LUMOS_C3L-00004_T.mzML.quant.

[61:30] File #3/20
[61:30] Loading run C:\Data\ccRCC\CCRCC_Discovery_20\mzml\CPTAC_CCRCC_W_JHU_20190112_LUMOS_C3L-00010_NAT.mzML
[61:40] 5621709 library precursors are potentially detectable
[61:40] Processing...
[63:00] RT window set to 9.32626
[63:00] Recommended MS1 mass accuracy setting: 8.93531 ppm
[75:40] Removing low confidence identifications
[75:40] Searching PTM decoys
[75:54] Removing interfering precursors
[75:59] Training neural networks: 151564 targets, 122745 decoys
[76:07] Number of IDs at 0.01 FDR: 83132
[76:08] Calculating protein q-values
[76:08] Number of genes identified at 1% FDR: 7546 (precursor-level), 6718 (protein-level) (inference performed using proteotypic peptides only)
[76:08] Quantification
[76:09] Precursors with monitored PTMs at 1% FDR: 854 out of 1021
[76:09] Unmodified precursors with monitored PTM sites at 1% FDR: 207 out of 263
[76:10] Quantification information saved to C:\Data\ccRCC\CCRCC_Discovery_20\mzml\CPTAC_CCRCC_W_JHU_20190112_LUMOS_C3L-00010_NAT.mzML.quant.

[76:11] File #4/20
[76:11] Loading run C:\Data\ccRCC\CCRCC_Discovery_20\mzml\CPTAC_CCRCC_W_JHU_20190112_LUMOS_C3L-00010_T.mzML
[76:21] 5621709 library precursors are potentially detectable
[76:22] Processing...
[78:13] RT window set to 9.44712
[78:14] Recommended MS1 mass accuracy setting: 9.98289 ppm
[90:56] Removing low confidence identifications
[90:56] Searching PTM decoys
[91:10] Removing interfering precursors
[91:14] Training neural networks: 128811 targets, 102310 decoys
[91:21] Number of IDs at 0.01 FDR: 69710
[91:22] Calculating protein q-values
[91:22] Number of genes identified at 1% FDR: 6944 (precursor-level), 6178 (protein-level) (inference performed using proteotypic peptides only)
[91:22] Quantification
[91:23] Precursors with monitored PTMs at 1% FDR: 609 out of 892
[91:23] Unmodified precursors with monitored PTM sites at 1% FDR: 137 out of 234
[91:24] Quantification information saved to C:\Data\ccRCC\CCRCC_Discovery_20\mzml\CPTAC_CCRCC_W_JHU_20190112_LUMOS_C3L-00010_T.mzML.quant.

[91:25] File #5/20
[91:25] Loading run C:\Data\ccRCC\CCRCC_Discovery_20\mzml\CPTAC_CCRCC_W_JHU_20190112_LUMOS_C3L-00011_NAT.mzML
[91:35] 5621709 library precursors are potentially detectable
[91:35] Processing...
[92:54] RT window set to 9.50545
[92:54] Recommended MS1 mass accuracy setting: 9.09886 ppm
[105:28] Removing low confidence identifications
[105:28] Searching PTM decoys
[105:42] Removing interfering precursors
[105:47] Training neural networks: 154857 targets, 123049 decoys
[105:55] Number of IDs at 0.01 FDR: 83554
[105:56] Calculating protein q-values
[105:56] Number of genes identified at 1% FDR: 7564 (precursor-level), 6842 (protein-level) (inference performed using proteotypic peptides only)
[105:56] Quantification
[105:57] Precursors with monitored PTMs at 1% FDR: 953 out of 1061
[105:57] Unmodified precursors with monitored PTM sites at 1% FDR: 249 out of 278
[105:58] Quantification information saved to C:\Data\ccRCC\CCRCC_Discovery_20\mzml\CPTAC_CCRCC_W_JHU_20190112_LUMOS_C3L-00011_NAT.mzML.quant.

[106:00] File #6/20
[106:00] Loading run C:\Data\ccRCC\CCRCC_Discovery_20\mzml\CPTAC_CCRCC_W_JHU_20190112_LUMOS_C3L-00011_T.mzML
[106:09] 5621709 library precursors are potentially detectable
[106:10] Processing...
[107:27] RT window set to 9.40998
[107:27] Recommended MS1 mass accuracy setting: 7.62707 ppm
[119:25] Removing low confidence identifications
[119:26] Searching PTM decoys
[119:39] Removing interfering precursors
[119:45] Training neural networks: 161592 targets, 130130 decoys
[119:53] Number of IDs at 0.01 FDR: 89964
[119:54] Calculating protein q-values
[119:54] Number of genes identified at 1% FDR: 7835 (precursor-level), 7025 (protein-level) (inference performed using proteotypic peptides only)
[119:54] Quantification
[119:55] Precursors with monitored PTMs at 1% FDR: 1074 out of 1242
[119:55] Unmodified precursors with monitored PTM sites at 1% FDR: 253 out of 305
[119:56] Quantification information saved to C:\Data\ccRCC\CCRCC_Discovery_20\mzml\CPTAC_CCRCC_W_JHU_20190112_LUMOS_C3L-00011_T.mzML.quant.

[119:57] File #7/20
[119:57] Loading run C:\Data\ccRCC\CCRCC_Discovery_20\mzml\CPTAC_CCRCC_W_JHU_20190112_LUMOS_C3L-00026_NAT.mzML
[120:07] 5621709 library precursors are potentially detectable
[120:07] Processing...
[121:36] RT window set to 9.48063
[121:36] Recommended MS1 mass accuracy setting: 8.2095 ppm
[133:33] Removing low confidence identifications
[133:34] Searching PTM decoys
[133:47] Removing interfering precursors
[133:52] Training neural networks: 149236 targets, 119597 decoys
[134:00] Number of IDs at 0.01 FDR: 81774
[134:01] Calculating protein q-values
[134:01] Number of genes identified at 1% FDR: 7593 (precursor-level), 6790 (protein-level) (inference performed using proteotypic peptides only)
[134:01] Quantification
[134:02] Precursors with monitored PTMs at 1% FDR: 874 out of 1021
[134:02] Unmodified precursors with monitored PTM sites at 1% FDR: 207 out of 257
[134:03] Quantification information saved to C:\Data\ccRCC\CCRCC_Discovery_20\mzml\CPTAC_CCRCC_W_JHU_20190112_LUMOS_C3L-00026_NAT.mzML.quant.

[134:04] File #8/20
[134:04] Loading run C:\Data\ccRCC\CCRCC_Discovery_20\mzml\CPTAC_CCRCC_W_JHU_20190112_LUMOS_C3L-00026_T.mzML
[134:14] 5621709 library precursors are potentially detectable
[134:14] Processing...
[136:13] RT window set to 9.65753
[136:14] Recommended MS1 mass accuracy setting: 9.25702 ppm
[148:02] Removing low confidence identifications
[148:02] Searching PTM decoys
[148:15] Removing interfering precursors
[148:20] Training neural networks: 118176 targets, 93568 decoys
[148:26] Number of IDs at 0.01 FDR: 61690
[148:26] Calculating protein q-values
[148:27] Number of genes identified at 1% FDR: 6587 (precursor-level), 5904 (protein-level) (inference performed using proteotypic peptides only)
[148:27] Quantification
[148:27] Precursors with monitored PTMs at 1% FDR: 831 out of 900
[148:27] Unmodified precursors with monitored PTM sites at 1% FDR: 185 out of 199
[148:28] Quantification information saved to C:\Data\ccRCC\CCRCC_Discovery_20\mzml\CPTAC_CCRCC_W_JHU_20190112_LUMOS_C3L-00026_T.mzML.quant.

[148:29] File #9/20
[148:29] Loading run C:\Data\ccRCC\CCRCC_Discovery_20\mzml\CPTAC_CCRCC_W_JHU_20190112_LUMOS_C3L-00079_NAT.mzML
[148:39] 5621709 library precursors are potentially detectable
[148:39] Processing...
[149:57] RT window set to 9.29417
[149:57] Recommended MS1 mass accuracy setting: 7.35298 ppm
[161:55] Removing low confidence identifications
[161:56] Searching PTM decoys
[162:09] Removing interfering precursors
[162:14] Training neural networks: 164301 targets, 131554 decoys
[162:23] Number of IDs at 0.01 FDR: 91434
[162:24] Calculating protein q-values
[162:24] Number of genes identified at 1% FDR: 8092 (precursor-level), 7368 (protein-level) (inference performed using proteotypic peptides only)
[162:24] Quantification
[162:25] Precursors with monitored PTMs at 1% FDR: 1105 out of 1214
[162:25] Unmodified precursors with monitored PTM sites at 1% FDR: 237 out of 266
[162:26] Quantification information saved to C:\Data\ccRCC\CCRCC_Discovery_20\mzml\CPTAC_CCRCC_W_JHU_20190112_LUMOS_C3L-00079_NAT.mzML.quant.

[162:27] File #10/20
[162:27] Loading run C:\Data\ccRCC\CCRCC_Discovery_20\mzml\CPTAC_CCRCC_W_JHU_20190112_LUMOS_C3L-00079_T.mzML
[162:37] 5621709 library precursors are potentially detectable
[162:37] Processing...
[163:54] RT window set to 9.34803
[163:55] Recommended MS1 mass accuracy setting: 8.94893 ppm
[175:46] Removing low confidence identifications
[175:46] Searching PTM decoys
[175:59] Removing interfering precursors
[176:05] Training neural networks: 158205 targets, 126867 decoys
[176:13] Number of IDs at 0.01 FDR: 88347
[176:14] Calculating protein q-values
[176:14] Number of genes identified at 1% FDR: 7993 (precursor-level), 7235 (protein-level) (inference performed using proteotypic peptides only)
[176:14] Quantification
[176:15] Precursors with monitored PTMs at 1% FDR: 960 out of 1133
[176:15] Unmodified precursors with monitored PTM sites at 1% FDR: 237 out of 274
[176:16] Quantification information saved to C:\Data\ccRCC\CCRCC_Discovery_20\mzml\CPTAC_CCRCC_W_JHU_20190112_LUMOS_C3L-00079_T.mzML.quant.

[176:17] File #11/20
[176:17] Loading run C:\Data\ccRCC\CCRCC_Discovery_20\mzml\CPTAC_CCRCC_W_JHU_20190112_LUMOS_C3L-00088_NAT.mzML
[176:27] 5621709 library precursors are potentially detectable
[176:27] Processing...
[177:57] RT window set to 9.29967
[177:58] Recommended MS1 mass accuracy setting: 7.67936 ppm
[190:00] Removing low confidence identifications
[190:00] Searching PTM decoys
[190:14] Removing interfering precursors
[190:19] Training neural networks: 153953 targets, 121514 decoys
[190:26] Number of IDs at 0.01 FDR: 83943
[190:27] Calculating protein q-values
[190:28] Number of genes identified at 1% FDR: 7718 (precursor-level), 6910 (protein-level) (inference performed using proteotypic peptides only)
[190:28] Quantification
[190:28] Precursors with monitored PTMs at 1% FDR: 903 out of 1094
[190:28] Unmodified precursors with monitored PTM sites at 1% FDR: 207 out of 264
[190:29] Quantification information saved to C:\Data\ccRCC\CCRCC_Discovery_20\mzml\CPTAC_CCRCC_W_JHU_20190112_LUMOS_C3L-00088_NAT.mzML.quant.

[190:31] File #12/20
[190:31] Loading run C:\Data\ccRCC\CCRCC_Discovery_20\mzml\CPTAC_CCRCC_W_JHU_20190112_LUMOS_C3L-00088_T.mzML
[190:40] 5621709 library precursors are potentially detectable
[190:41] Processing...
[192:27] RT window set to 9.26378
[192:28] Recommended MS1 mass accuracy setting: 9.34024 ppm
[204:36] Removing low confidence identifications
[204:36] Searching PTM decoys
[204:50] Removing interfering precursors
[204:54] Training neural networks: 131887 targets, 101908 decoys
[205:01] Number of IDs at 0.01 FDR: 70007
[205:02] Calculating protein q-values
[205:02] Number of genes identified at 1% FDR: 7034 (precursor-level), 6283 (protein-level) (inference performed using proteotypic peptides only)
[205:02] Quantification
[205:03] Precursors with monitored PTMs at 1% FDR: 730 out of 910
[205:03] Unmodified precursors with monitored PTM sites at 1% FDR: 159 out of 204
[205:04] Quantification information saved to C:\Data\ccRCC\CCRCC_Discovery_20\mzml\CPTAC_CCRCC_W_JHU_20190112_LUMOS_C3L-00088_T.mzML.quant.

[205:05] File #13/20
[205:05] Loading run C:\Data\ccRCC\CCRCC_Discovery_20\mzml\CPTAC_CCRCC_W_JHU_20190112_LUMOS_C3L-00096_NAT.mzML
[205:15] 5621709 library precursors are potentially detectable
[205:15] Processing...
[206:33] RT window set to 9.17502
[206:33] Recommended MS1 mass accuracy setting: 7.38256 ppm
[218:18] Removing low confidence identifications
[218:18] Searching PTM decoys
[218:32] Removing interfering precursors
[218:37] Training neural networks: 159626 targets, 127842 decoys
[218:45] Number of IDs at 0.01 FDR: 88446
[218:46] Calculating protein q-values
[218:46] Number of genes identified at 1% FDR: 7947 (precursor-level), 7195 (protein-level) (inference performed using proteotypic peptides only)
[218:46] Quantification
[218:47] Precursors with monitored PTMs at 1% FDR: 919 out of 1198
[218:47] Unmodified precursors with monitored PTM sites at 1% FDR: 189 out of 262
[218:48] Quantification information saved to C:\Data\ccRCC\CCRCC_Discovery_20\mzml\CPTAC_CCRCC_W_JHU_20190112_LUMOS_C3L-00096_NAT.mzML.quant.

[218:49] File #14/20
[218:49] Loading run C:\Data\ccRCC\CCRCC_Discovery_20\mzml\CPTAC_CCRCC_W_JHU_20190112_LUMOS_C3L-00096_T.mzML
[218:59] 5621709 library precursors are potentially detectable
[218:59] Processing...
[220:17] RT window set to 8.84149
[220:18] Recommended MS1 mass accuracy setting: 8.11784 ppm
[231:52] Removing low confidence identifications
[231:52] Searching PTM decoys
[232:05] Removing interfering precursors
[232:10] Training neural networks: 158701 targets, 126459 decoys
[232:18] Number of IDs at 0.01 FDR: 87469
[232:19] Calculating protein q-values
[232:20] Number of genes identified at 1% FDR: 7907 (precursor-level), 7209 (protein-level) (inference performed using proteotypic peptides only)
[232:20] Quantification
[232:20] Precursors with monitored PTMs at 1% FDR: 927 out of 1155
[232:20] Unmodified precursors with monitored PTM sites at 1% FDR: 206 out of 272
[232:21] Quantification information saved to C:\Data\ccRCC\CCRCC_Discovery_20\mzml\CPTAC_CCRCC_W_JHU_20190112_LUMOS_C3L-00096_T.mzML.quant.

[232:23] File #15/20
[232:23] Loading run C:\Data\ccRCC\CCRCC_Discovery_20\mzml\CPTAC_CCRCC_W_JHU_20190112_LUMOS_C3L-00097_NAT.mzML
[232:32] 5621709 library precursors are potentially detectable
[232:33] Processing...
[234:05] RT window set to 9.1028
[234:05] Recommended MS1 mass accuracy setting: 8.5046 ppm
[245:45] Removing low confidence identifications
[245:46] Searching PTM decoys
[245:59] Removing interfering precursors
[246:03] Training neural networks: 138468 targets, 109450 decoys
[246:10] Number of IDs at 0.01 FDR: 75548
[246:11] Calculating protein q-values
[246:11] Number of genes identified at 1% FDR: 7114 (precursor-level), 6337 (protein-level) (inference performed using proteotypic peptides only)
[246:11] Quantification
[246:12] Precursors with monitored PTMs at 1% FDR: 901 out of 1005
[246:12] Unmodified precursors with monitored PTM sites at 1% FDR: 209 out of 246
[246:13] Quantification information saved to C:\Data\ccRCC\CCRCC_Discovery_20\mzml\CPTAC_CCRCC_W_JHU_20190112_LUMOS_C3L-00097_NAT.mzML.quant.

[246:14] File #16/20
[246:14] Loading run C:\Data\ccRCC\CCRCC_Discovery_20\mzml\CPTAC_CCRCC_W_JHU_20190112_LUMOS_C3L-00097_T.mzML
[246:24] 5621709 library precursors are potentially detectable
[246:24] Processing...
[248:12] RT window set to 9.1673
[248:12] Recommended MS1 mass accuracy setting: 7.94611 ppm
[259:09] Removing low confidence identifications
[259:10] Searching PTM decoys
[259:22] Removing interfering precursors
[259:26] Training neural networks: 120373 targets, 95118 decoys
[259:32] Number of IDs at 0.01 FDR: 63928
[259:33] Calculating protein q-values
[259:33] Number of genes identified at 1% FDR: 6566 (precursor-level), 5879 (protein-level) (inference performed using proteotypic peptides only)
[259:33] Quantification
[259:34] Precursors with monitored PTMs at 1% FDR: 785 out of 873
[259:34] Unmodified precursors with monitored PTM sites at 1% FDR: 167 out of 193
[259:35] Quantification information saved to C:\Data\ccRCC\CCRCC_Discovery_20\mzml\CPTAC_CCRCC_W_JHU_20190112_LUMOS_C3L-00097_T.mzML.quant.

[259:36] File #17/20
[259:36] Loading run C:\Data\ccRCC\CCRCC_Discovery_20\mzml\CPTAC_CCRCC_W_JHU_20190112_LUMOS_C3L-00103_NAT.mzML
[259:46] 5621709 library precursors are potentially detectable
[259:46] Processing...
[261:03] RT window set to 8.84131
[261:03] Recommended MS1 mass accuracy setting: 7.86947 ppm
[272:21] Removing low confidence identifications
[272:21] Searching PTM decoys
[272:34] Removing interfering precursors
[272:39] Training neural networks: 162714 targets, 129749 decoys
[272:47] Number of IDs at 0.01 FDR: 91093
[272:48] Calculating protein q-values
[272:48] Number of genes identified at 1% FDR: 8044 (precursor-level), 7237 (protein-level) (inference performed using proteotypic peptides only)
[272:48] Quantification
[272:49] Precursors with monitored PTMs at 1% FDR: 1208 out of 1310
[272:49] Unmodified precursors with monitored PTM sites at 1% FDR: 246 out of 279
[272:50] Quantification information saved to C:\Data\ccRCC\CCRCC_Discovery_20\mzml\CPTAC_CCRCC_W_JHU_20190112_LUMOS_C3L-00103_NAT.mzML.quant.

[272:51] File #18/20
[272:51] Loading run C:\Data\ccRCC\CCRCC_Discovery_20\mzml\CPTAC_CCRCC_W_JHU_20190112_LUMOS_C3L-00103_T.mzML
[273:01] 5621709 library precursors are potentially detectable
[273:01] Processing...
[274:44] RT window set to 9.29698
[274:44] Recommended MS1 mass accuracy setting: 9.08225 ppm
[285:52] Removing low confidence identifications
[285:52] Searching PTM decoys
[286:05] Removing interfering precursors
[286:09] Training neural networks: 128868 targets, 100252 decoys
[286:16] Number of IDs at 0.01 FDR: 68877
[286:16] Calculating protein q-values
[286:17] Number of genes identified at 1% FDR: 6967 (precursor-level), 6269 (protein-level) (inference performed using proteotypic peptides only)
[286:17] Quantification
[286:17] Precursors with monitored PTMs at 1% FDR: 762 out of 821
[286:17] Unmodified precursors with monitored PTM sites at 1% FDR: 187 out of 211
[286:18] Quantification information saved to C:\Data\ccRCC\CCRCC_Discovery_20\mzml\CPTAC_CCRCC_W_JHU_20190112_LUMOS_C3L-00103_T.mzML.quant.

[286:20] File #19/20
[286:20] Loading run C:\Data\ccRCC\CCRCC_Discovery_20\mzml\CPTAC_CCRCC_W_JHU_20190112_LUMOS_C3L-00183_NAT.mzML
[286:29] 5621709 library precursors are potentially detectable
[286:30] Processing...
[288:01] RT window set to 8.87525
[288:01] Recommended MS1 mass accuracy setting: 7.7888 ppm
[299:07] Removing low confidence identifications
[299:07] Searching PTM decoys
[299:20] Removing interfering precursors
[299:24] Training neural networks: 139011 targets, 107608 decoys
[299:31] Number of IDs at 0.01 FDR: 73522
[299:32] Calculating protein q-values
[299:32] Number of genes identified at 1% FDR: 6918 (precursor-level), 6232 (protein-level) (inference performed using proteotypic peptides only)
[299:32] Quantification
[299:33] Precursors with monitored PTMs at 1% FDR: 845 out of 954
[299:33] Unmodified precursors with monitored PTM sites at 1% FDR: 213 out of 237
[299:34] Quantification information saved to C:\Data\ccRCC\CCRCC_Discovery_20\mzml\CPTAC_CCRCC_W_JHU_20190112_LUMOS_C3L-00183_NAT.mzML.quant.

[299:35] File #20/20
[299:35] Loading run C:\Data\ccRCC\CCRCC_Discovery_20\mzml\CPTAC_CCRCC_W_JHU_20190112_LUMOS_C3L-00183_T.mzML
[299:45] 5621709 library precursors are potentially detectable
[299:45] Processing...
[301:41] RT window set to 10.4751
[301:41] Recommended MS1 mass accuracy setting: 8.54079 ppm
[312:54] Removing low confidence identifications
[312:55] Searching PTM decoys
[313:07] Removing interfering precursors
[313:11] Training neural networks: 98830 targets, 78886 decoys
[313:16] Number of IDs at 0.01 FDR: 51503
[313:16] Calculating protein q-values
[313:17] Number of genes identified at 1% FDR: 5989 (precursor-level), 5383 (protein-level) (inference performed using proteotypic peptides only)
[313:17] Quantification
[313:17] Precursors with monitored PTMs at 1% FDR: 736 out of 813
[313:17] Unmodified precursors with monitored PTM sites at 1% FDR: 145 out of 164
[313:18] Quantification information saved to C:\Data\ccRCC\CCRCC_Discovery_20\mzml\CPTAC_CCRCC_W_JHU_20190112_LUMOS_C3L-00183_T.mzML.quant.

[313:20] Cross-run analysis
[313:20] Reading quantification information: 20 files
[313:24] Quantifying peptides
[313:36] Assembling protein groups
[313:39] Quantifying proteins
[313:40] Calculating q-values for protein and gene groups
[313:41] Calculating global q-values for protein and gene groups
[313:41] Writing report
[314:59] Report saved to C:\Data\ccRCC\CCRCC_Discovery_20\DIANN-results_max3\report-first-pass.tsv.
[314:59] Saving precursor levels matrix
[315:00] Precursor levels matrix (1% precursor and protein group FDR) saved to C:\Data\ccRCC\CCRCC_Discovery_20\DIANN-results_max3\report-first-pass.pr_matrix.tsv.
[315:00] Saving protein group levels matrix
[315:00] Protein group levels matrix (1% precursor FDR and protein group FDR) saved to C:\Data\ccRCC\CCRCC_Discovery_20\DIANN-results_max3\report-first-pass.pg_matrix.tsv.
[315:00] Saving gene group levels matrix
[315:01] Gene groups levels matrix (1% precursor FDR and protein group FDR) saved to C:\Data\ccRCC\CCRCC_Discovery_20\DIANN-results_max3\report-first-pass.gg_matrix.tsv.
[315:01] Saving unique genes levels matrix
[315:01] Unique genes levels matrix (1% precursor FDR and protein group FDR) saved to C:\Data\ccRCC\CCRCC_Discovery_20\DIANN-results_max3\report-first-pass.unique_genes_matrix.tsv.
[315:01] Stats report saved to C:\Data\ccRCC\CCRCC_Discovery_20\DIANN-results_max3\report-first-pass.stats.tsv
[315:01] Generating spectral library:
[315:01] 147797 precursors passing the FDR threshold are to be extracted
[315:01] Loading run C:\Data\ccRCC\CCRCC_Discovery_20\mzml\CPTAC_CCRCC_W_JHU_20190112_LUMOS_C3L-00004_NAT.mzML
[315:11] 5621709 library precursors are potentially detectable
[315:12] 8313 spectra added to the library
[315:13] Loading run C:\Data\ccRCC\CCRCC_Discovery_20\mzml\CPTAC_CCRCC_W_JHU_20190112_LUMOS_C3L-00004_T.mzML
[315:24] 5621709 library precursors are potentially detectable
[315:25] 6807 spectra added to the library
[315:25] Loading run C:\Data\ccRCC\CCRCC_Discovery_20\mzml\CPTAC_CCRCC_W_JHU_20190112_LUMOS_C3L-00010_NAT.mzML
[315:36] 5621709 library precursors are potentially detectable
[315:36] 4729 spectra added to the library
[315:37] Loading run C:\Data\ccRCC\CCRCC_Discovery_20\mzml\CPTAC_CCRCC_W_JHU_20190112_LUMOS_C3L-00010_T.mzML
[315:47] 5621709 library precursors are potentially detectable
[315:48] 5856 spectra added to the library
[315:48] Loading run C:\Data\ccRCC\CCRCC_Discovery_20\mzml\CPTAC_CCRCC_W_JHU_20190112_LUMOS_C3L-00011_NAT.mzML
[315:58] 5621709 library precursors are potentially detectable
[315:59] 3381 spectra added to the library
[316:00] Loading run C:\Data\ccRCC\CCRCC_Discovery_20\mzml\CPTAC_CCRCC_W_JHU_20190112_LUMOS_C3L-00011_T.mzML
[316:09] 5621709 library precursors are potentially detectable
[316:10] 11920 spectra added to the library
[316:11] Loading run C:\Data\ccRCC\CCRCC_Discovery_20\mzml\CPTAC_CCRCC_W_JHU_20190112_LUMOS_C3L-00026_NAT.mzML
[316:21] 5621709 library precursors are potentially detectable
[316:22] 7618 spectra added to the library
[316:22] Loading run C:\Data\ccRCC\CCRCC_Discovery_20\mzml\CPTAC_CCRCC_W_JHU_20190112_LUMOS_C3L-00026_T.mzML
[316:31] 5621709 library precursors are potentially detectable
[316:32] 2319 spectra added to the library
[316:33] Loading run C:\Data\ccRCC\CCRCC_Discovery_20\mzml\CPTAC_CCRCC_W_JHU_20190112_LUMOS_C3L-00079_NAT.mzML
[316:42] 5621709 library precursors are potentially detectable
[316:43] 7675 spectra added to the library
[316:44] Loading run C:\Data\ccRCC\CCRCC_Discovery_20\mzml\CPTAC_CCRCC_W_JHU_20190112_LUMOS_C3L-00079_T.mzML
[316:53] 5621709 library precursors are potentially detectable
[316:54] 7068 spectra added to the library
[316:55] Loading run C:\Data\ccRCC\CCRCC_Discovery_20\mzml\CPTAC_CCRCC_W_JHU_20190112_LUMOS_C3L-00088_NAT.mzML
[317:05] 5621709 library precursors are potentially detectable
[317:06] 6292 spectra added to the library
[317:06] Loading run C:\Data\ccRCC\CCRCC_Discovery_20\mzml\CPTAC_CCRCC_W_JHU_20190112_LUMOS_C3L-00088_T.mzML
[317:16] 5621709 library precursors are potentially detectable
[317:17] 2827 spectra added to the library
[317:17] Loading run C:\Data\ccRCC\CCRCC_Discovery_20\mzml\CPTAC_CCRCC_W_JHU_20190112_LUMOS_C3L-00096_NAT.mzML
[317:27] 5621709 library precursors are potentially detectable
[317:28] 4253 spectra added to the library
[317:28] Loading run C:\Data\ccRCC\CCRCC_Discovery_20\mzml\CPTAC_CCRCC_W_JHU_20190112_LUMOS_C3L-00096_T.mzML
[317:38] 5621709 library precursors are potentially detectable
[317:39] 7272 spectra added to the library
[317:40] Loading run C:\Data\ccRCC\CCRCC_Discovery_20\mzml\CPTAC_CCRCC_W_JHU_20190112_LUMOS_C3L-00097_NAT.mzML
[317:50] 5621709 library precursors are potentially detectable
[317:50] 3948 spectra added to the library
[317:51] Loading run C:\Data\ccRCC\CCRCC_Discovery_20\mzml\CPTAC_CCRCC_W_JHU_20190112_LUMOS_C3L-00097_T.mzML
[318:00] 5621709 library precursors are potentially detectable
[318:01] 1105 spectra added to the library
[318:02] Loading run C:\Data\ccRCC\CCRCC_Discovery_20\mzml\CPTAC_CCRCC_W_JHU_20190112_LUMOS_C3L-00103_NAT.mzML
[318:11] 5621709 library precursors are potentially detectable
[318:12] 7860 spectra added to the library
[318:13] Loading run C:\Data\ccRCC\CCRCC_Discovery_20\mzml\CPTAC_CCRCC_W_JHU_20190112_LUMOS_C3L-00103_T.mzML
[318:23] 5621709 library precursors are potentially detectable
[318:24] 4943 spectra added to the library
[318:24] Loading run C:\Data\ccRCC\CCRCC_Discovery_20\mzml\CPTAC_CCRCC_W_JHU_20190112_LUMOS_C3L-00183_NAT.mzML
[318:34] 5621709 library precursors are potentially detectable
[318:35] 2395 spectra added to the library
[318:35] Loading run C:\Data\ccRCC\CCRCC_Discovery_20\mzml\CPTAC_CCRCC_W_JHU_20190112_LUMOS_C3L-00183_T.mzML
[318:44] 5621709 library precursors are potentially detectable
[318:45] 2764 spectra added to the library
[318:46] Saving spectral library to C:\Data\ccRCC\CCRCC_Discovery_20\DIANN-results_max3\report-lib.tsv
[319:02] 147797 precursors saved
[319:02] Loading the generated library and saving it in the .speclib format
[319:02] Loading spectral library C:\Data\ccRCC\CCRCC_Discovery_20\DIANN-results_max3\report-lib.tsv
[319:08] Spectral library loaded: 11333 protein isoforms, 12150 protein groups and 147797 precursors in 111308 elution groups.
[319:08] Loading protein annotations from FASTA C:\Data\PXD022992_DIA_Melanoma\2021-03-23-reviewed-contam-UP000005640.fas
[319:09] Protein names missing for some isoforms
[319:09] Gene names missing for some isoforms
[319:09] Library contains 11332 proteins, and 11296 genes
[319:09] Saving the library to C:\Data\ccRCC\CCRCC_Discovery_20\DIANN-results_max3\report-lib.tsv.speclib

[319:13] Second pass: using the newly created spectral library to reanalyse the data
[319:13] File #1/20
[319:13] Loading run C:\Data\ccRCC\CCRCC_Discovery_20\mzml\CPTAC_CCRCC_W_JHU_20190112_LUMOS_C3L-00004_NAT.mzML
[319:23] 147797 library precursors are potentially detectable
[319:23] Processing...
[319:26] RT window set to 2.88273
[319:26] Recommended MS1 mass accuracy setting: 8.97851 ppm
[319:36] Removing low confidence identifications
[319:36] Searching PTM decoys
[319:36] Removing interfering precursors
[319:39] Training neural networks: 131601 targets, 135532 decoys
[319:46] Number of IDs at 0.01 FDR: 96550
[319:47] Calculating protein q-values
[319:47] Number of genes identified at 1% FDR: 8062 (precursor-level), 7268 (protein-level) (inference performed using proteotypic peptides only)
[319:47] Quantification
[319:47] Precursors with monitored PTMs at 1% FDR: 1060 out of 1169
[319:47] Unmodified precursors with monitored PTM sites at 1% FDR: 247 out of 271

[319:48] File #2/20
[319:48] Loading run C:\Data\ccRCC\CCRCC_Discovery_20\mzml\CPTAC_CCRCC_W_JHU_20190112_LUMOS_C3L-00004_T.mzML
[319:58] 147797 library precursors are potentially detectable
[319:58] Processing...
[320:01] RT window set to 2.9027
[320:01] Recommended MS1 mass accuracy setting: 8.86057 ppm
[320:10] Removing low confidence identifications
[320:10] Searching PTM decoys
[320:10] Removing interfering precursors
[320:13] Training neural networks: 134003 targets, 136306 decoys
[320:20] Number of IDs at 0.01 FDR: 107392
[320:21] Calculating protein q-values
[320:21] Number of genes identified at 1% FDR: 8419 (precursor-level), 7598 (protein-level) (inference performed using proteotypic peptides only)
[320:21] Quantification
[320:22] Precursors with monitored PTMs at 1% FDR: 1253 out of 1310
[320:22] Unmodified precursors with monitored PTM sites at 1% FDR: 282 out of 294

[320:22] File #3/20
[320:22] Loading run C:\Data\ccRCC\CCRCC_Discovery_20\mzml\CPTAC_CCRCC_W_JHU_20190112_LUMOS_C3L-00010_NAT.mzML
[320:32] 147797 library precursors are potentially detectable
[320:32] Processing...
[320:36] RT window set to 2.85718
[320:36] Recommended MS1 mass accuracy setting: 8.35572 ppm
[320:45] Removing low confidence identifications
[320:45] Searching PTM decoys
[320:45] Removing interfering precursors
[320:48] Training neural networks: 132146 targets, 135188 decoys
[320:55] Number of IDs at 0.01 FDR: 101183
[320:56] Calculating protein q-values
[320:56] Number of genes identified at 1% FDR: 8138 (precursor-level), 7376 (protein-level) (inference performed using proteotypic peptides only)
[320:56] Quantification
[320:57] Precursors with monitored PTMs at 1% FDR: 1122 out of 1232
[320:57] Unmodified precursors with monitored PTM sites at 1% FDR: 262 out of 295

[320:58] File #4/20
[320:58] Loading run C:\Data\ccRCC\CCRCC_Discovery_20\mzml\CPTAC_CCRCC_W_JHU_20190112_LUMOS_C3L-00010_T.mzML
[321:07] 147797 library precursors are potentially detectable
[321:07] Processing...
[321:11] RT window set to 2.86177
[321:11] Recommended MS1 mass accuracy setting: 9.94355 ppm
[321:20] Removing low confidence identifications
[321:20] Searching PTM decoys
[321:20] Removing interfering precursors
[321:23] Training neural networks: 129030 targets, 134006 decoys
[321:30] Number of IDs at 0.01 FDR: 88734
[321:31] Calculating protein q-values
[321:31] Number of genes identified at 1% FDR: 7656 (precursor-level), 7056 (protein-level) (inference performed using proteotypic peptides only)
[321:31] Quantification
[321:31] Precursors with monitored PTMs at 1% FDR: 961 out of 1079
[321:31] Unmodified precursors with monitored PTM sites at 1% FDR: 225 out of 254

[321:32] File #5/20
[321:32] Loading run C:\Data\ccRCC\CCRCC_Discovery_20\mzml\CPTAC_CCRCC_W_JHU_20190112_LUMOS_C3L-00011_NAT.mzML
[321:41] 147797 library precursors are potentially detectable
[321:41] Processing...
[321:45] RT window set to 2.88423
[321:45] Recommended MS1 mass accuracy setting: 8.25733 ppm
[321:54] Removing low confidence identifications
[321:54] Searching PTM decoys
[321:54] Removing interfering precursors
[321:57] Training neural networks: 132854 targets, 135224 decoys
[322:04] Number of IDs at 0.01 FDR: 103650
[322:05] Calculating protein q-values
[322:05] Number of genes identified at 1% FDR: 8379 (precursor-level), 7559 (protein-level) (inference performed using proteotypic peptides only)
[322:05] Quantification
[322:06] Precursors with monitored PTMs at 1% FDR: 1205 out of 1298
[322:06] Unmodified precursors with monitored PTM sites at 1% FDR: 280 out of 301

[322:07] File #6/20
[322:07] Loading run C:\Data\ccRCC\CCRCC_Discovery_20\mzml\CPTAC_CCRCC_W_JHU_20190112_LUMOS_C3L-00011_T.mzML
[322:16] 147797 library precursors are potentially detectable
[322:16] Processing...
[322:18] RT window set to 2.89515
[322:18] Recommended MS1 mass accuracy setting: 7.41229 ppm
[322:27] Removing low confidence identifications
[322:27] Searching PTM decoys
[322:27] Removing interfering precursors
[322:30] Training neural networks: 131446 targets, 133480 decoys
[322:37] Number of IDs at 0.01 FDR: 102514
[322:38] Calculating protein q-values
[322:38] Number of genes identified at 1% FDR: 8231 (precursor-level), 7485 (protein-level) (inference performed using proteotypic peptides only)
[322:38] Quantification
[322:39] Precursors with monitored PTMs at 1% FDR: 1196 out of 1256
[322:39] Unmodified precursors with monitored PTM sites at 1% FDR: 270 out of 286

[322:39] File #7/20
[322:39] Loading run C:\Data\ccRCC\CCRCC_Discovery_20\mzml\CPTAC_CCRCC_W_JHU_20190112_LUMOS_C3L-00026_NAT.mzML
[322:48] 147797 library precursors are potentially detectable
[322:48] Processing...
[322:52] RT window set to 2.83811
[322:52] Recommended MS1 mass accuracy setting: 8.15751 ppm
[323:01] Removing low confidence identifications
[323:01] Searching PTM decoys
[323:01] Removing interfering precursors
[323:04] Training neural networks: 132259 targets, 133699 decoys
[323:11] Number of IDs at 0.01 FDR: 103585
[323:12] Calculating protein q-values
[323:12] Number of genes identified at 1% FDR: 8466 (precursor-level), 7608 (protein-level) (inference performed using proteotypic peptides only)
[323:12] Quantification
[323:13] Precursors with monitored PTMs at 1% FDR: 1139 out of 1270
[323:13] Unmodified precursors with monitored PTM sites at 1% FDR: 267 out of 295

[323:13] File #8/20
[323:13] Loading run C:\Data\ccRCC\CCRCC_Discovery_20\mzml\CPTAC_CCRCC_W_JHU_20190112_LUMOS_C3L-00026_T.mzML
[323:22] 147797 library precursors are potentially detectable
[323:22] Processing...
[323:25] RT window set to 2.83695
[323:25] Recommended MS1 mass accuracy setting: 9.33766 ppm
[323:34] Removing low confidence identifications
[323:34] Searching PTM decoys
[323:34] Removing interfering precursors
[323:37] Training neural networks: 123782 targets, 129771 decoys
[323:44] Number of IDs at 0.01 FDR: 77618
[323:44] Calculating protein q-values
[323:44] Number of genes identified at 1% FDR: 7351 (precursor-level), 6485 (protein-level) (inference performed using proteotypic peptides only)
[323:44] Quantification
[323:45] Precursors with monitored PTMs at 1% FDR: 1009 out of 1123
[323:45] Unmodified precursors with monitored PTM sites at 1% FDR: 223 out of 254

[323:46] File #9/20
[323:46] Loading run C:\Data\ccRCC\CCRCC_Discovery_20\mzml\CPTAC_CCRCC_W_JHU_20190112_LUMOS_C3L-00079_NAT.mzML
[323:55] 147797 library precursors are potentially detectable
[323:55] Processing...
[323:56] RT window set to 2.86882
[323:56] Recommended MS1 mass accuracy setting: 7.46795 ppm
[324:05] Removing low confidence identifications
[324:05] Searching PTM decoys
[324:05] Removing interfering precursors
[324:08] Training neural networks: 134450 targets, 134618 decoys
[324:15] Number of IDs at 0.01 FDR: 111350
[324:16] Calculating protein q-values
[324:16] Number of genes identified at 1% FDR: 8740 (precursor-level), 7972 (protein-level) (inference performed using proteotypic peptides only)
[324:16] Quantification
[324:17] Precursors with monitored PTMs at 1% FDR: 1306 out of 1352
[324:17] Unmodified precursors with monitored PTM sites at 1% FDR: 287 out of 304

[324:17] File #10/20
[324:17] Loading run C:\Data\ccRCC\CCRCC_Discovery_20\mzml\CPTAC_CCRCC_W_JHU_20190112_LUMOS_C3L-00079_T.mzML
[324:27] 147797 library precursors are potentially detectable
[324:27] Processing...
[324:29] RT window set to 2.90805
[324:29] Recommended MS1 mass accuracy setting: 8.22632 ppm
[324:38] Removing low confidence identifications
[324:38] Searching PTM decoys
[324:38] Removing interfering precursors
[324:41] Training neural networks: 133469 targets, 134051 decoys
[324:48] Number of IDs at 0.01 FDR: 106983
[324:49] Calculating protein q-values
[324:49] Number of genes identified at 1% FDR: 8517 (precursor-level), 7798 (protein-level) (inference performed using proteotypic peptides only)
[324:49] Quantification
[324:50] Precursors with monitored PTMs at 1% FDR: 1211 out of 1286
[324:50] Unmodified precursors with monitored PTM sites at 1% FDR: 275 out of 290

[324:50] File #11/20
[324:50] Loading run C:\Data\ccRCC\CCRCC_Discovery_20\mzml\CPTAC_CCRCC_W_JHU_20190112_LUMOS_C3L-00088_NAT.mzML
[324:59] 147797 library precursors are potentially detectable
[324:59] Processing...
[325:02] RT window set to 2.89151
[325:02] Recommended MS1 mass accuracy setting: 7.85861 ppm
[325:11] Removing low confidence identifications
[325:11] Searching PTM decoys
[325:11] Removing interfering precursors
[325:14] Training neural networks: 132849 targets, 134894 decoys
[325:21] Number of IDs at 0.01 FDR: 103977
[325:22] Calculating protein q-values
[325:22] Number of genes identified at 1% FDR: 8453 (precursor-level), 7594 (protein-level) (inference performed using proteotypic peptides only)
[325:22] Quantification
[325:23] Precursors with monitored PTMs at 1% FDR: 1196 out of 1292
[325:23] Unmodified precursors with monitored PTM sites at 1% FDR: 274 out of 295

[325:23] File #12/20
[325:23] Loading run C:\Data\ccRCC\CCRCC_Discovery_20\mzml\CPTAC_CCRCC_W_JHU_20190112_LUMOS_C3L-00088_T.mzML
[325:32] 147797 library precursors are potentially detectable
[325:32] Processing...
[325:36] RT window set to 2.8617
[325:36] Recommended MS1 mass accuracy setting: 8.60112 ppm
[325:45] Removing low confidence identifications
[325:45] Searching PTM decoys
[325:45] Removing interfering precursors
[325:48] Training neural networks: 128622 targets, 132163 decoys
[325:55] Number of IDs at 0.01 FDR: 89348
[325:56] Calculating protein q-values
[325:56] Number of genes identified at 1% FDR: 7826 (precursor-level), 7093 (protein-level) (inference performed using proteotypic peptides only)
[325:56] Quantification
[325:56] Precursors with monitored PTMs at 1% FDR: 1027 out of 1170
[325:56] Unmodified precursors with monitored PTM sites at 1% FDR: 219 out of 257

[325:57] File #13/20
[325:57] Loading run C:\Data\ccRCC\CCRCC_Discovery_20\mzml\CPTAC_CCRCC_W_JHU_20190112_LUMOS_C3L-00096_NAT.mzML
[326:06] 147797 library precursors are potentially detectable
[326:06] Processing...
[326:08] RT window set to 2.8901
[326:08] Recommended MS1 mass accuracy setting: 7.8035 ppm
[326:17] Removing low confidence identifications
[326:17] Searching PTM decoys
[326:17] Removing interfering precursors
[326:20] Training neural networks: 133601 targets, 134484 decoys
[326:27] Number of IDs at 0.01 FDR: 108913
[326:28] Calculating protein q-values
[326:28] Number of genes identified at 1% FDR: 8665 (precursor-level), 7773 (protein-level) (inference performed using proteotypic peptides only)
[326:28] Quantification
[326:29] Precursors with monitored PTMs at 1% FDR: 1256 out of 1333
[326:29] Unmodified precursors with monitored PTM sites at 1% FDR: 285 out of 309

[326:29] File #14/20
[326:29] Loading run C:\Data\ccRCC\CCRCC_Discovery_20\mzml\CPTAC_CCRCC_W_JHU_20190112_LUMOS_C3L-00096_T.mzML
[326:39] 147797 library precursors are potentially detectable
[326:39] Processing...
[326:41] RT window set to 2.89688
[326:41] Recommended MS1 mass accuracy setting: 7.66417 ppm
[326:50] Removing low confidence identifications
[326:50] Searching PTM decoys
[326:50] Removing interfering precursors
[326:53] Training neural networks: 132769 targets, 134570 decoys
[327:00] Number of IDs at 0.01 FDR: 105142
[327:01] Calculating protein q-values
[327:01] Number of genes identified at 1% FDR: 8444 (precursor-level), 7589 (protein-level) (inference performed using proteotypic peptides only)
[327:01] Quantification
[327:02] Precursors with monitored PTMs at 1% FDR: 1223 out of 1303
[327:02] Unmodified precursors with monitored PTM sites at 1% FDR: 261 out of 291

[327:02] File #15/20
[327:02] Loading run C:\Data\ccRCC\CCRCC_Discovery_20\mzml\CPTAC_CCRCC_W_JHU_20190112_LUMOS_C3L-00097_NAT.mzML
[327:11] 147797 library precursors are potentially detectable
[327:12] Processing...
[327:15] RT window set to 2.88988
[327:15] Recommended MS1 mass accuracy setting: 8.76333 ppm
[327:24] Removing low confidence identifications
[327:24] Searching PTM decoys
[327:24] Removing interfering precursors
[327:27] Training neural networks: 130556 targets, 133733 decoys
[327:34] Number of IDs at 0.01 FDR: 97422
[327:35] Calculating protein q-values
[327:35] Number of genes identified at 1% FDR: 8124 (precursor-level), 7305 (protein-level) (inference performed using proteotypic peptides only)
[327:35] Quantification
[327:35] Precursors with monitored PTMs at 1% FDR: 1147 out of 1253
[327:35] Unmodified precursors with monitored PTM sites at 1% FDR: 249 out of 277

[327:36] File #16/20
[327:36] Loading run C:\Data\ccRCC\CCRCC_Discovery_20\mzml\CPTAC_CCRCC_W_JHU_20190112_LUMOS_C3L-00097_T.mzML
[327:45] 147797 library precursors are potentially detectable
[327:45] Processing...
[327:48] RT window set to 2.88845
[327:48] Recommended MS1 mass accuracy setting: 9.06356 ppm
[327:56] Removing low confidence identifications
[327:56] Searching PTM decoys
[327:56] Removing interfering precursors
[327:59] Training neural networks: 125545 targets, 129446 decoys
[328:06] Number of IDs at 0.01 FDR: 83168
[328:07] Calculating protein q-values
[328:07] Number of genes identified at 1% FDR: 7420 (precursor-level), 6593 (protein-level) (inference performed using proteotypic peptides only)
[328:07] Quantification
[328:07] Precursors with monitored PTMs at 1% FDR: 986 out of 1119
[328:07] Unmodified precursors with monitored PTM sites at 1% FDR: 211 out of 236

[328:08] File #17/20
[328:08] Loading run C:\Data\ccRCC\CCRCC_Discovery_20\mzml\CPTAC_CCRCC_W_JHU_20190112_LUMOS_C3L-00103_NAT.mzML
[328:17] 147797 library precursors are potentially detectable
[328:17] Processing...
[328:19] RT window set to 2.88246
[328:19] Recommended MS1 mass accuracy setting: 7.22325 ppm
[328:28] Removing low confidence identifications
[328:28] Searching PTM decoys
[328:28] Removing interfering precursors
[328:31] Training neural networks: 133683 targets, 134151 decoys
[328:39] Number of IDs at 0.01 FDR: 109165
[328:40] Calculating protein q-values
[328:40] Number of genes identified at 1% FDR: 8605 (precursor-level), 7828 (protein-level) (inference performed using proteotypic peptides only)
[328:40] Quantification
[328:41] Precursors with monitored PTMs at 1% FDR: 1366 out of 1408
[328:41] Unmodified precursors with monitored PTM sites at 1% FDR: 300 out of 311

[328:41] File #18/20
[328:41] Loading run C:\Data\ccRCC\CCRCC_Discovery_20\mzml\CPTAC_CCRCC_W_JHU_20190112_LUMOS_C3L-00103_T.mzML
[328:50] 147797 library precursors are potentially detectable
[328:50] Processing...
[328:54] RT window set to 2.88131
[328:54] Recommended MS1 mass accuracy setting: 9.0788 ppm
[329:02] Removing low confidence identifications
[329:02] Searching PTM decoys
[329:02] Removing interfering precursors
[329:05] Training neural networks: 126697 targets, 130416 decoys
[329:12] Number of IDs at 0.01 FDR: 86120
[329:12] Calculating protein q-values
[329:12] Number of genes identified at 1% FDR: 7669 (precursor-level), 6889 (protein-level) (inference performed using proteotypic peptides only)
[329:12] Quantification
[329:13] Precursors with monitored PTMs at 1% FDR: 991 out of 1092
[329:13] Unmodified precursors with monitored PTM sites at 1% FDR: 208 out of 231

[329:14] File #19/20
[329:14] Loading run C:\Data\ccRCC\CCRCC_Discovery_20\mzml\CPTAC_CCRCC_W_JHU_20190112_LUMOS_C3L-00183_NAT.mzML
[329:23] 147797 library precursors are potentially detectable
[329:23] Processing...
[329:26] RT window set to 2.89523
[329:26] Recommended MS1 mass accuracy setting: 7.50772 ppm
[329:35] Removing low confidence identifications
[329:35] Searching PTM decoys
[329:35] Removing interfering precursors
[329:38] Training neural networks: 129041 targets, 132602 decoys
[329:45] Number of IDs at 0.01 FDR: 93838
[329:46] Calculating protein q-values
[329:46] Number of genes identified at 1% FDR: 7873 (precursor-level), 7160 (protein-level) (inference performed using proteotypic peptides only)
[329:46] Quantification
[329:46] Precursors with monitored PTMs at 1% FDR: 1070 out of 1156
[329:46] Unmodified precursors with monitored PTM sites at 1% FDR: 251 out of 277

[329:47] File #20/20
[329:47] Loading run C:\Data\ccRCC\CCRCC_Discovery_20\mzml\CPTAC_CCRCC_W_JHU_20190112_LUMOS_C3L-00183_T.mzML
[329:56] 147797 library precursors are potentially detectable
[329:56] Processing...
[329:59] RT window set to 2.86205
[329:59] Recommended MS1 mass accuracy setting: 8.18279 ppm
[330:06] Removing low confidence identifications
[330:06] Searching PTM decoys
[330:07] Removing interfering precursors
[330:09] Training neural networks: 115982 targets, 123525 decoys
[330:16] Number of IDs at 0.01 FDR: 64770
[330:16] Calculating protein q-values
[330:16] Number of genes identified at 1% FDR: 6700 (precursor-level), 5936 (protein-level) (inference performed using proteotypic peptides only)
[330:16] Quantification
[330:17] Precursors with monitored PTMs at 1% FDR: 842 out of 990
[330:17] Unmodified precursors with monitored PTM sites at 1% FDR: 160 out of 191

[330:17] Cross-run analysis
[330:17] Reading quantification information: 20 files
[330:22] Quantifying peptides
[330:35] Quantifying proteins
[330:37] Calculating q-values for protein and gene groups
[330:38] Calculating global q-values for protein and gene groups
[330:38] Writing report
[332:05] Report saved to C:\Data\ccRCC\CCRCC_Discovery_20\DIANN-results_max3\report.tsv.
[332:05] Saving precursor levels matrix
[332:07] Precursor levels matrix (1% precursor and protein group FDR) saved to C:\Data\ccRCC\CCRCC_Discovery_20\DIANN-results_max3\report.pr_matrix.tsv.
[332:07] Saving protein group levels matrix
[332:07] Protein group levels matrix (1% precursor FDR and protein group FDR) saved to C:\Data\ccRCC\CCRCC_Discovery_20\DIANN-results_max3\report.pg_matrix.tsv.
[332:07] Saving gene group levels matrix
[332:07] Gene groups levels matrix (1% precursor FDR and protein group FDR) saved to C:\Data\ccRCC\CCRCC_Discovery_20\DIANN-results_max3\report.gg_matrix.tsv.
[332:07] Saving unique genes levels matrix
[332:07] Unique genes levels matrix (1% precursor FDR and protein group FDR) saved to C:\Data\ccRCC\CCRCC_Discovery_20\DIANN-results_max3\report.unique_genes_matrix.tsv.
[332:07] Stats report saved to C:\Data\ccRCC\CCRCC_Discovery_20\DIANN-results_max3\report.stats.tsv
[332:07] Log saved to C:\Data\ccRCC\CCRCC_Discovery_20\DIANN-results_max3\report.log.txt
Finished


DIA-NN exited
DIA-NN-plotter.exe "C:\Data\ccRCC\CCRCC_Discovery_20\DIANN-results_max3\report.stats.tsv" "C:\Data\ccRCC\CCRCC_Discovery_20\DIANN-results_max3\report.tsv" "C:\Data\ccRCC\CCRCC_Discovery_20\DIANN-results_max3\report.pdf"
PDF report will be generated in the background
