## Supplementary Data 1 for "One-stop analysis of DIA proteomics data using MSFragger-DIA and FragPipe computational platform": diann-output.log.txt

DIA-NN 1.8.1 (Data-Independent Acquisition by Neural Networks)
Compiled on Apr 15 2022 08:45:18
Current date and time: Sun Aug 21 01:07:52 2022
Logical CPU cores: 56
/storage/yufe/tools/FragPipe-18.0/fragpipe/tools/diann/1.8.1/linux/diann-1.8.1 --lib library.tsv --threads 56 --verbose 1 --out diann-output.tsv --qvalue 0.01 --matrix-spec-q 0.01 --matrices --no-prot-inf --smart-profiling --no-quant-files --peak-center --no-ifs-removal --predictor --dl-no-rt --dl-no-im --strip-unknown-mods --cfg /storage/yufe/results/msfragger_dia_paper/runtime_melanoma_phospho/dev02/diaumpire/filelist_diann.txt 

Thread number set to 56
Output will be filtered at 0.01 FDR
Protein x sample matrices will be also filtered using run-specific protein q-value
Precursor/protein x samples expression level matrices will be saved along with the main report
Protein inference will not be performed
When generating a spectral library, in silico predicted spectra will be retained if deemed more reliable than experimental ones
.quant files will not be saved to the disk
Fixed-width center of each elution peak will be used for quantification
Interference removal from fragment elution curves disabled
Deep learning will be used to generate a new in silico spectral library from peptides provided
RTs will not be predicted using deep learning
IMs will not be predicted using deep learning
DIA-NN will use deep learning to predict spectra/RTs/IMs even for peptides carrying modifications which are not recognised by the deep learning predictor. In this scenario, if also generating a spectral library from the DIA data or using the MBR mode, it might or might not be better (depends on the data) to also use the --out-measured-rt option - it's recommended to test it with and without this option
DIA-NN will optimise the mass accuracy automatically using the first run in the experiment. This is useful primarily for quick initial analyses, when it is not yet known which mass accuracy setting works best for a particular acquisition scheme.

12 files will be processed
[0:00] Loading spectral library library.tsv
[0:00] Finding proteotypic peptides (assuming that the list of UniProt ids provided for each peptide is complete)
[0:00] Spectral library loaded: 130 protein isoforms, 130 protein groups and 208 precursors in 196 elution groups.
[0:02] [0:02] [0:03] [0:03] Saving the library to lib.predicted.speclib
[0:03] Initialising library

[0:03] File #1/12
[0:03] Loading run /storage/yufe/data/PXD022992/mzml/phos/201905024_S_SK_phos_2.mzML
[0:59] 208 library precursors are potentially detectable
[0:59] Processing...
[0:59] RT window set to 2.43803
[0:59] Peak width: 3.11613
[0:59] Scan window radius set to 6
[0:59] Recommended MS1 mass accuracy setting: 5.36651 ppm
[0:59] Optimised mass accuracy: 5.086 ppm
[0:59] Removing low confidence identifications
[0:59] Removing interfering precursors
[0:59] Training neural networks: 195 targets, 107 decoys
[0:59] Number of IDs at 0.01 FDR: 183
[0:59] Calculating protein q-values
[0:59] Number of protein isoforms identified at 1% FDR: 122 (precursor-level), 37 (protein-level) (inference performed using proteotypic peptides only)
[0:59] Quantification

[0:59] File #2/12
[0:59] Loading run /storage/yufe/data/PXD022992/mzml/phos/201905024_S_A375_phos_2.mzML
[1:52] 208 library precursors are potentially detectable
[1:52] Processing...
[1:52] RT window set to 2.57973
[1:52] Recommended MS1 mass accuracy setting: 2.27798 ppm
[1:52] Removing low confidence identifications
[1:52] Removing interfering precursors
[1:52] Training neural networks: 187 targets, 101 decoys
[1:53] Number of IDs at 0.01 FDR: 177
[1:53] Calculating protein q-values
[1:53] Number of protein isoforms identified at 1% FDR: 117 (precursor-level), 114 (protein-level) (inference performed using proteotypic peptides only)
[1:53] Quantification

[1:53] File #3/12
[1:53] Loading run /storage/yufe/data/PXD022992/mzml/phos/201905024_S_A375_phos_1.mzML
[2:48] 208 library precursors are potentially detectable
[2:48] Processing...
[2:48] RT window set to 2.41962
[2:48] Recommended MS1 mass accuracy setting: 3.0853 ppm
[2:48] Removing low confidence identifications
[2:48] Removing interfering precursors
[2:48] Too few confident identifications, neural networks will not be used
[2:48] Number of IDs at 0.01 FDR: 164
[2:48] Number of IDs at 0.01 FDR: 163
[2:48] Calculating protein q-values
[2:48] Number of protein isoforms identified at 1% FDR: 111 (precursor-level), 35 (protein-level) (inference performed using proteotypic peptides only)
[2:48] Quantification

[2:48] File #4/12
[2:48] Loading run /storage/yufe/data/PXD022992/mzml/phos/201905024_S_G361_phos_2.mzML
[3:44] 208 library precursors are potentially detectable
[3:44] Processing...
[3:45] RT window set to 2.6015
[3:45] Recommended MS1 mass accuracy setting: 2.74726 ppm
[3:45] Removing low confidence identifications
[3:45] Removing interfering precursors
[3:45] Training neural networks: 193 targets, 101 decoys
[3:45] Number of IDs at 0.01 FDR: 188
[3:45] Calculating protein q-values
[3:45] Number of protein isoforms identified at 1% FDR: 124 (precursor-level), 0 (protein-level) (inference performed using proteotypic peptides only)
[3:45] Quantification

[3:45] File #5/12
[3:45] Loading run /storage/yufe/data/PXD022992/mzml/phos/201905024_F_SH4_phos_1.mzML
[4:39] 208 library precursors are potentially detectable
[4:39] Processing...
[4:39] RT window set to 3.15967
[4:39] Recommended MS1 mass accuracy setting: 1.5682 ppm
[4:39] Removing low confidence identifications
[4:39] Removing interfering precursors
[4:39] Training neural networks: 197 targets, 109 decoys
[4:39] Number of IDs at 0.01 FDR: 185
[4:39] Calculating protein q-values
[4:39] Number of protein isoforms identified at 1% FDR: 124 (precursor-level), 0 (protein-level) (inference performed using proteotypic peptides only)
[4:39] Quantification

[4:39] File #6/12
[4:39] Loading run /storage/yufe/data/PXD022992/mzml/phos/201905024_F_SH4_phos_2.mzML
[5:38] 208 library precursors are potentially detectable
[5:38] Processing...
[5:38] RT window set to 2.41765
[5:38] Recommended MS1 mass accuracy setting: 2.01325 ppm
[5:38] Removing low confidence identifications
[5:38] Removing interfering precursors
[5:38] Too few confident identifications, neural networks will not be used
[5:38] Number of IDs at 0.01 FDR: 151
[5:38] Number of IDs at 0.01 FDR: 166
[5:38] Calculating protein q-values
[5:38] Number of protein isoforms identified at 1% FDR: 113 (precursor-level), 113 (protein-level) (inference performed using proteotypic peptides only)
[5:38] Quantification

[5:38] File #7/12
[5:38] Loading run /storage/yufe/data/PXD022992/mzml/phos/201905024_F_7951_phos_2.mzML
[6:31] 208 library precursors are potentially detectable
[6:31] Processing...
[6:31] RT window set to 2.28921
[6:31] Recommended MS1 mass accuracy setting: 1.35127 ppm
[6:31] Removing low confidence identifications
[6:31] Removing interfering precursors
[6:31] Too few confident identifications, neural networks will not be used
[6:31] Number of IDs at 0.01 FDR: 165
[6:31] Number of IDs at 0.01 FDR: 160
[6:31] Calculating protein q-values
[6:31] Number of protein isoforms identified at 1% FDR: 107 (precursor-level), 107 (protein-level) (inference performed using proteotypic peptides only)
[6:31] Quantification

[6:31] File #8/12
[6:31] Loading run /storage/yufe/data/PXD022992/mzml/phos/201905024_F_HTB69_phos_2.mzML
[7:28] 208 library precursors are potentially detectable
[7:28] Processing...
[7:28] RT window set to 2.64008
[7:28] Recommended MS1 mass accuracy setting: 3.03348 ppm
[7:28] Removing low confidence identifications
[7:28] Removing interfering precursors
[7:28] Training neural networks: 197 targets, 103 decoys
[7:28] Number of IDs at 0.01 FDR: 194
[7:28] Calculating protein q-values
[7:28] Number of protein isoforms identified at 1% FDR: 127 (precursor-level), 0 (protein-level) (inference performed using proteotypic peptides only)
[7:28] Quantification

[7:28] File #9/12
[7:28] Loading run /storage/yufe/data/PXD022992/mzml/phos/201905024_S_G361_phoso_1.mzML
[8:23] 208 library precursors are potentially detectable
[8:23] Processing...
[8:23] RT window set to 2.72779
[8:23] Recommended MS1 mass accuracy setting: 2.43431 ppm
[8:23] Removing low confidence identifications
[8:23] Removing interfering precursors
[8:23] Too few confident identifications, neural networks will not be used
[8:23] Number of IDs at 0.01 FDR: 164
[8:23] Calculating protein q-values
[8:23] Number of protein isoforms identified at 1% FDR: 115 (precursor-level), 89 (protein-level) (inference performed using proteotypic peptides only)
[8:23] Quantification

[8:23] File #10/12
[8:23] Loading run /storage/yufe/data/PXD022992/mzml/phos/201905024_S_SK_phos_1.mzML
[9:23] 208 library precursors are potentially detectable
[9:23] Processing...
[9:23] RT window set to 2.30788
[9:23] Cannot perform MS1 mass calibration, too few confidently identified precursors
[9:23] Recommended MS1 mass accuracy setting: 2.20645 ppm
[9:23] Removing low confidence identifications
[9:23] Removing interfering precursors
[9:23] Too few confident identifications, neural networks will not be used
[9:23] Number of IDs at 0.01 FDR: 157
[9:23] Calculating protein q-values
[9:23] Number of protein isoforms identified at 1% FDR: 113 (precursor-level), 99 (protein-level) (inference performed using proteotypic peptides only)
[9:23] Quantification

[9:23] File #11/12
[9:23] Loading run /storage/yufe/data/PXD022992/mzml/phos/201905024_F_7951_phos_1.mzML
[10:17] 208 library precursors are potentially detectable
[10:17] Processing...
[10:18] RT window set to 2.17685
[10:18] Recommended MS1 mass accuracy setting: 2.7741 ppm
[10:18] Removing low confidence identifications
[10:18] Removing interfering precursors
[10:18] Too few confident identifications, neural networks will not be used
[10:18] Number of IDs at 0.01 FDR: 166
[10:18] Number of IDs at 0.01 FDR: 163
[10:18] Calculating protein q-values
[10:18] Number of protein isoforms identified at 1% FDR: 112 (precursor-level), 114 (protein-level) (inference performed using proteotypic peptides only)
[10:18] Quantification

[10:18] File #12/12
[10:18] Loading run /storage/yufe/data/PXD022992/mzml/phos/201905024_F_HTB69_phos_1.mzML
[11:11] 208 library precursors are potentially detectable
[11:11] Processing...
[11:11] RT window set to 2.29721
[11:11] Recommended MS1 mass accuracy setting: 3.48861 ppm
[11:11] Removing low confidence identifications
[11:11] Removing interfering precursors
[11:11] Too few confident identifications, neural networks will not be used
[11:11] Number of IDs at 0.01 FDR: 159
[11:11] Number of IDs at 0.01 FDR: 182
[11:11] Calculating protein q-values
[11:11] Number of protein isoforms identified at 1% FDR: 124 (precursor-level), 72 (protein-level) (inference performed using proteotypic peptides only)
[11:11] Quantification

[11:11] Cross-run analysis
[11:11] Reading quantification information: 12 files
[11:11] Quantifying peptides
[11:11] Quantifying proteins
[11:11] Calculating q-values for protein and gene groups
[11:11] Calculating global q-values for protein and gene groups
[11:11] Writing report
[11:11] Report saved to diann-output.tsv.
[11:11] Saving precursor levels matrix
[11:11] Precursor levels matrix (1% precursor and protein group FDR) saved to diann-output.pr_matrix.tsv.
[11:11] Saving protein group levels matrix
[11:11] Protein group levels matrix (1% precursor FDR and protein group FDR) saved to diann-output.pg_matrix.tsv.
[11:11] Saving gene group levels matrix
[11:11] Gene groups levels matrix (1% precursor FDR and protein group FDR) saved to diann-output.gg_matrix.tsv.
[11:11] Saving unique genes levels matrix
[11:11] Unique genes levels matrix (1% precursor FDR and protein group FDR) saved to diann-output.unique_genes_matrix.tsv.
[11:11] Stats report saved to diann-output.stats.tsv

Finished
