## Supplementary Data 2 for "One-stop analysis of DIA proteomics data using MSFragger-DIA and FragPipe computational platform": ccRCC_OmicsEV.html

Omics datasets evaluation report


### Omics datasets evaluation report

#### 2022-08-19

### 1 Introduction

In this evaluation, there are total **6** datasets. We used the evaluation metrics implemented in **OmicsEV** package to evaluate these datasets. The sample and class information for each dataset are shown in the table below.

| class | diann\_diann | diaumpire\_diann | diaumpiredda\_diann | msfraggerdia\_diann | msfraggerdiadda\_diann | TMT\_MD |
| --- | --- | --- | --- | --- | --- | --- |
| Normal | 79 | 79 | 79 | 79 | 79 | 79 |
| Tumor | 103 | 103 | 103 | 103 | 103 | 103 |

The detailed sample information is shown below.

| sample | class | batch | order |
| --- | --- | --- | --- |
| C3L-01287-N | Normal | 1 | 1 |
| C3L-00561-N | Normal | 1 | 2 |
| C3L-01603-N | Normal | 1 | 3 |
| C3N-00834-N | Normal | 1 | 4 |
| C3N-01214-N | Normal | 1 | 5 |
| C3N-01261-N | Normal | 1 | 6 |
| C3L-00917-N | Normal | 1 | 7 |
| C3L-00607-N | Normal | 1 | 8 |
| C3N-00194-N | Normal | 1 | 9 |
| C3L-00010-N | Normal | 1 | 10 |
| C3L-01861-N | Normal | 1 | 11 |
| C3N-01646-N | Normal | 1 | 12 |
| C3L-01281-N | Normal | 1 | 13 |
| C3N-00495-N | Normal | 1 | 14 |
| C3N-00831-N | Normal | 1 | 15 |
| C3L-00183-N | Normal | 1 | 16 |
| C3N-00168-N | Normal | 1 | 17 |
| C3L-00369-N | Normal | 1 | 18 |
| C3L-00791-N | Normal | 1 | 19 |
| C3L-00097-N | Normal | 1 | 20 |
| C3L-00004-N | Normal | 1 | 21 |
| C3N-00953-N | Normal | 1 | 22 |
| C3N-00150-N | Normal | 1 | 23 |
| C3N-00244-N | Normal | 1 | 24 |
| C3N-01178-N | Normal | 1 | 25 |
| C3L-00583-N | Normal | 1 | 26 |
| C3L-00088-N | Normal | 1 | 27 |
| C3L-00814-N | Normal | 1 | 28 |
| C3L-01885-N | Normal | 1 | 29 |
| C3L-00908-N | Normal | 1 | 30 |
| C3L-00026-N | Normal | 1 | 31 |
| C3L-00447-N | Normal | 1 | 32 |
| C3L-00416-N | Normal | 1 | 33 |
| C3N-00246-N | Normal | 1 | 34 |
| C3N-00148-N | Normal | 1 | 35 |
| C3N-00646-N | Normal | 1 | 36 |
| C3L-00902-N | Normal | 1 | 37 |
| C3L-00096-N | Normal | 1 | 38 |
| C3L-01836-N | Normal | 1 | 39 |
| C3N-00177-N | Normal | 1 | 40 |
| C3N-01200-N | Normal | 1 | 41 |
| C3L-00011-N | Normal | 1 | 42 |
| C3N-01808-N | Normal | 1 | 43 |
| C3N-01176-N | Normal | 1 | 44 |
| C3L-00103-N | Normal | 1 | 45 |
| C3N-01649-N | Normal | 1 | 46 |
| C3N-00149-N | Normal | 1 | 47 |
| C3L-00581-N | Normal | 1 | 48 |
| C3L-01313-N | Normal | 1 | 49 |
| C3N-00573-N | Normal | 1 | 50 |
| C3L-01302-N | Normal | 1 | 51 |
| C3N-01361-N | Normal | 1 | 52 |
| C3N-01651-N | Normal | 1 | 53 |
| C3L-00360-N | Normal | 1 | 54 |
| C3L-00448-N | Normal | 1 | 55 |
| C3L-00907-N | Normal | 1 | 56 |
| C3L-01286-N | Normal | 1 | 57 |
| C3L-01607-N | Normal | 1 | 58 |
| C3N-01179-N | Normal | 1 | 59 |
| C3L-00606-N | Normal | 1 | 60 |
| C3N-01648-N | Normal | 1 | 61 |
| C3N-00242-N | Normal | 1 | 62 |
| C3L-00418-N | Normal | 1 | 63 |
| C3N-00577-N | Normal | 1 | 64 |
| C3L-01882-N | Normal | 1 | 65 |
| C3N-00733-N | Normal | 1 | 66 |
| C3L-00910-N | Normal | 1 | 67 |
| C3L-00079-N | Normal | 1 | 68 |
| C3N-01522-N | Normal | 1 | 69 |
| C3N-01220-N | Normal | 1 | 70 |
| C3N-00852-N | Normal | 1 | 71 |
| C3N-00320-N | Normal | 1 | 72 |
| C3N-00437-N | Normal | 1 | 73 |
| C3N-00491-N | Normal | 1 | 74 |
| C3N-00317-N | Normal | 1 | 75 |
| C3N-00310-N | Normal | 1 | 76 |
| C3N-00390-N | Normal | 1 | 77 |
| C3N-00312-N | Normal | 1 | 78 |
| C3N-00494-N | Normal | 1 | 79 |
| C3N-01648-T | Tumor | 1 | 80 |
| C3N-01214-T | Tumor | 1 | 81 |
| C3N-00646-T | Tumor | 1 | 82 |
| C3L-00766-T | Tumor | 1 | 83 |
| C3L-00360-T | Tumor | 1 | 84 |
| C3L-00790-T | Tumor | 1 | 85 |
| C3N-00154-T | Tumor | 1 | 86 |
| C3L-00765-T | Tumor | 1 | 87 |
| C3N-00150-T | Tumor | 1 | 88 |
| C3L-01283-T | Tumor | 1 | 89 |
| C3N-00315-T | Tumor | 1 | 90 |
| C3N-00177-T | Tumor | 1 | 91 |
| C3L-00561-T | Tumor | 1 | 92 |
| C3L-00800-T | Tumor | 1 | 93 |
| C3L-01882-T | Tumor | 1 | 94 |
| C3N-00491-T | Tumor | 1 | 95 |
| C3N-00577-T | Tumor | 1 | 96 |
| C3L-00079-T | Tumor | 1 | 97 |
| C3N-01220-T | Tumor | 1 | 98 |
| C3L-01607-T | Tumor | 1 | 99 |
| C3N-00953-T | Tumor | 1 | 100 |
| C3N-00380-T | Tumor | 1 | 101 |
| C3N-00317-T | Tumor | 1 | 102 |
| C3N-00305-T | Tumor | 1 | 103 |
| C3N-00437-T | Tumor | 1 | 104 |
| C3N-01651-T | Tumor | 1 | 105 |
| C3L-00908-T | Tumor | 1 | 106 |
| C3L-00416-T | Tumor | 1 | 107 |
| C3N-00733-T | Tumor | 1 | 108 |
| C3L-00610-T | Tumor | 1 | 109 |
| C3N-01213-T | Tumor | 1 | 110 |
| C3N-01200-T | Tumor | 1 | 111 |
| C3N-01361-T | Tumor | 1 | 112 |
| C3L-00183-T | Tumor | 1 | 113 |
| C3L-01861-T | Tumor | 1 | 114 |
| C3L-00917-T | Tumor | 1 | 115 |
| C3L-00817-T | Tumor | 1 | 116 |
| C3L-00581-T | Tumor | 1 | 117 |
| C3L-00004-T | Tumor | 1 | 118 |
| C3N-01524-T | Tumor | 1 | 119 |
| C3L-01302-T | Tumor | 1 | 120 |
| C3L-01836-T | Tumor | 1 | 121 |
| C3N-00314-T | Tumor | 1 | 122 |
| C3L-01287-T | Tumor | 1 | 123 |
| C3N-00149-T | Tumor | 1 | 124 |
| C3N-00148-T | Tumor | 1 | 125 |
| C3N-00390-T | Tumor | 1 | 126 |
| C3L-01885-T | Tumor | 1 | 127 |
| C3N-00495-T | Tumor | 1 | 128 |
| C3N-01646-T | Tumor | 1 | 129 |
| C3L-00813-T | Tumor | 1 | 130 |
| C3L-00812-T | Tumor | 1 | 131 |
| C3N-00494-T | Tumor | 1 | 132 |
| C3N-00573-T | Tumor | 1 | 133 |
| C3L-01560-T | Tumor | 1 | 134 |
| C3N-01649-T | Tumor | 1 | 135 |
| C3L-00792-T | Tumor | 1 | 136 |
| C3N-01179-T | Tumor | 1 | 137 |
| C3N-00168-T | Tumor | 1 | 138 |
| C3L-00607-T | Tumor | 1 | 139 |
| C3L-00907-T | Tumor | 1 | 140 |
| C3N-01178-T | Tumor | 1 | 141 |
| C3L-00902-T | Tumor | 1 | 142 |
| C3N-00831-T | Tumor | 1 | 143 |
| C3L-00096-T | Tumor | 1 | 144 |
| C3N-00194-T | Tumor | 1 | 145 |
| C3L-00010-T | Tumor | 1 | 146 |
| C3N-00852-T | Tumor | 1 | 147 |
| C3N-01522-T | Tumor | 1 | 148 |
| C3L-01288-T | Tumor | 1 | 149 |
| C3L-00799-T | Tumor | 1 | 150 |
| C3L-00088-T | Tumor | 1 | 151 |
| C3L-00910-T | Tumor | 1 | 152 |
| C3N-01261-T | Tumor | 1 | 153 |
| C3L-00606-T | Tumor | 1 | 154 |
| C3L-01603-T | Tumor | 1 | 155 |
| C3L-01553-T | Tumor | 1 | 156 |
| C3L-01281-T | Tumor | 1 | 157 |
| C3N-00320-T | Tumor | 1 | 158 |
| C3L-01557-T | Tumor | 1 | 159 |
| C3L-01352-T | Tumor | 1 | 160 |
| C3N-01176-T | Tumor | 1 | 161 |
| C3N-00834-T | Tumor | 1 | 162 |
| C3L-00418-T | Tumor | 1 | 163 |
| C3L-00796-T | Tumor | 1 | 164 |
| C3L-00103-T | Tumor | 1 | 165 |
| C3L-00097-T | Tumor | 1 | 166 |
| C3N-00312-T | Tumor | 1 | 167 |
| C3L-00583-T | Tumor | 1 | 168 |
| C3L-00814-T | Tumor | 1 | 169 |
| C3L-01313-T | Tumor | 1 | 170 |
| C3L-00026-T | Tumor | 1 | 171 |
| C3L-00447-T | Tumor | 1 | 172 |
| C3L-00448-T | Tumor | 1 | 173 |
| C3N-01808-T | Tumor | 1 | 174 |
| C3L-00011-T | Tumor | 1 | 175 |
| C3N-00310-T | Tumor | 1 | 176 |
| C3N-00244-T | Tumor | 1 | 177 |
| C3L-00791-T | Tumor | 1 | 178 |
| C3L-01286-T | Tumor | 1 | 179 |
| C3N-00246-T | Tumor | 1 | 180 |
| C3N-00242-T | Tumor | 1 | 181 |
| C3L-00369-T | Tumor | 1 | 182 |

### 2 Overview

| dataSet | # proteins (genes) | # proteins (genes) [50%] | complex\_ks | gene\_wise\_cor | sample\_wise\_cor | func\_auc |
| --- | --- | --- | --- | --- | --- | --- |
| diann\_diann | 9415 | 6760 | 0.4343524 | 0.3667847 | 0.4262393 | 0.8112997 |
| diaumpire\_diann | 7940 | 6945 | 0.4459407 | 0.3694263 | 0.4263805 | 0.8136781 |
| diaumpiredda\_diann | 8478 | 7160 | 0.4551142 | 0.3594841 | 0.4266302 | 0.7983272 |
| msfraggerdia\_diann | 7973 | 6861 | 0.4388348 | 0.3620459 | 0.4240319 | 0.7961930 |
| msfraggerdiadda\_diann | 8485 | 7063 | 0.4459102 | 0.3595583 | 0.4215754 | 0.8175395 |
| TMT\_MD | 12082 | 9526 | 0.6023089 | 0.4042745 | 0.4595397 | 0.8159911 |

### 3 Descriptive

#### 3.1 Protein/gene identification and quantification

The table below shows the number of identified proteins or genes for each dataset. We take the proteins or genes filtered by 50% missing value as quantified proteins or genes.

|  | dataSet | # proteins (genes) | # proteins (genes) [50%] |
| --- | --- | --- | --- |
| diann\_diann | diann\_diann | 9415 | 6760 |
| diaumpire\_diann | diaumpire\_diann | 7940 | 6945 |
| diaumpiredda\_diann | diaumpiredda\_diann | 8478 | 7160 |
| msfraggerdia\_diann | msfraggerdia\_diann | 7973 | 6861 |
| msfraggerdiadda\_diann | msfraggerdiadda\_diann | 8485 | 7063 |
| TMT\_MD | TMT\_MD | 12082 | 9526 |

Upset chart below showing overlap in proteins or genes identified in each dataset. Numbers of identified proteins or genes shared between different datasets are indicated in the top bar chart and the specific datasets in each set are indicated with solid points below the bar chart. Total identifications for each dataset are indicated on the left as ‘Set size’.

#### 3.2 Protein/gene number distribution

The figures below show the number of proteins or genes identified in each sample. The samples from different batches are coded in different shapes and the samples from different classes are coded in different colors.

diann\_dianndiaumpire\_dianndiaumpiredda\_diannmsfraggerdia\_diannmsfraggerdiadda\_diannTMT\_MD

### 4 Data visualization

#### 4.1 Protein or gene expression distribution

The boxplots show the protein or gene expression distribution across samples. X axis is sample ordered by input order. Y axis is log2 transformed protein or gene expression. The samples from different classes are coded in different colors.

diann\_dianndiaumpire\_dianndiaumpiredda\_diannmsfraggerdia\_diannmsfraggerdiadda\_diannTMT\_MD

The density plots show the protein or gene expression distribution across samples. X axis is log2 transformed protein or gene expression. Y axis is density.

#### 4.2 Batch effect (Heatmap ordered by batches)

In these figures, each column is a sample, each row is also a sample. The color indicates the correlation between samples. The samples are ordered by batches.

diann\_dianndiaumpire\_dianndiaumpiredda\_diannmsfraggerdia\_diannmsfraggerdiadda\_diannTMT\_MD

#### 4.3 Protein or gene coefficient of variation (CV) distribution

diann\_dianndiaumpire\_dianndiaumpiredda\_diannmsfraggerdia\_diannmsfraggerdiadda\_diannTMT\_MD

#### 4.4 Missing value distribution

The missing value distribution can give an overview of the percent of missing values of all proteins or genes in both the QC and experiment samples.

diann\_dianndiaumpire\_dianndiaumpiredda\_diannmsfraggerdia\_diannmsfraggerdiadda\_diannTMT\_MD

#### 4.5 Unsupervised analysis of samples: PCA

diann\_dianndiaumpire\_dianndiaumpiredda\_diannmsfraggerdia\_diannmsfraggerdiadda\_diannTMT\_MD

#### 4.6 Unsupervised analysis of samples: Cluster analysis

diann\_dianndiaumpire\_dianndiaumpiredda\_diannmsfraggerdia\_diannmsfraggerdiadda\_diannTMT\_MD

### 5 Quantitative evaluation

#### 5.1 Correlation between proteins: within vs between protein complexes

The table showing below is a summary of the evaluation. ‘diff’ is Cor(intra) - Cor(inter). ‘ks’ is the statistic value of Kolmogorov-Smirnov test.

| dataSet | InterComplex | IntraComplex | diff | ks |
| --- | --- | --- | --- | --- |
| diann\_diann | 0.027 | 0.259 | 0.232 | 0.434 |
| diaumpire\_diann | 0.042 | 0.288 | 0.246 | 0.446 |
| diaumpiredda\_diann | 0.042 | 0.291 | 0.249 | 0.455 |
| msfraggerdia\_diann | 0.042 | 0.282 | 0.240 | 0.439 |
| msfraggerdiadda\_diann | 0.036 | 0.276 | 0.240 | 0.446 |
| RNA | 0.021 | 0.159 | 0.138 | 0.245 |
| TMT\_MD | 0.025 | 0.454 | 0.429 | 0.602 |

#### 5.2 Correlation between mRNA and protein: gene-wise correlation

| dataSet | n | n5 | n6 | n7 | n8 | median\_cor |
| --- | --- | --- | --- | --- | --- | --- |
| diann\_diann | 6454 | 2067 | 1354 | 701 | 245 | 0.367 |
| diaumpire\_diann | 6635 | 2166 | 1430 | 738 | 253 | 0.369 |
| diaumpiredda\_diann | 6845 | 2138 | 1433 | 738 | 256 | 0.359 |
| msfraggerdia\_diann | 6545 | 2090 | 1411 | 712 | 254 | 0.362 |
| msfraggerdiadda\_diann | 6742 | 2112 | 1400 | 715 | 246 | 0.360 |
| TMT\_MD | 9107 | 3366 | 2194 | 1198 | 459 | 0.404 |

diann\_dianndiaumpire\_dianndiaumpiredda\_diannmsfraggerdia\_diannmsfraggerdiadda\_diannTMT\_MD

#### 5.3 Correlation between mRNA and protein: sample-wise correlation

| dataSet | median\_cor |
| --- | --- |
| diann\_diann | 0.426 |
| diaumpire\_diann | 0.426 |
| diaumpiredda\_diann | 0.427 |
| msfraggerdia\_diann | 0.424 |
| msfraggerdiadda\_diann | 0.422 |
| TMT\_MD | 0.460 |

#### 5.4 Co-expression network based function prediction

In this evaluation, each dataset was used to build co-expression network. For a selected network and a selected function term (such as GO or KEGG), proteins/genes annotated to the term and also included in the network were defined as a positive protein/gene set and other proteins/genes in the network constituted the negative protein/gene set for the term. For a selected function term, we use some of the proteins/genes as the seed protein/gene, then we use random walk algorithm to calculate scores for other proteins/genes. A higher score of a protein/gene represents a closer relationship between the protein/gene and the seed proteins/genes. Finally, for each selected function term, we calculate an AUROC to evaluate the prediction performance.

|  | diann\_diann | diaumpire\_diann | diaumpiredda\_diann | msfraggerdia\_diann | msfraggerdiadda\_diann | RNA | TMT\_MD |
| --- | --- | --- | --- | --- | --- | --- | --- |
| Acute myeloid leukemia | 0.828 | 0.644 | 0.76 | 0.662 | 0.691 | 0.596 | 0.633 |
| Adherens junction | 0.772 | 0.717 | 0.715 | 0.739 | 0.721 | 0.585 | 0.725 |
| Adipocytokine signaling pathway | 0.69 | 0.611 | 0.677 | 0.71 | 0.72 | 0.589 | 0.64 |
| Alanine, aspartate and glutamate metabolism | 0.672 | 0.729 | 0.632 | 0.678 | 0.599 | 0.597 | 0.808 |
| Alzheimers disease | 0.836 | 0.85 | 0.842 | 0.806 | 0.855 | 0.808 | 0.84 |
| Amino sugar and nucleotide sugar metabolism | 0.798 | 0.749 | 0.71 | 0.8 | 0.697 | 0.782 | 0.747 |
| Amoebiasis | 0.732 | 0.759 | 0.733 | 0.708 | 0.749 | 0.737 | 0.726 |
| Amyotrophic lateral sclerosis (ALS) | 0.62 | 0.618 | 0.784 | 0.596 | 0.607 | 0.561 | 0.635 |
| Antigen processing and presentation | 0.945 | 0.935 | 0.907 | 0.903 | 0.904 | 0.903 | 0.942 |
| Apoptosis | 0.75 | 0.571 | 0.681 | 0.599 | 0.61 | 0.633 | 0.681 |
| Arachidonic acid metabolism | 0.558 | 0.69 | 0.756 | 0.639 | 0.663 | 0.554 | 0.609 |
| Arginine and proline metabolism | 0.746 | 0.803 | 0.717 | 0.693 | 0.714 | 0.617 | 0.739 |
| Arrhythmogenic right ventricular cardiomyopathy (ARVC) | 0.807 | 0.779 | 0.82 | 0.817 | 0.754 | 0.711 | 0.817 |
| Ascorbate and aldarate metabolism | 0.857 | 0.897 | 0.793 | 0.875 | 0.85 | 0.734 | 0.88 |
| Axon guidance | 0.567 | 0.7 | 0.624 | 0.693 | 0.702 | 0.581 | 0.739 |
| B cell receptor signaling pathway | 0.72 | 0.769 | 0.72 | 0.654 | 0.717 | 0.682 | 0.78 |
| Bacterial invasion of epithelial cells | 0.744 | 0.754 | 0.725 | 0.675 | 0.648 | 0.641 | 0.807 |
| beta-Alanine metabolism | 0.73 | 0.803 | 0.764 | 0.815 | 0.731 | 0.586 | 0.79 |
| Bile secretion | 0.826 | 0.666 | 0.754 | 0.774 | 0.663 | 0.628 | 0.792 |
| Bladder cancer | 0.752 | 0.697 | 0.677 | 0.629 | 0.645 | 0.634 | 0.669 |
| Butanoate metabolism | 0.839 | 0.865 | 0.812 | 0.779 | 0.808 | 0.774 | 0.872 |
| Calcium signaling pathway | 0.617 | 0.654 | 0.644 | 0.64 | 0.751 | 0.576 | 0.661 |
| Cardiac muscle contraction | 0.888 | 0.916 | 0.88 | 0.878 | 0.866 | 0.783 | 0.771 |
| Cell adhesion molecules (CAMs) | 0.81 | 0.824 | 0.798 | 0.831 | 0.813 | 0.734 | 0.794 |
| Cell cycle | 0.807 | 0.814 | 0.831 | 0.776 | 0.826 | 0.736 | 0.858 |
| Chagas disease (American trypanosomiasis) | 0.714 | 0.649 | 0.627 | 0.526 | 0.72 | 0.605 | 0.712 |
| Chemokine signaling pathway | 0.754 | 0.798 | 0.769 | 0.795 | 0.765 | 0.63 | 0.748 |
| Chronic myeloid leukemia | 0.639 | 0.653 | 0.644 | 0.585 | 0.663 | 0.566 | 0.75 |
| Citrate cycle (TCA cycle) | 0.921 | 0.92 | 0.875 | 0.925 | 0.93 | 0.707 | 0.903 |
| Collecting duct acid secretion | 0.882 | 0.878 | 0.763 | 0.8 | 0.867 | 0.633 | 0.875 |
| Colorectal cancer | 0.655 | 0.662 | 0.641 | 0.647 | 0.745 | 0.687 | 0.695 |
| Complement and coagulation cascades | 0.88 | 0.91 | 0.957 | 0.94 | 0.943 | 0.787 | 0.991 |
| Cysteine and methionine metabolism | 0.655 | 0.772 | 0.74 | 0.695 | 0.814 | 0.592 | 0.767 |
| Dilated cardiomyopathy | 0.771 | 0.765 | 0.74 | 0.626 | 0.682 | 0.697 | 0.773 |
| DNA replication | 0.939 | 0.951 | 0.946 | 0.943 | 0.861 | 0.936 | 0.879 |
| Drug metabolism - cytochrome P450 | 0.739 | 0.677 | 0.768 | 0.842 | 0.697 | 0.739 | 0.816 |
| Drug metabolism - other enzymes | 0.673 | 0.709 | 0.648 | 0.72 | 0.677 | 0.601 | 0.614 |
| ECM-receptor interaction | 0.847 | 0.888 | 0.897 | 0.825 | 0.903 | 0.752 | 0.857 |
| Endocytosis | 0.727 | 0.682 | 0.685 | 0.675 | 0.693 | 0.62 | 0.775 |
| Endometrial cancer | 0.781 | 0.784 | 0.812 | 0.696 | 0.652 | 0.598 | 0.711 |
| Epithelial cell signaling in Helicobacter pylori infection | 0.697 | 0.751 | 0.688 | 0.748 | 0.715 | 0.617 | 0.721 |
| ErbB signaling pathway | 0.591 | 0.711 | 0.61 | 0.595 | 0.639 | 0.563 | 0.638 |
| Fatty acid metabolism | 0.916 | 0.852 | 0.867 | 0.843 | 0.849 | 0.79 | 0.839 |
| Fc epsilon RI signaling pathway | 0.56 | 0.708 | 0.626 | 0.657 | 0.675 | 0.541 | 0.742 |
| Fc gamma R-mediated phagocytosis | 0.786 | 0.787 | 0.822 | 0.715 | 0.774 | 0.742 | 0.724 |
| Focal adhesion | 0.784 | 0.76 | 0.792 | 0.737 | 0.786 | 0.71 | 0.782 |
| Fructose and mannose metabolism | 0.829 | 0.654 | 0.741 | 0.741 | 0.843 | 0.76 | 0.797 |
| Galactose metabolism | 0.723 | 0.677 | 0.73 | 0.744 | 0.818 | 0.701 | 0.801 |
| Gap junction | 0.635 | 0.715 | 0.672 | 0.674 | 0.687 | 0.59 | 0.645 |
| Gastric acid secretion | 0.671 | 0.581 | 0.604 | 0.558 | 0.669 | 0.601 | 0.687 |
| Glutathione metabolism | 0.556 | 0.601 | 0.628 | 0.615 | 0.681 | 0.59 | 0.565 |
| Glycerolipid metabolism | 0.788 | 0.687 | 0.737 | 0.757 | 0.81 | 0.562 | 0.777 |
| Glycerophospholipid metabolism | 0.71 | 0.612 | 0.725 | 0.62 | 0.635 | 0.587 | 0.716 |
| Glycine, serine and threonine metabolism | 0.769 | 0.777 | 0.659 | 0.718 | 0.648 | 0.792 | 0.795 |
| Glycolysis / Gluconeogenesis | 0.786 | 0.689 | 0.769 | 0.729 | 0.792 | 0.705 | 0.779 |
| Glyoxylate and dicarboxylate metabolism | 0.688 | 0.734 | 0.637 | 0.679 | 0.731 | 0.735 | 0.794 |
| GnRH signaling pathway | 0.614 | 0.603 | 0.695 | 0.585 | 0.602 | 0.543 | 0.624 |
| Hematopoietic cell lineage | 0.71 | 0.634 | 0.669 | 0.733 | 0.737 | 0.681 | 0.816 |
| Hepatitis C | 0.716 | 0.782 | 0.783 | 0.746 | 0.759 | 0.59 | 0.745 |
| Histidine metabolism | 0.801 | 0.769 | 0.836 | 0.839 | 0.801 | 0.818 | 0.77 |
| Huntingtons disease | 0.859 | 0.871 | 0.839 | 0.792 | 0.845 | 0.763 | 0.86 |
| Hypertrophic cardiomyopathy (HCM) | 0.778 | 0.82 | 0.69 | 0.734 | 0.73 | 0.683 | 0.788 |
| Inositol phosphate metabolism | 0.656 | 0.693 | 0.654 | 0.591 | 0.673 | 0.703 | 0.742 |
| Insulin signaling pathway | 0.625 | 0.702 | 0.663 | 0.639 | 0.637 | 0.57 | 0.685 |
| Leishmaniasis | 0.83 | 0.855 | 0.849 | 0.798 | 0.778 | 0.802 | 0.884 |
| Leukocyte transendothelial migration | 0.804 | 0.791 | 0.763 | 0.743 | 0.797 | 0.673 | 0.747 |
| Long-term depression | 0.694 | 0.7 | 0.702 | 0.693 | 0.695 | 0.507 | 0.728 |
| Long-term potentiation | 0.709 | 0.664 | 0.69 | 0.631 | 0.633 | 0.605 | 0.783 |
| Lysine degradation | 0.911 | 0.876 | 0.852 | 0.884 | 0.857 | 0.664 | 0.79 |
| Lysosome | 0.838 | 0.886 | 0.888 | 0.846 | 0.829 | 0.773 | 0.884 |
| Malaria | 0.828 | 0.752 | 0.778 | 0.77 | 0.848 | 0.725 | 0.845 |
| MAPK signaling pathway | 0.576 | 0.608 | 0.588 | 0.646 | 0.58 | 0.648 | 0.655 |
| Melanogenesis | 0.632 | 0.634 | 0.668 | 0.661 | 0.592 | 0.601 | 0.728 |
| Metabolic pathways | 0.73 | 0.733 | 0.752 | 0.761 | 0.745 | 0.664 | 0.785 |
| Metabolism of xenobiotics by cytochrome P450 | 0.734 | 0.743 | 0.85 | 0.846 | 0.752 | 0.713 | 0.743 |
| mRNA surveillance pathway | 0.781 | 0.787 | 0.802 | 0.829 | 0.793 | 0.719 | 0.843 |
| mTOR signaling pathway | 0.655 | 0.794 | 0.745 | 0.658 | 0.737 | 0.615 | 0.681 |
| N-Glycan biosynthesis | 0.924 | 0.96 | 0.925 | 0.817 | 0.801 | 0.684 | 0.938 |
| Natural killer cell mediated cytotoxicity | 0.677 | 0.794 | 0.734 | 0.667 | 0.697 | 0.695 | 0.735 |
| Neurotrophin signaling pathway | 0.733 | 0.687 | 0.628 | 0.604 | 0.603 | 0.573 | 0.683 |
| NOD-like receptor signaling pathway | 0.67 | 0.625 | 0.712 | 0.55 | 0.644 | 0.637 | 0.781 |
| Oocyte meiosis | 0.802 | 0.754 | 0.78 | 0.69 | 0.733 | 0.569 | 0.762 |
| Osteoclast differentiation | 0.677 | 0.772 | 0.689 | 0.678 | 0.643 | 0.641 | 0.805 |
| Oxidative phosphorylation | 0.922 | 0.961 | 0.941 | 0.926 | 0.926 | 0.863 | 0.933 |
| Pancreatic cancer | 0.643 | 0.646 | 0.677 | 0.564 | 0.671 | 0.564 | 0.694 |
| Pancreatic secretion | 0.652 | 0.626 | 0.625 | 0.655 | 0.684 | 0.581 | 0.74 |
| Parkinsons disease | 0.841 | 0.846 | 0.847 | 0.803 | 0.823 | 0.805 | 0.874 |
| Pathogenic Escherichia coli infection | 0.69 | 0.763 | 0.753 | 0.765 | 0.765 | 0.629 | 0.69 |
| Pathways in cancer | 0.716 | 0.665 | 0.632 | 0.612 | 0.669 | 0.539 | 0.637 |
| Pentose and glucuronate interconversions | 0.894 | 0.751 | 0.861 | 0.807 | 0.862 | 0.623 | 0.832 |
| Pentose phosphate pathway | 0.829 | 0.74 | 0.703 | 0.744 | 0.83 | 0.623 | 0.657 |
| Peroxisome | 0.763 | 0.763 | 0.758 | 0.706 | 0.829 | 0.768 | 0.842 |
| Phagosome | 0.823 | 0.844 | 0.829 | 0.854 | 0.815 | 0.697 | 0.847 |
| Phosphatidylinositol signaling system | 0.682 | 0.66 | 0.646 | 0.634 | 0.654 | 0.601 | 0.552 |
| Porphyrin and chlorophyll metabolism | 0.689 | 0.619 | 0.584 | 0.805 | 0.669 | 0.598 | 0.652 |
| PPAR signaling pathway | 0.722 | 0.663 | 0.728 | 0.723 | 0.74 | 0.704 | 0.689 |
| Prion diseases | 0.822 | 0.694 | 0.77 | 0.731 | 0.701 | 0.659 | 0.745 |
| Progesterone-mediated oocyte maturation | 0.674 | 0.816 | 0.669 | 0.699 | 0.646 | 0.611 | 0.702 |
| Propanoate metabolism | 0.87 | 0.81 | 0.884 | 0.833 | 0.889 | 0.747 | 0.916 |
| Prostate cancer | 0.734 | 0.649 | 0.6 | 0.682 | 0.54 | 0.574 | 0.673 |
| Proteasome | 0.993 | 0.993 | 0.993 | 0.987 | 0.998 | 0.893 | 0.984 |
| Protein digestion and absorption | 0.865 | 0.811 | 0.816 | 0.837 | 0.868 | 0.662 | 0.87 |
| Protein export | 0.889 | 0.99 | 0.976 | 0.991 | 0.898 | 0.812 | 0.961 |
| Protein processing in endoplasmic reticulum | 0.811 | 0.815 | 0.835 | 0.807 | 0.802 | 0.756 | 0.791 |
| Purine metabolism | 0.6 | 0.655 | 0.645 | 0.645 | 0.618 | 0.594 | 0.622 |
| Pyrimidine metabolism | 0.53 | 0.557 | 0.58 | 0.585 | 0.618 | 0.625 | 0.6 |
| Pyruvate metabolism | 0.831 | 0.752 | 0.705 | 0.758 | 0.778 | 0.602 | 0.82 |
| Regulation of actin cytoskeleton | 0.728 | 0.751 | 0.725 | 0.751 | 0.762 | 0.706 | 0.732 |
| Renal cell carcinoma | 0.693 | 0.667 | 0.61 | 0.597 | 0.632 | 0.585 | 0.628 |
| Retinol metabolism | 0.824 | 0.83 | 0.836 | 0.831 | 0.804 | 0.706 | 0.922 |
| Rheumatoid arthritis | 0.858 | 0.913 | 0.875 | 0.853 | 0.865 | 0.708 | 0.886 |
| Ribosome | 0.964 | 0.945 | 0.971 | 0.96 | 0.956 | 0.906 | 0.985 |
| Ribosome biogenesis in eukaryotes | 0.771 | 0.812 | 0.804 | 0.828 | 0.814 | 0.689 | 0.867 |
| RIG-I-like receptor signaling pathway | 0.773 | 0.742 | 0.666 | 0.651 | 0.682 | 0.751 | 0.84 |
| RNA degradation | 0.737 | 0.662 | 0.767 | 0.758 | 0.763 | 0.695 | 0.641 |
| RNA transport | 0.803 | 0.807 | 0.851 | 0.789 | 0.812 | 0.643 | 0.803 |
| Salivary secretion | 0.628 | 0.57 | 0.691 | 0.655 | 0.63 | 0.663 | 0.703 |
| Shigellosis | 0.741 | 0.8 | 0.78 | 0.76 | 0.732 | 0.707 | 0.84 |
| Small cell lung cancer | 0.757 | 0.838 | 0.741 | 0.711 | 0.827 | 0.727 | 0.73 |
| SNARE interactions in vesicular transport | 0.641 | 0.798 | 0.845 | 0.766 | 0.74 | 0.682 | 0.786 |
| Spliceosome | 0.92 | 0.905 | 0.893 | 0.916 | 0.897 | 0.782 | 0.894 |
| Staphylococcus aureus infection | 0.974 | 0.971 | 0.981 | 0.971 | 0.98 | 0.862 | 0.964 |
| Starch and sucrose metabolism | 0.757 | 0.697 | 0.751 | 0.756 | 0.9 | 0.598 | 0.857 |
| Systemic lupus erythematosus | 0.841 | 0.914 | 0.905 | 0.87 | 0.891 | 0.812 | 0.9 |
| T cell receptor signaling pathway | 0.659 | 0.746 | 0.706 | 0.555 | 0.667 | 0.687 | 0.734 |
| TGF-beta signaling pathway | 0.788 | 0.83 | 0.7 | 0.824 | 0.78 | 0.585 | 0.745 |
| Tight junction | 0.708 | 0.749 | 0.752 | 0.702 | 0.714 | 0.602 | 0.749 |
| Toll-like receptor signaling pathway | 0.658 | 0.682 | 0.552 | 0.658 | 0.633 | 0.604 | 0.742 |
| Toxoplasmosis | 0.653 | 0.749 | 0.756 | 0.684 | 0.743 | 0.592 | 0.67 |
| Tryptophan metabolism | 0.759 | 0.693 | 0.806 | 0.75 | 0.693 | 0.715 | 0.852 |
| Tyrosine metabolism | 0.602 | 0.657 | 0.669 | 0.675 | 0.66 | 0.77 | 0.695 |
| Ubiquitin mediated proteolysis | 0.714 | 0.75 | 0.793 | 0.688 | 0.701 | 0.679 | 0.751 |
| Valine, leucine and isoleucine degradation | 0.889 | 0.853 | 0.87 | 0.839 | 0.853 | 0.799 | 0.88 |
| Vascular smooth muscle contraction | 0.764 | 0.726 | 0.76 | 0.688 | 0.702 | 0.632 | 0.751 |
| Vasopressin-regulated water reabsorption | 0.824 | 0.792 | 0.77 | 0.776 | 0.732 | 0.637 | 0.76 |
| VEGF signaling pathway | 0.671 | 0.736 | 0.735 | 0.56 | 0.687 | 0.671 | 0.676 |
| Vibrio cholerae infection | 0.725 | 0.815 | 0.749 | 0.783 | 0.678 | 0.66 | 0.678 |
| Viral myocarditis | 0.909 | 0.85 | 0.879 | 0.878 | 0.83 | 0.826 | 0.747 |
| Wnt signaling pathway | 0.627 | 0.722 | 0.525 | 0.501 | 0.716 | 0.575 | 0.612 |

diann\_dianndiaumpire\_dianndiaumpiredda\_diannmsfraggerdia\_diannmsfraggerdiadda\_diannTMT\_MD
