## Supplementary Figures for "One-stop analysis of DIA proteomics data using MSFragger-DIA and FragPipe computational platform"

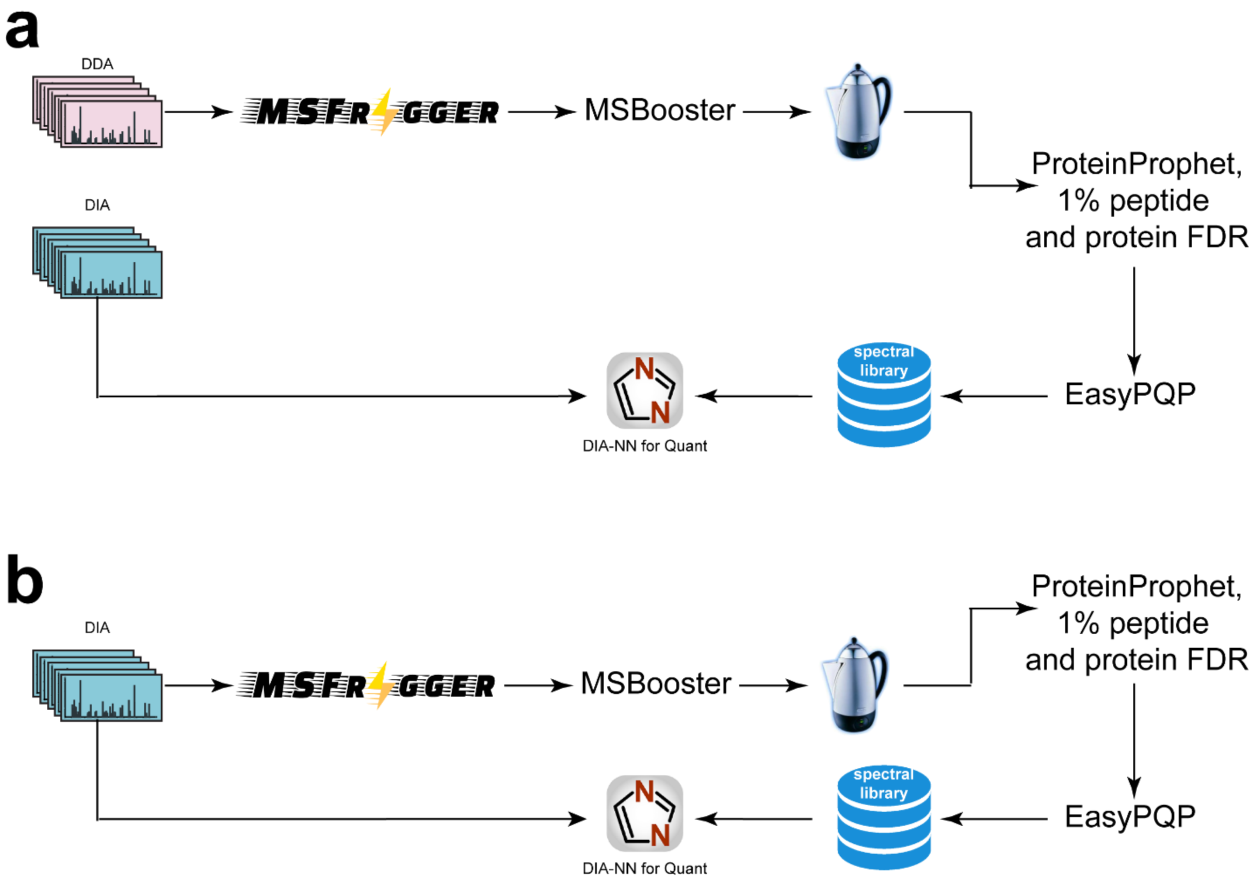


**Supplementary Figure 1.** DIA data analysis pipelines covering library-based, library-free, and hybrid approaches. **(a) Library-based analysis.** MSFragger in DDA mode is used to search the DDA data, followed by MSBooster, Percolator, ProteinProphet, FDR filtering, and spectral library generation with EasyPQP. DIA-NN is used to quantify DIA data using the spectral library from the DDA data. **(b) Library-free analysis.** MSFragger-DIA is used to search the DIA data to build a spectral library. Then, the spectral library is used to quantify the DIA data.

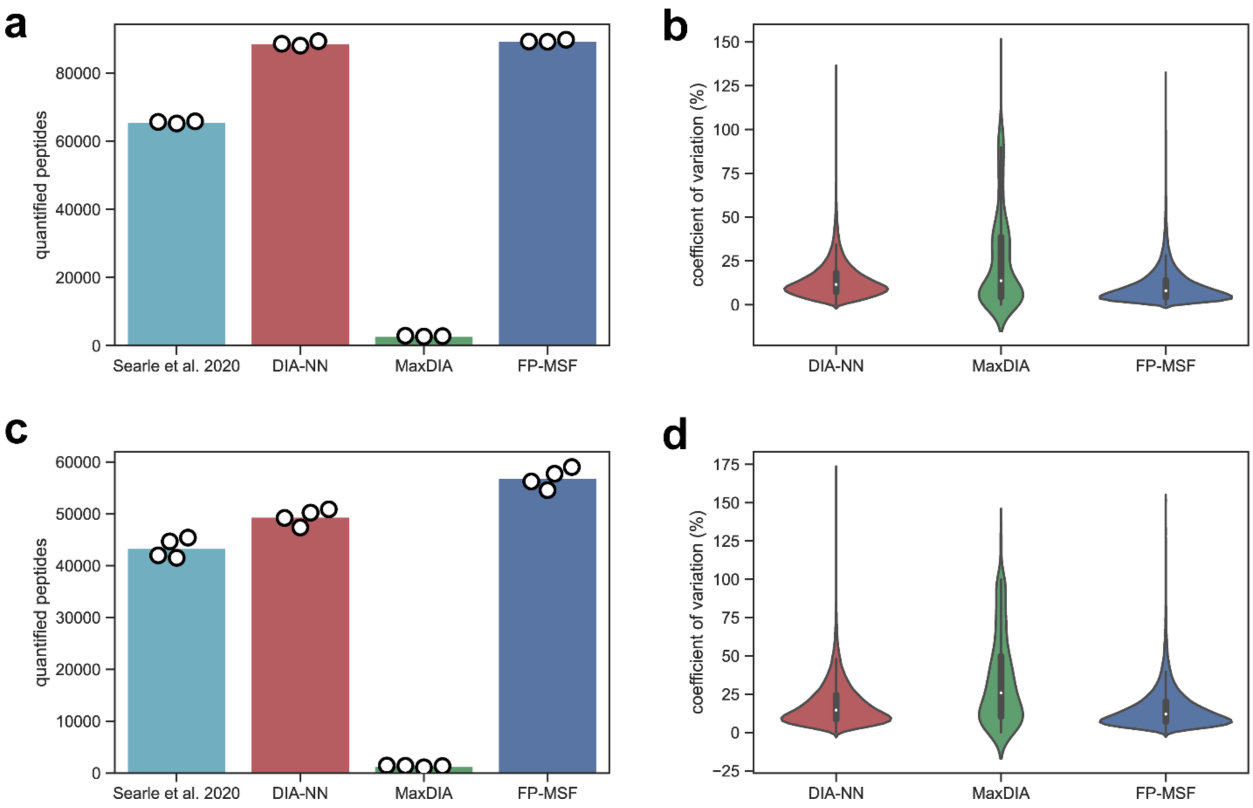


**Supplementary Figure 2.** Quantified peptides and coefficient of variation (CV) from the **2018-HeLa** and the **2020-Yeast** datasets using **DIA-NN *in-silico* library-based**, **MaxDIA**, and **FP-MSF** pipelines. The results from the publication were also used as benchmarks. **(a)** Bar plots of the quantified peptides from **2018-HeLa**. The bar height is the average number of three replicates. The white dots indicate the numbers from individual replicates. **(b)** Violin plots of peptide CVs from **2018-HeLa**. **(c)** Bar plots of the quantified peptides from **2020-Yeast**. **(d)** Violin plots of peptide CVs from **2020-Yeast**.

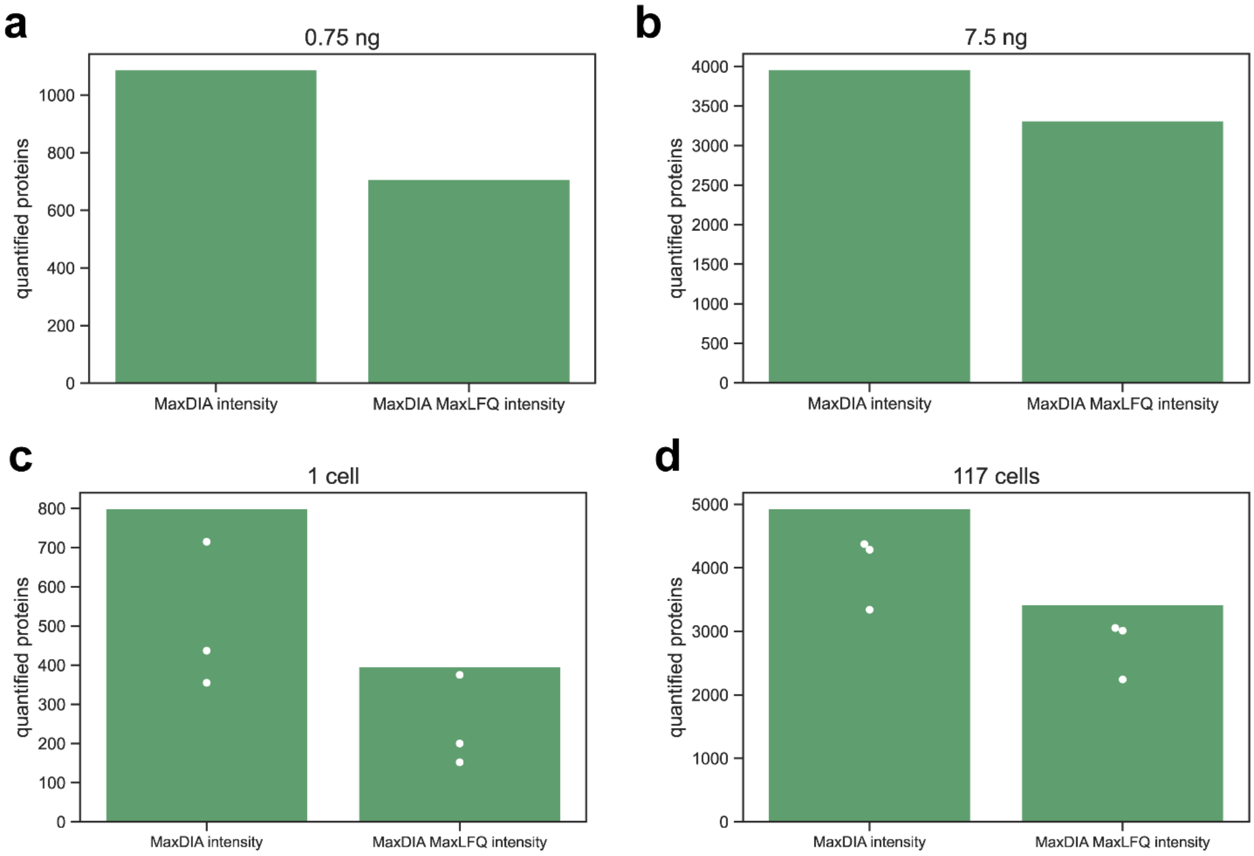


**Supplementary Figure 3**. Quantified proteins from the **low-input-cell** and the **single-cell** datasets using MaxDIA. The original intensity and the MaxLFQ intensity were used to count the quantified proteins. **(a-b)** Bar plots from analyzing the **low-input-cell** dataset. Proteins with zero intensities are discarded. **(c-d)** Bar plots from analyzing the **single-cell** datasets. The bar height is the total number of proteins from the three replicates. The white dots are the numbers of proteins quantified in each replicate.
